# Stepwise DNA unwinding gates TnpB genome-editing activity

**DOI:** 10.64898/2026.01.09.698545

**Authors:** Zehan Zhou, Iren Saffarian-Deemyad, Honglue Shi, Trevor Weiss, Muhammad Moez Ur-Rehman, Kamakshi Vohra, Petr Skopintsev, Peter H. Yoon, Marena I. Trinidad, Conner J. Langeberg, Maris Kamalu, Jasmine Amerasekera, Yuxing Zhou, Erin E. Doherty, Kevin D.P. Aris, Noor Al-Sayyad, Brittney W. Thornton, Rachel F. Weissman, Kevin M. Wasko, Isabel Esain-Garcia, Evan C. DeTurk, David F. Savage, Steven E. Jacobsen, Zev Bryant, Jennifer A. Doudna

**Affiliations:** Innovative Genomics Institute, University of California, Berkeley, CA, USA, 94720; Department of Molecular and Cell Biology, University of California, Berkeley, CA, USA, 94720; Department of Physics, Stanford University, Stanford, CA, USA, 94305; Howard Hughes Medical Institute, University of California, Berkeley, CA, USA, 94720; California Institute for Quantitative Biosciences (QB3), University of California, Berkeley, Berkeley, CA, USA, 94720; Department of Molecular, Cell and Developmental Biology, University of California at Los Angeles, Los Angeles, CA, USA, 90095; University of California, Berkeley-University of California, San Francisco Graduate Program in Bioengineering, University of California, Berkeley, Berkeley, CA, USA, 94720; Department of Bioengineering, Stanford University, Stanford, CA, USA, 94305; Biophysics Program, Stanford University, Stanford, CA, USA, 94305; Howard Hughes Medical Institute, University of California at Los Angeles, Los Angeles, CA, USA, 90095; Department of Structural Biology, Stanford University Medical Center, Stanford, CA, USA, 94305; Li Ka Shing Center for Genomic Engineering, University of California, Berkeley, Berkeley, CA, USA, 94720; Department of Chemistry, University of California, Berkeley, Berkeley, CA, USA, 94720; Molecular Biophysics and Integrated Bioimaging Division, Lawrence Berkeley National Laboratory, Berkeley, CA, USA, 94720; Gladstone Institute of Data Science and Biotechnology, San Francisco, CA, USA, 94158; Gladstone-UCSF Institute of Genomic Immunology, San Francisco, CA, USA, 94158

**Keywords:** Genome editing, TnpB, CRISPR-Cas, RNA-guided nuclease, DNA unwinding, *R*-loop formation, Single-molecule biophysics, Plant genome editing, Protein engineering, Energy landscape

## Abstract

TnpB is a compact RNA-guided endonuclease and evolutionary ancestor of CRISPR-Cas12 that offers a promising platform for genome engineering. However, the genome-editing activity of TnpBs remains limited and its underlying determinants are poorly understood. Here, we used biochemical and single-molecule assays to examine the DNA-unwinding mechanism of *Youngiibacter multivorans* TnpB (Ymu1 TnpB). DNA unwinding proceeds through a discrete, long-lived partially unwound intermediate state before reaching a fully unwound open state. The open state forms inefficiently and collapses readily in the absence of negative supercoiling. An optimized variant, Ymu1-WFR, stabilizes formation of both the intermediate and open states, resulting in enhanced DNA cleavage *in vitro* and increased genome editing in plants. These findings identify the physical basis for the observed minimal activities of natural TnpBs, revealing how stabilizing specific unwinding states enables efficient DNA targeting.

## INTRODUCTION

TnpB proteins are bacterial RNA-guided homing endonucleases encoded by IS605 and IS607 transposable elements and proposed to be the evolutionary ancestors of CRISPR-Cas12 enzymes^1–4^. Recent cryo-EM structures revealed that, like Cas12, TnpB adopts a bi-lobed architecture with conserved wedge (WED), recognition (REC) and RuvC domains, along with a lid subdomain that is hypothesized to gate nuclease activity^2,5^. TnpB activity depends on a non-coding RNA, called the reRNA (also termed ωRNA) that contains a scaffold for protein binding and a ∼16-nucleotide (16-nt) programmable guide region^1^. Guided by the reRNA, TnpB recognizes complementary DNA sequences adjacent to a transposon-adjacent motif (TAM), enabling RNA-DNA base pairing to form an *R*-loop structure prior to cleavage^1,2,5^.

The compact size of TnpBs makes them attractive for genome editing in systems where delivery constraints require smaller editing enzymes^6,7^, particularly for plant editing applications for which enzyme delivery using size-constrained RNA viruses is desirable^8^. However, proof-of-principle studies in plants have thus far reported only modest TnpB-catalyzed editing efficiencies, including in *Arabidopsis thaliana* and other plant systems^8–12^. To develop more effective variants, both directed evolution^13^ and deep mutational scanning (DMS)^12^ were used to identify numerous activity-enhancing mutations that cluster in the nucleic acid binding and catalytic regions of TnpB structures. Nonetheless, the mechanistic basis for the low activity of natural TnpBs, and how engineered variants can overcome this limitation, remain poorly understood.

Here, we address this gap by identifying molecular determinants that limit TnpB efficiency. Combining bulk biochemical assays and single-molecule gold rotor-bead tracking (AuRBT), we found that *Youngiibacter multivorans* TnpB (Ymu1 TnpB) unwinds DNA through a stepwise pathway comprising two distinct transitions: an initial closed-to-intermediate transition and a subsequent intermediate-to-open completion, forming a fully unwound state. Mutational analysis revealed that these transitions are modulated by the WED domain and an α-helix flanking the lid subdomain (hereafter, lid-adjacent helix): a WED-domain mutation promoted early transition, whereas mutations in the lid-adjacent helix stabilize more extensive unwinding and favor progression to the fully unwound state. Together, these perturbations differentially stabilize specific unwound conformations, enabling enhanced DNA cleavage *in vitro* and editing efficiency in plants. These findings establish a stepwise DNA unwinding model for the conformational control in TnpB and provide a mechanistic framework for rational engineering hypercompact, high-performance RNA-guided nucleases.

## RESULTS

### Reconstituted Ymu1 TnpB for mechanistic studies

TnpB enzymes are found in the IS605 and IS607 transposon families and share a deep evolutionary relationship with type V CRISPR-Cas12 systems, with Cas12f proteins representing the closest related clade (Fig. 1A). Ymu1 TnpB stands out for its unusually compact size (382 aa), less than one third that of LbCas12a (Fig. 1B). Among the TnpB orthologs tested, Ymu1 TnpB expressed robustly in *E. coli* and formed stable, homogeneous ribonucleoprotein (RNP) complexes with its reRNA (Fig. S1A-C), comprising a 127-nt scaffold and a 16-nt guide sequence (Fig. 1C and Fig. S1D), whereas other orthologs exhibited lower yields and reduced stability (Fig. S1B-C). Purified Ymu1 RNPs displayed reproducible cleavage of double-stranded DNA (dsDNA) substrates *in vitro* using a previously validated target sequence (Target 1: 5′-TCTTCTGGATTGTTGT-3′) from biochemical and single-molecule studies of Cas12 enzymes^14,15^ (Fig. S1E-F), enabling mechanistic *in vitro* studies of the reaction pathway (Fig. 1D). Previous studies have demonstrated genome-editing activities of Ymu1 TnpB across bacterial, mammalian, and plant systems^6,8^. To establish a quantitative cellular baseline for mechanistic analysis, we evaluated Ymu1 TnpB activity across multiple endogenous genomic targets in *Arabidopsis* protoplasts using a previously established single-transcript expression system (TnpB-reRNA-HDV) (Fig. 1E; Methods)^8^. Across all fourteen tested loci, editing efficiencies remained below 10% (Fig. 1F), consistent with prior reports^8^. Together, these results establish Ymu1 TnpB as a compact and biochemically tractable RNA-guided nuclease with constrained genome-editing efficiency in plants.

**Fig. 1.**
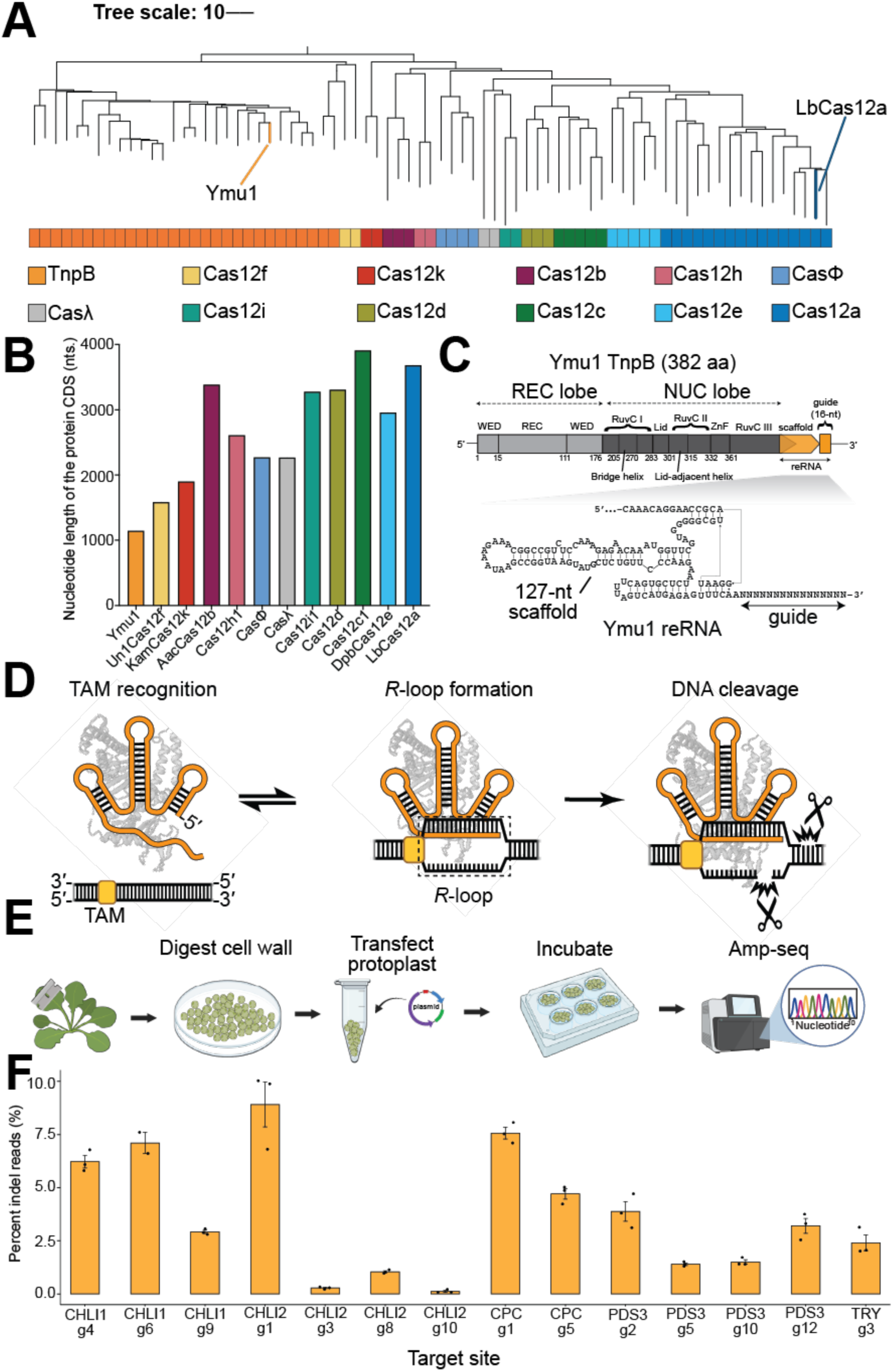
A reconstituted Ymu1 TnpB system for mechanistic and genome-editing studies. **(A)** Phylogenetic placement of Ymu1 TnpB. Dendrogram of selected TnpB orthologs (N = 29) and Cas12-family nucleases (N = 46) showing their evolutionary relationships. Ymu1 TnpB and LbCas12a are highlighted. **(B)** Size of Ymu1 TnpB relative to other Cas12 enzymes. Bar plot comparing coding-sequence (CDS) length (nucleotides) of Ymu1 TnpB with representative Cas12 subtypes. **(C)** Genomic architecture and RNA-protein organization of Ymu1 TnpB. Domain map showing recognition lobe (WED and REC domains) and catalytic lobe (RuvC, lid, ZnF domains), with the bridge helix positioned within RuvC I and the lid-adjacent helix positioned within RuvC II. The reRNA locus lies immediately downstream of the TnpB gene; the guide region is 16-nt. The secondary structure model of the 127-nt reRNA is shown below. **(D)** Cartoon of the TnpB reaction pathway. TAM binding, *R*-loop formation, and DNA cleavage are illustrated schematically, with TAM and *R*-loop positions indicated. **(E)** Schematic of the *Arabidopsis* protoplast editing assay. Workflow consists of leaf digestion, protoplast isolation, plasmid transfection, incubation, genomic DNA extraction, amplicon library preparation, and next-generation sequencing. **(F)** Genome-editing activity of Ymu1 TnpB across endogenous *Arabidopsis* loci (Table S1). All plant experiments were conducted using a 16-nt guide length, which produced the highest indel frequency in a guide length screen ranging from 14 to 20 nts in Arabidopsis protoplasts (Fig. S1G). Target sites are labeled by gene name and guide number (e.g., *CHLI1* g4), where the guide number designates the programmable 16-nt region of the reRNA directed to that site. Percentage of insertion-deletion (indel) reads (y-axis) obtained from amplicon sequencing at multiple genomic sites (x-axis). Bars represent mean ± SEM (n = 3 biological replicates).

### Ymu1 TnpB mutagenesis identifies determinants of enhanced genome editing

The modest editing efficiency of Ymu1 TnpB prompted us to explore whether amino-acid substitutions could improve its performance. Candidate residues were selected based on their predicted proximity to nucleic acids in an AF3 structural model of the Ymu1 ternary complex, guided in part by positions identified as mutationally sensitive in DMS of the related Dra2 TnpB^12^ (Methods). We targeted three structural regions: (i) the WED domain, where H4 is positioned adjacent to the first RNA-DNA base pair; (ii) the bridge helix (K229 and R230), which may couple to distal *R*-loop formation; and (iii) a cluster of residues spanning the lid and lid-adjacent helix (N282-V305), a region predicted to influence conformational dynamics during *R*-loop progression and that includes mutationally sensitive positions in Dra2 TnpB (Methods). To screen amino-acid substitutions, we adopted a short reRNA scaffold engineered through SHAPE-guided truncation and stabilization of the 127-nt reRNA (Methods) that preserves the core secondary structure, as confirmed by SHAPE-MaP analysis (Fig. S2A-B). Both reRNA scaffolds show comparable editing efficiency in protoplast assays (Fig. S2C). We examined a total of 54 variants (both single and combinatorial substitutions) in *Arabidopsis* protoplasts targeting the *PDS3* g2 site (also termed Target 2: 5′-AAGGCAAATTCGCCGC-3′) (Fig. 2A). Two mutations, a WED-domain mutation H4W and a lid-adjacent-helix mutation L304F, increased editing activity relative to wild-type (WT) Ymu1 (Fig. 2A). The nearby mutation V305R, which lies in the same lid-adjacent helix, performed similarly to WT in this assay and was retained for further analysis. Analogous substitutions at all three positions have precedent in prior mutagenesis of Dra2 TnpB: hydrophobic aromatic substitutions are favored at the position homologous to H4, the native Phe of Dra2 TnpB is preferred over Leu at the position homologous to L304, and mutation to Arg is favored at the position homologous to V305^12^. Combining these substitutions yielded a triple variant (H4W-L304F-V305R, hereafter Ymu1-WFR) that consistently outperformed WT and the individual single mutants in the protoplast genome-editing assay (Fig. 2A).

**Fig. 2.**
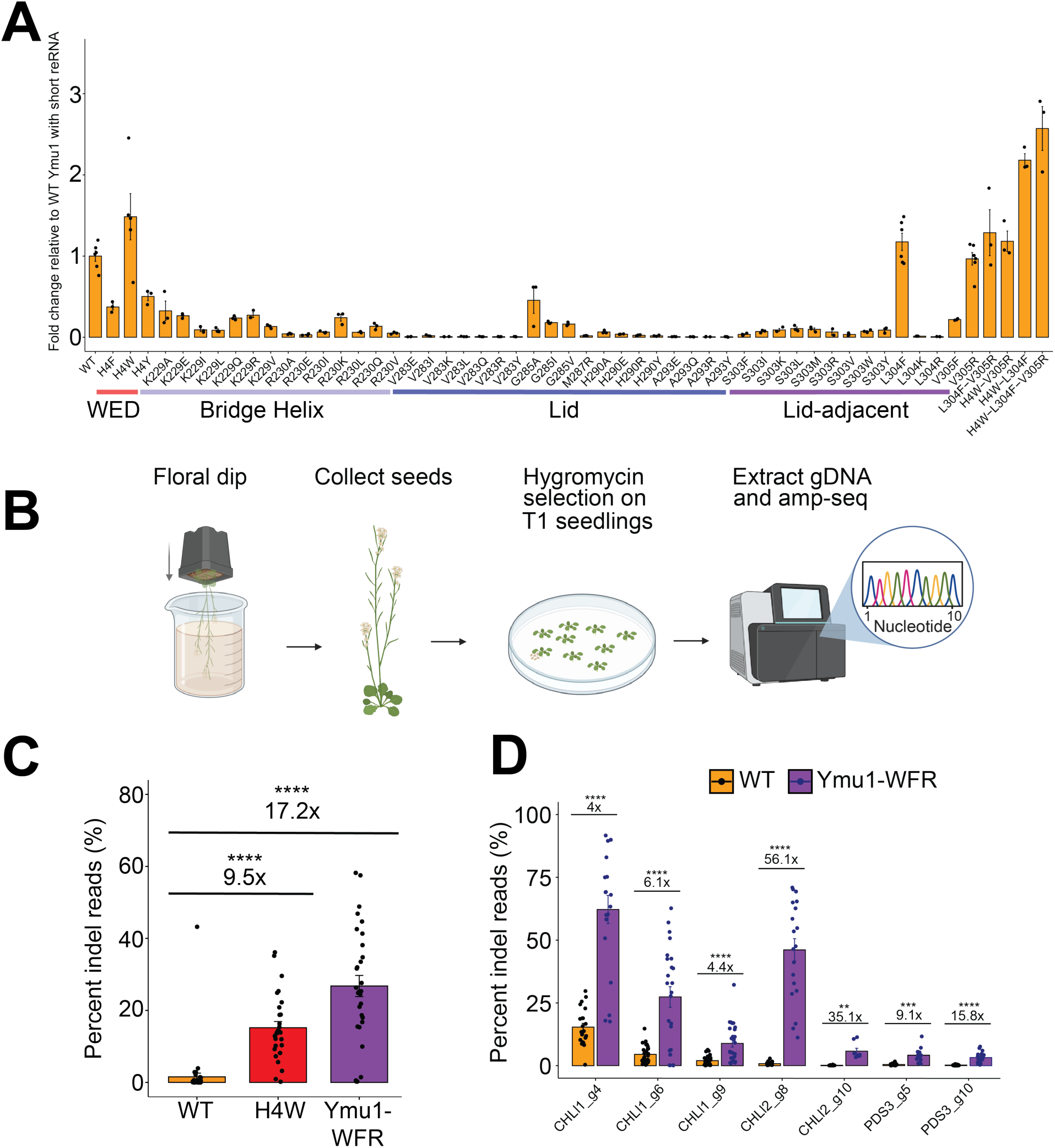
Targeted mutagenesis of Ymu1 TnpB identifies structural determinants that enhance genome-editing activity. **(A)** Mutational screen of Ymu1 TnpB in *Arabidopsis* protoplasts using the short reRNA (Fig. S2B). Bar plot showing fold change in indel percentage relative to WT Ymu1 TnpB with the short reRNA (y-axis) for amino-acid substitutions (x-axis) at the endogenous *PDS3* g2 site (Target 2: 5′-AAGGCAAATTCGCCGC-3′). Structural domains are indicated beneath residue labels. Combinations of selected activating mutations are shown on the right side of the plot, including the triple variant H4W-L304F-V305R (“Ymu1-WFR”). Bars represent mean ± SEM (n = 3 or 6 biological replicates). **(B)** Workflow for T1 whole-plant genome-editing assays. Schematic showing Agrobacterium floral dip, T1 seed harvest, hygromycin selection, genomic DNA isolation, and amplicon sequencing. **(C)** Editing efficiencies of WT Ymu1, H4W, and Ymu1-WFR with the 127-nt reRNA in T1 *Arabidopsis* plants at the *PDS3* g2 site. Bar plot shows percentage of indel reads (mean ± SEM). Fold change and p-values relative to WT are indicated above bars. WT Ymu1 editing data were previously published only for this locus^8^ and are replotted here for direct comparison with mutant variants generated in this study; all experiments were performed using identical methods. **(D)** Editing efficiencies of WT Ymu1 and Ymu1-WFR with the 127-nt reRNA across multiple endogenous genomic loci in T1 plants. Bar plot shows percentage of indel reads (mean ± SEM). Fold change and p-values relative to WT are indicated above each site. All the data in **(D)** were generated in this study.

To test whether these improvements extend to whole plants (Fig. 2B; Methods), we first evaluated H4W and Ymu1-WFR at the *PDS3* g2 site in T1 transgenic *Arabidopsis* plants. Because H4W and Ymu1-WFR showed higher editing efficiency with the 127-nt reRNA than with the short reRNA (Fig. S2D), the 127-nt reRNA was used for all subsequent plant genome-editing, biochemical, and biophysical experiments in this study. Relative to previously reported WT Ymu1 data obtained under identical conditions^8^, H4W and Ymu1-WFR exhibited ∼10-fold and ∼17-fold higher editing efficiency, respectively (Fig. 2C). To confirm the enhanced editing activity was not specific to the *PDS3* g2 site, we compared editing between Ymu1-WFR and WT across seven additional targets and observed significantly increased editing at all sites, with improvement of up to 56-fold (Fig. 2D). Independently, Ymu1-WFR has been shown to achieve up to 9.8-fold higher editing efficiency than WT Ymu1 via viral delivery in *Arabidopsis*^16^. These results identify mutations in the WED and lid-adjacent helix that enhance Ymu1 activity in plants. Consistent with the location of the mutations outside the TAM-interacting region (Fig. 3A), WT Ymu1 and Ymu1-WFR exhibited indistinguishable TAM preferences (5′-TTGAT-3′; Fig. S2E).

**Fig. 3.**
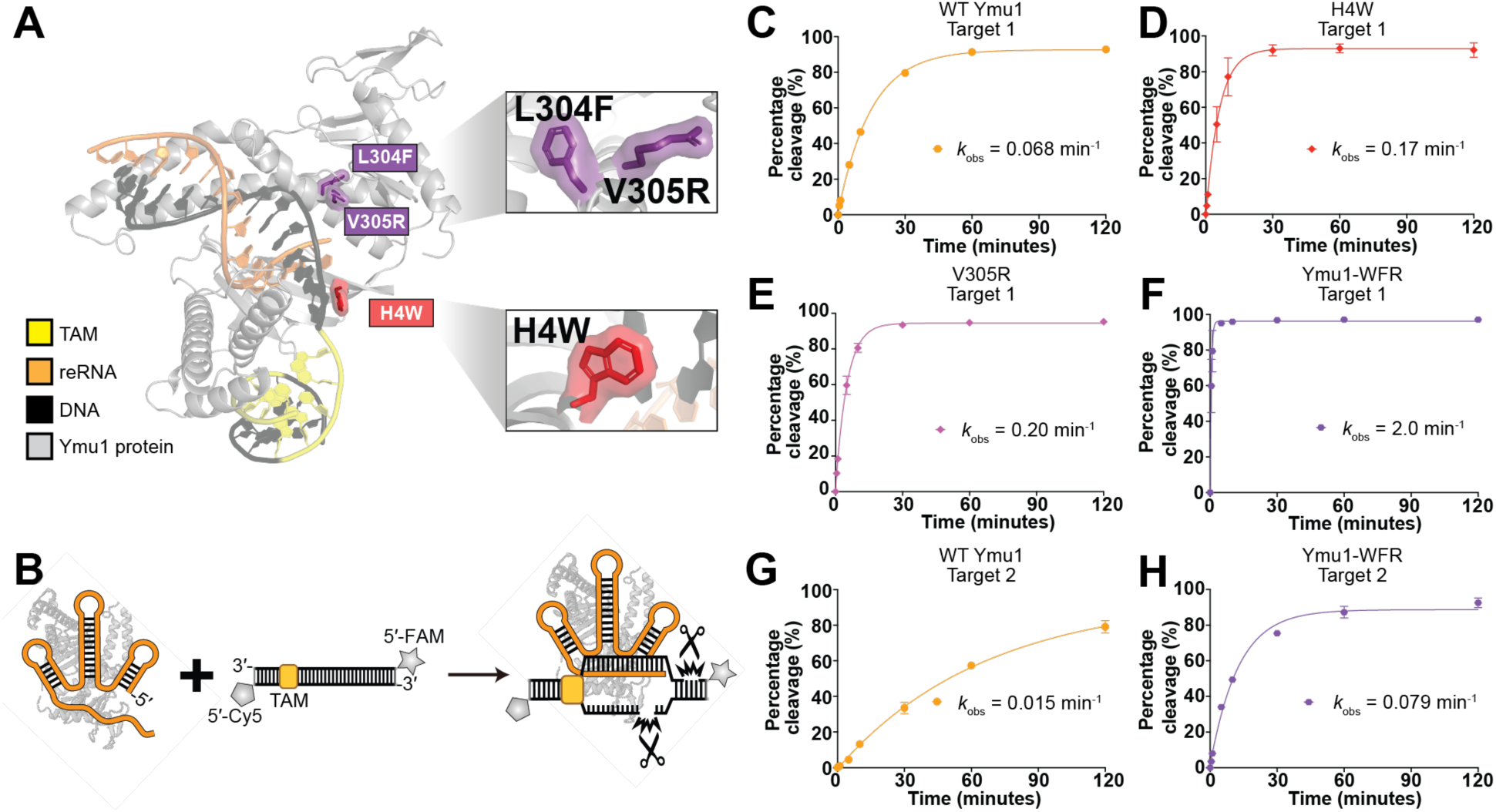
Ymu1 TnpB variants accelerate DNA cleavage *in vitro*. **(A)** Alphafold3 (AF3) model of the Ymu1-WFR ternary complex. The three substitutions (H4W, L304F, V305R) are highlighted, with zoomed views showing their local environments within the WED and RuvC lobes. **(B)** Schematic of the DNA cleavage assay using a labeled double-stranded DNA substrate. The non-target strand (NTS) is 5′-labeled with Cy5 and the target strand (TS) is 5′-labeled with FAM. **(C-H)** NTS cleavage profiles for Ymu1 TnpB variants under enzyme-excess conditions (100 nM RNP, 10 nM dsDNA) (Methods). The homogeneity of all the RNP samples were analyzed in Fig. S3A-D. Percentage of cleaved substrate (y-axis) is plotted over time (x-axis). Each point represents mean ± SD (n = 3 independent reactions); *k*_obs_ values reflect the mean from three independent mono-exponential fits on each reaction time course. Shown are **(C)** WT Ymu1 (Target 1), **(D)** H4W (Target 1), **(E)** V305R (Target 1), **(F)** Ymu1-WFR (Target 1), **(G)** WT Ymu1 (Target 2), **(H)** Ymu1-WFR (Target 2). L304F (Target 1) NTS, cleavage in the presence of salmon sperm competitor DNA, and all TS cleavage profiles are provided in Fig. S3E-P. Single-turnover behavior of the enzyme was verified under substrate-excess conditions (100 nM dsDNA, 20 nM RNP) (Fig. S3Q-R) (Methods). All *k*_obs_ values (mean ± SD) are summarized in Table S2.

### Ymu1 TnpB variants accelerate DNA cleavage *in vitro*

To understand how the enhanced variants achieve higher efficiency, we compared their biochemical cleavage kinetics *in vitro* (Fig. 3B; Methods). We measured the cleavage activities of the three single mutants (H4W, L304F, and V305R) compared to WT Ymu1 on a 60-bp dsDNA substrate corresponding to Target 1 (Fig. 3C-E and Fig. S3E-I). These cleavage assays were performed under excess enzyme conditions (Methods), and measured the overall time required for completing all steps of a single turnover starting from binding, progressing through R-loop propagation, and ending with strand scission (Note S1). Complementary assays performed with excess substrate show that the enzyme has single-turnover behavior as expected (Fig. S3Q-R; Methods). Compared to WT Ymu1 (Fig. 3C and Fig. S3F), both H4W and V305R mutants cleaved DNA faster (∼3-fold) (Fig. 3D-E and Fig. S3G-H), while the L304F mutant did not display significantly improved kinetics (Fig. S3E, S3I). Ymu1-WFR exhibited a pronounced ∼28-fold increase in cleavage rate relative to WT (Fig. 3F and Fig. S3J). To test whether this enhancement persists in the presence of excess non-target DNA, we performed cleavage assays with a 30-fold mass excess of salmon sperm DNA. Under these competitive conditions, Ymu1-WFR remained 58-fold faster than WT Ymu1 (Fig. S3K-N). To test whether the kinetic improvement was target-specific, we performed the same cleavage assays using RNPs targeting Target 2, corresponding to the *Arabidopsis PDS3* g2 site used in plants. Under the same conditions, Ymu1-WFR cleaved Target 2 > 5-fold faster than WT Ymu1 (Fig. 3G-H and Fig. S3O-P), mirroring the trend in editing efficiencies observed in plant assays (Fig. 2A, 2C and Fig. S2D). Together, these results suggest that the superior genome-editing performance of Ymu1-WFR arises from enhanced biochemical activity of the purified enzyme, motivating the mechanistic analyses that follow.

### AuRBT revealed step-specific enhancement of DNA unwinding by Ymu1 mutations

Bulk biochemical assays showed that WT Ymu1 cleaved DNA at a markedly slower rate than enhanced variants. Such kinetic limitations sometimes arise during DNA unwinding and *R*-loop formation in CRISPR-Cas enzymes^15,17–19^. Although biophysical models of *R*-loop progression and intermediate states have been established for Cas enzymes^15,19,20^, analogous information remains unexplored for TnpB. To directly resolve DNA unwinding dynamics and identify potential kinetic bottlenecks in Ymu1 TnpB, we employed gold rotor-bead tracking (AuRBT), a single-molecule torque spectroscopy method which directly reports DNA unwinding in real time with base-pair resolution^21^. In this assay, a dsDNA tether containing a single TAM adjacent to Target 1 is coupled to a gold nanoparticle, whose rotation reports changes in DNA twist (Δθ_0_) as the duplex unwinds and rewinds (Fig. 4A and Fig. S4A; Methods). To prevent cleavage and enable repeated sampling of unwinding transitions, all measurements were performed using catalytically dead WT Ymu1 (WT dYmu1, E279A) and the corresponding H4W dYmu1 and dYmu1-WFR variants (Fig. S4B-D).

**Fig. 4.**
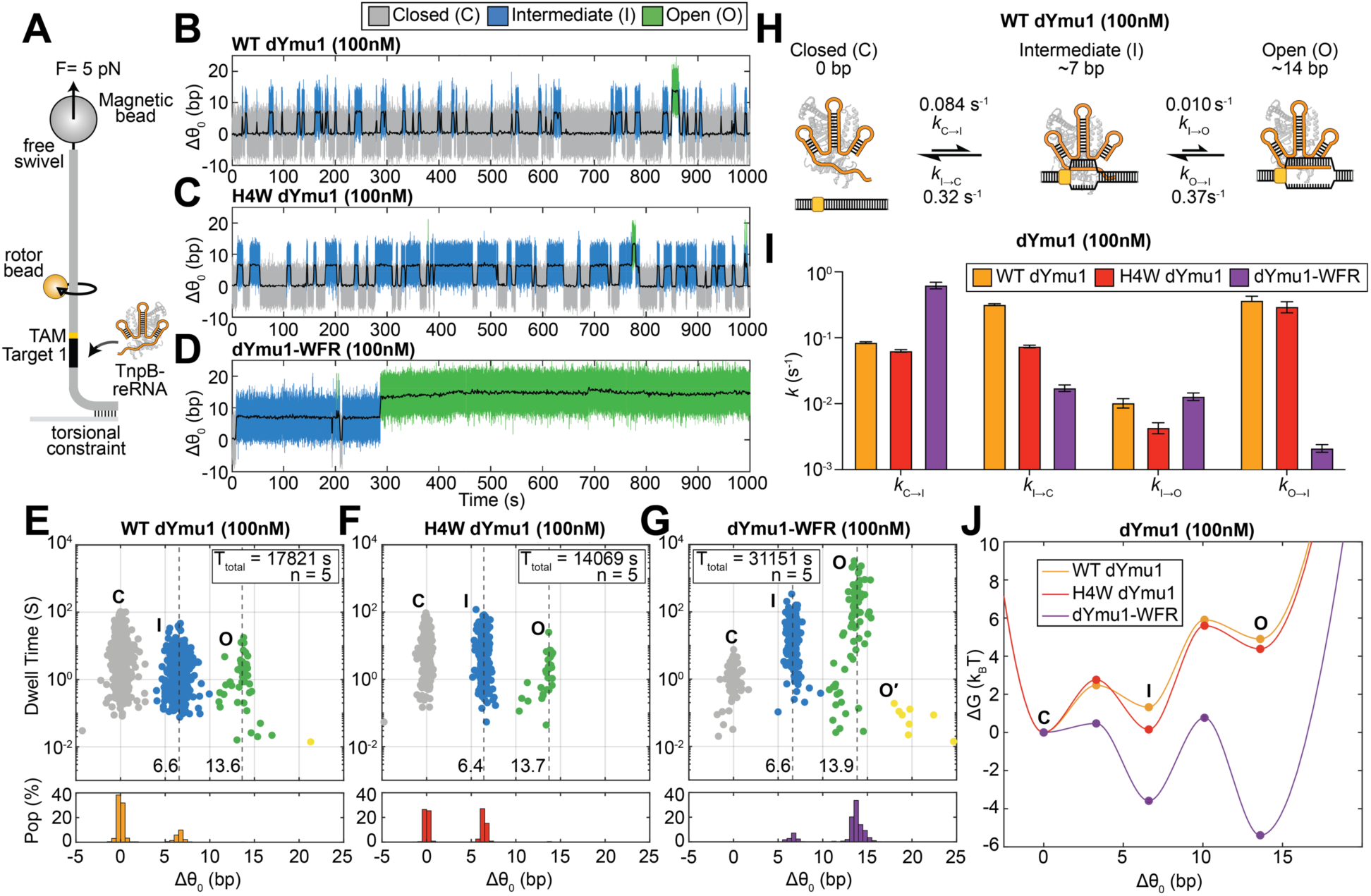
Mutations in distinct domains of Ymu1 TnpB enhance sequential steps of DNA unwinding. **(A)** Schematic AuRBT experiment for measuring *R*-loop dynamics. A single, torsionally relaxed ∼5-kb DNA molecule containing a single TAM (5′-TTGAT-3′) flanking the Target 1 is stretched between a coverslip and a magnetic bead under 5 pN tension. An 80-nm gold rotor bead (orange) is attached above the target site. In this geometry, changes in the mean rotor-bead angle report directly on changes in equilibrium twist (Δθ_0_, expressed in units of base-pairs unwound by assuming unwinding of B-form DNA). dYmu1 TnpB RNP (Fig. S4B-C), which shows no detectable dsDNA cleavage in the AuRBT buffer (Fig. S4D), was introduced at 100 nM. **(B-D)** Representative trajectories showing Δθ₀ (bp) over time (s) for **(B)** WT Ymu1, **(C)** H4W, **(D)** Ymu1-WFR. Distinct states are scored using the automated change-point detection algorithm Steppi^22,23^, including closed (C, gray), intermediate (I, blue), and open (O, green). Low-pass-filtered traces (1 Hz) are shown in black. **(E-G)** (Top) Scatter plots of state dwell times (s) versus Δθ₀ (bp) for all merged states (Methods) under equilibrium conditions. (Bottom) Histogram of occupation probability, representing the lifetime-weighted population at each corresponding Δθ_0_ with a bin size of 0.5 bp. Distinct unwinding states include closed (C, gray), intermediate (I, blue), and open (O, green). Some short-lived events are also seen with extended levels of opening (O’, yellow). The total collection time (T_total_) and the number of DNA tethers analyzed (n) are reported in the figure legend. **(E)** WT Ymu1. **(F)** H4W. **(G)** Ymu1-WFR. The unmerged scatter plots are shown in Fig. S4I-K. **(H)** Kinetic model summarizing transition rates between the closed, intermediate, and open states of WT dYmu1 at 100 nM RNP. **(I)** Bar graph of RBT transition kinetics of WT dYmu1, H4W dYmu1, and dYmu1-WFR at 100 nM RNP (Table S3). Errors were calculated assuming Poisson statistics of state transitions. **(J)** Free-energy landscapes for DNA unwinding, overlaid for WT (orange), H4W (red), and Ymu1-WFR (purple). AuRBT trace statistics, including the number of tethers analyzed, total tracking time, and number of detected transitions, are summarized in Table S4.

We examined unwinding dynamics in a torsionally relaxed DNA tether in the presence of 100 nM RNP (Fig. 4A) and observed equilibrium fluctuations in twist resulting from reversible RNP binding and *R*-loop formation. To resolve unwinding intermediates and quantify transition dynamics, we applied automated change-point detection^22,23^ to continuous trajectories, enabling identification of unwinding states as well as their dwell times and interconversion rates. All three proteins tested produced transitions between three well-separated states: a closed duplex state (C), a partially unwound ∼7 bp intermediate state (I), and a ∼14 bp fully unwound open state (O) (Fig. 4B-G). The extent of the O state is consistent with the RNA-DNA heteroduplex lengths (∼12-16 bp) required for efficient TnpB cleavage by a related ortholog (Dra2 TnpB)^2,5^. The ∼7 bp I state aligns with prior observations^2^ that Dra2 TnpB can stably bind targets containing TAM-distal mismatches beyond position ∼8, indicating a binding-competent but cleavage-incompetent *R*-loop state, reminiscent of the “seed” intermediates observed in other CRISPR-Cas enzymes^15,19,20,24,25^. Thus, Ymu1 unwinding involves a C→I initiation step that can be followed by an I→O completion step. Introducing mismatches within the first 1-4 nt of the RNA-DNA heteroduplex abolished the observed unwinding states (Fig. S4E-H), as expected for selective on-target formation of the I state.

While displaying the same three-state behavior, the Ymu1 variants have markedly different state occupancies and transition rates. At 100 nM RNP, WT dYmu1 resided predominantly in the C state, with frequent reversions from the I state and only rare, short-lived excursions to the O state (Fig. 4B, 4E, 4H, and 4I). The H4W substitution, positioned near the first RNA-DNA base pair, stabilized the intermediate state (Fig. 4C, 4F) by suppressing collapse from I to C (Fig. 4I and Table S3). In contrast, the triple substitution dYmu1-WFR reshaped the unwinding kinetics at both C↔I and I↔O stages (Fig. 4D, 4G, and 4I). dYmu1-WFR strongly suppressed collapse back to C (*k*_I→C_) (Fig. 4I and Table S3), which increases the probability of progression to the O state. In addition, dYmu1-WFR dramatically reduced reversal from the O state (*k*_O→I_), biasing the equilibrium population toward full *R*-loop formation (Fig. 4I and Table S3). Together, these kinetic effects enable efficient and persistent formation of the fully unwound state under single-molecule conditions (Fig. 4J). Collectively, these results show that H4W dYmu1 primarily enhances early unwinding by stabilizing the I state, whereas the combined dYmu1-WFR substitutions promote both initiation and propagation of the *R*-loop, providing a kinetic basis for the enhanced biochemical and cellular activities of Ymu1-WFR. Kinetic modeling of the complete reaction pathway quantitatively relates these microscopic transition rates to the bulk cleavage rates observed in biochemical assays (Note S1).

### Negative supercoiling stabilizes unwound states in WT Ymu1

Our equilibrium measurements indicate that WT dYmu1 rarely stabilizes a fully unwound *R*-loop on torsionally relaxed DNA, raising the question of how this enzyme achieves productive activity in its native bacterial context. In bacterial genomes, DNA is pervasively negatively supercoiled^26–29^. Because negative supercoiling is known to modulate strand separation and *R*-loop formation for diverse RNA-guided nucleases^15,20,24,30,31^, we tested whether DNA supercoiling alone could reshape the DNA unwinding landscape of WT Ymu1. We performed non-equilibrium AuRBT experiments in which the DNA tether was repeatedly ramped between positively and negatively supercoiled conditions (Fig. 5A; Methods)^15,20^, driving *R*-loop formation and collapse. As in prior work, changes in measured torque (Fig. S5C) were converted to changes in equilibrium twist (Δθ_0_) (Fig. 5B and Fig. S5D), and transition rates between scored *R*-loop states (Fig. 5C-D and Fig. S5E) were analyzed as a function of imposed twist (Fig. S5F-G) and used to determine supercoiling-dependent equilibrium constants (Fig. 5E) and construct free energy landscapes (Fig. 5F).

**Fig. 5.**
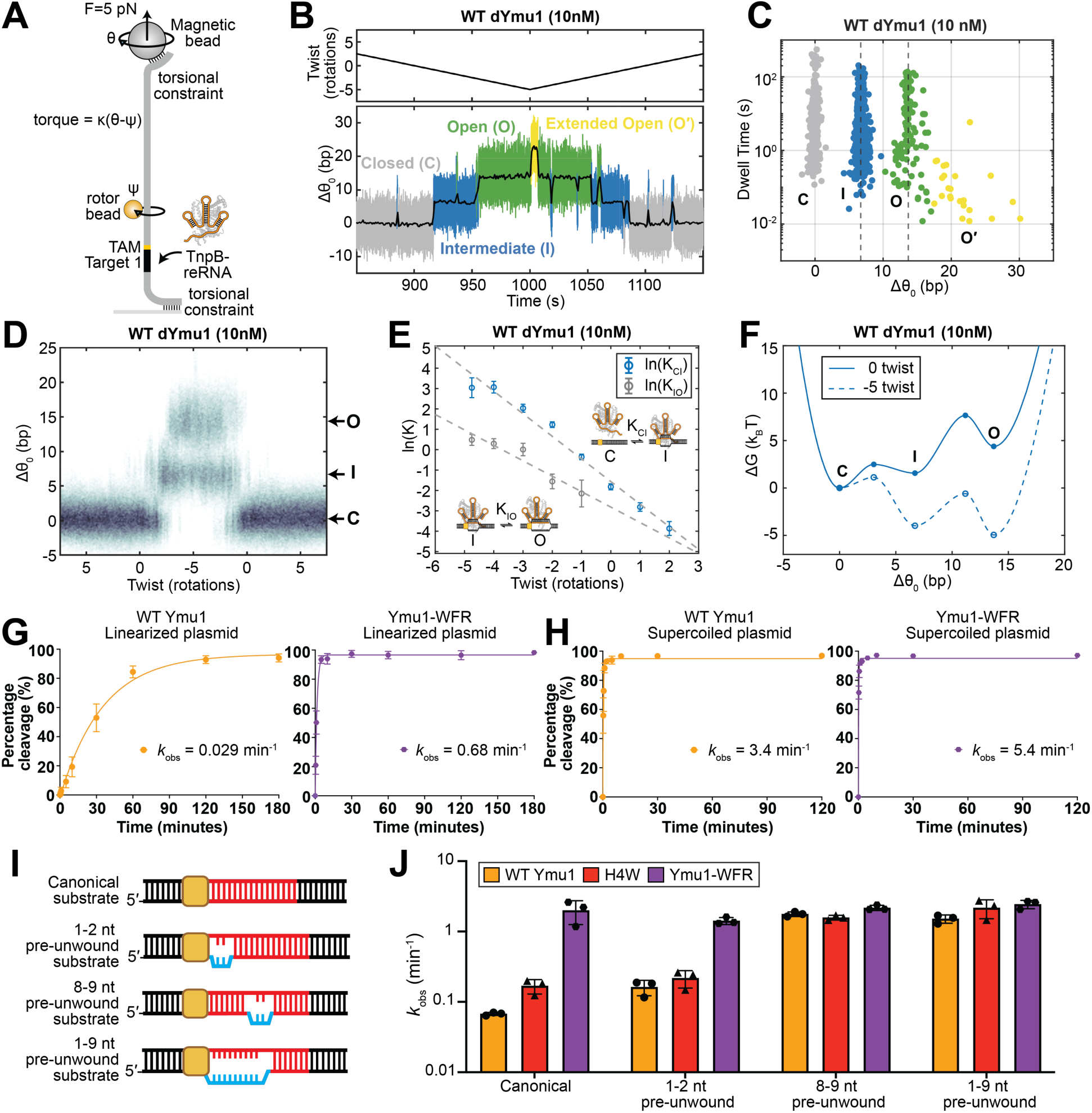
Negatively supercoiled or pre-unwound DNA enhances Ymu1 activity. **(A)** Schematic of the non-equilibrium, torque-driven AuRBT assay. Here the DNA molecule is torsionally constrained. Twist is imposed by rotating the magnets (θ), and torque is inferred from the difference in magnet position (θ), the torsional stiffness of the tether (κ) and rotor-bead angle (ψ) using τ = κ(θ − ψ). A separately prepared batch of dYmu1 TnpB RNP (Fig. S5A-B) was used at 10 nM for the torque-driven measurements. **(B)** Representative torque-driven unwinding trajectory of WT dYmu1. (Top) Imposed twist (rotations) and (Bottom) Δθ_0_ (in units of base-pairs unwound; see Methods) over time (s). Distinct states are scored using the automated change-point detection algorithm Steppi^22,23^, including closed (C, gray), intermediate (I, blue), open (O, green), and extended open (O′, yellow) states. Low-pass-filtered traces (1 Hz) are shown in black. This trace is a portion of the trajectory shown in Fig. S5D. **(C)** Scatter plot of dwell times (s) versus Δθ₀ (bp) after merging (Methods) for torque-driven measurements of WT dYmu1. Corresponding unmerged dwells are shown in Fig. S5E. **(D)** Heat map showing the distribution of Δθ₀ (bp) as a function of imposed twist (rotations). **(E)** Plot of ln(K_C→I_) and ln(K_I→O_) versus imposed twist (rotations), with linear fits. Data points were calculated from the ratio of the transition rate constants, which are plotted in Fig. S5F-G and listed in Table S5. Errors were calculated assuming Poisson statistics of state transitions. Fit parameters are summarized in Table S6. AuRBT trace statistics are summarized in Table S7. **(F)** Free-energy landscapes for DNA unwinding. Solid line: 0 rotations; dashed line: -5 rotations. Landscapes were constructed using linear fits to ln(K) and ln(*k*) data (Table S6). **(G-H)** Cleavage of linearized **(G)** and negatively supercoiled **(H)** 2.2 kb plasmid substrates by WT Ymu1 and Ymu1-WFR. Percentage of cleaved substrate (y-axis) is plotted over time (x-axis). Each point represents mean ± SD (n = 3 independent reactions); *k*_obs_ values reflect the mean from three independent mono-exponential fits on each reaction time course. See also Table S8. **(I)** Schematics of partially pre-unwound DNA substrates used for kinetic perturbation. **(J)** Bar graph of apparent cleavage rates (*k*_obs_) for WT Ymu1, H4W, and Ymu1-WFR across substrates with different initial unwinding (mean ± SD) (Tables S2 and S9). Shown here are NTS cleavage. TS cleavage profiles are provided in Fig. S5H.

The locations of metastable *R*-loop states observed during twist-ramping experiments closely matched those determined in Fig. 4H, and the O state remained poorly accessible under zero imposed twist (Fig. 5B-F). As negative supercoiling was progressively applied, both the C↔I initiation step and the I↔O propagation step shifted toward unwound states (Fig. 5B-D and Fig. S5C-E). At approximately -5 Twist (∼1% negative supercoiling), both the I and O states are substantially stabilized (Fig. 5D-F, Fig. S5F-G, and Table S5), resembling the persistent unwound states in Ymu1-WFR. Under these conditions, we occasionally observed short-lived excursions beyond the ∼14 bp O state (Fig. 5B-C and Fig. S5D-E), consistent with suggested transient downstream duplex opening required for the cleavage mechanism of TnpB^5^. As expected, negative supercoiling can partially compensate for intrinsic unwinding limitations in WT dYmu1 by stabilizing unwound conformations. Together with the variant comparisons, these measurements illustrate how DNA mechanics and protein mutations act on the same energetic landscape for *R*-loop formation.

To test whether the supercoiling dependence observed in single-molecule measurements extends to the overall cleavage activity of enzymes, we compared WT Ymu1 and Ymu1-WFR cleavage on 2.2 kb negatively supercoiled versus linearized plasmid DNA substrates. On the linearized substrate, Ymu1-WFR cleaved 23-fold faster than WT Ymu1 (*k*_obs_ = 0.68 and 0.029 min^-1^, respectively; Fig. 5G), consistent with the magnitude of enhancement observed on short oligonucleotide substrates (Fig. 3C and 3F). In contrast, on the negatively supercoiled plasmid, Ymu1-WFR and WT Ymu1 cleaved at comparable rates, differing by only 1.6-fold (*k*_obs_ = 5.4 and 3.4 min^-1^, respectively; Fig. 5H). Thus, under the conditions of this bulk assay, the rate difference between WT and WFR is largely eliminated on supercoiled DNA, with WT showing a ∼117-fold and WFR a ∼8-fold rate increase relative to linearized DNA. This is consistent with the single- molecule observation that both negative supercoiling (Fig. 5F) and WFR mutations (Fig. 4J) favor unwinding transitions, and with our conclusion that these transitions limit the rate of WT cleavage on torsionally relaxed DNA.

### Stepwise DNA substrate unwinding limits WT Ymu1 cleavage

To further challenge our model for Ymu1 function and investigate the impact of *R*-loop state stabilization in the context of the active endonuclease, we conducted cleavage assays in the presence of site-specific DNA-DNA mismatches designed to favor *R*-loop formation by reducing the energetic cost of duplex unwinding. Unlike supercoiling, which globally favors unwinding, site-specific mismatches specifically stabilize *R*-loops containing the targeted nucleotide positions. We used substrates containing localized duplex pre-unwinding to preferentially assist either C↔I (1-2 nt near the TAM), I↔O (8-9 nt), or both steps (1-9 nt) of *R*-loop formation (Fig. 5I).

For WT Ymu1, all three pre-unwound substrates produced improvements in cleavage rates, as expected, with substantially larger effects for substrates that stabilized the fully unwound O state (Fig. 5J and Fig. S5H). For H4W, the 1-2 nt pre-unwound substrate yielded only a small additional increase (Fig. 5J and Fig. S5H), which may be explained by this substitution’s stabilization of the I state, such that its formation is not strongly limiting under the conditions of the assay. By contrast, 8-9 nt or 1-9 nt pre-unwound substrates produced large enhancements (Fig. 5J and Fig. S5H), as expected since with H4W, the transition from the I to the O state remains highly inefficient, and can be favored by reducing the energetic cost of the I to O transition. Ymu1-WFR displayed uniformly high cleavage activity across all substrates (Fig. 5J and Fig. S5H). This suggests that once the C↔I and I↔O steps have both been strongly enhanced in WFR, stabilization of unwinding transitions is ineffective in accelerating overall cleavage under these conditions.

Together, these measurements show that H4W selectively stabilizes the I state, whereas WFR produces major stabilization of both the I and O states, providing a mechanistic explanation for the dramatically enhanced genome-editing efficiency of Ymu1-WFR in plants.

### Ymu1-WFR shows reduced cleavage on mismatched substrates and no detectable off-target editing

Having established that the WFR mutations enhance on-target cleavage by stabilizing DNA unwinding intermediates, we next examined the effect of guide-target mismatches on cleavage by both WT Ymu1 and Ymu1-WFR. To evaluate mismatch discrimination, we performed cleavage assays using dsDNA substrates containing single nucleotide RNA-DNA mismatches at positions 4, 7, 10, and 14 within the guide-target heteroduplex (Fig. S5I). Both enzymes displayed strong position-dependent mismatch sensitivity. Mismatches at positions 4, 7, and 10 severely inhibited cleavage for both enzymes: WT cleavage remained below 10% at 120 min (Fig. S5J-K and Table S10), and WFR cleavage rates were reduced >100-fold relative to the matched substrate (*k*_obs_ = 0.007-0.011 min⁻¹; Fig. S5J-K and Table S10). At position 14, the mismatch had a more modest effect, reducing WT cleavage ∼10-fold and WFR cleavage ∼3-fold relative to their respective matched substrates (*k*_obs_ = 0.007 and 0.62 min⁻¹, respectively; Fig. S5J-K and Table S10). These results indicate that both WT and WFR retain strong mismatch discrimination, particularly at TAM-proximal and mid-guide positions, with more modest discrimination at the TAM-distal position. The higher cleavage rates of WFR relative to WT are consistent with stabilization of the unwound states partially compensating for the energetic penalty of mismatches.

Having established the mismatch sensitivity of both enzymes *in vitro*, we next asked whether the enhanced activity of Ymu1-WFR leads to detectable off-target editing in plants. To address this, we performed whole-genome (2,406-5,605× coverage) sequencing of three transgene-free *Arabidopsis* progeny lines derived from a parent plant in which Ymu1-WFR achieved 70.3% and 65.1% somatic editing at the *CHLI1* g4 and *PDS3* g12 sites, respectively^16^. All three progeny harbored edits at both loci. Cas-OFFinder^32^ identified 82 and 48 candidate off-target sites for *CHLI1* g4 and *PDS3* g12 (Methods), respectively, and no off-target SNPs or indels were detected at any predicted sites (Tables S11-12). After filtering against control samples, only 54 unique variants were identified genome-wide, predominantly SNPs rather than the indels characteristic of TnpB cleavage, and none overlapped with predicted off-target sites (Table S13). Thus, under the conditions tested, no off-target editing by Ymu1-WFR was detected.

Together with the biochemical and single-molecule analyses described above, these data support a model in which the WFR mutations reshape the unwinding energy landscape to enhance on-target activity (Fig. 6) while preserving genome-wide specificity.

**Fig. 6.**
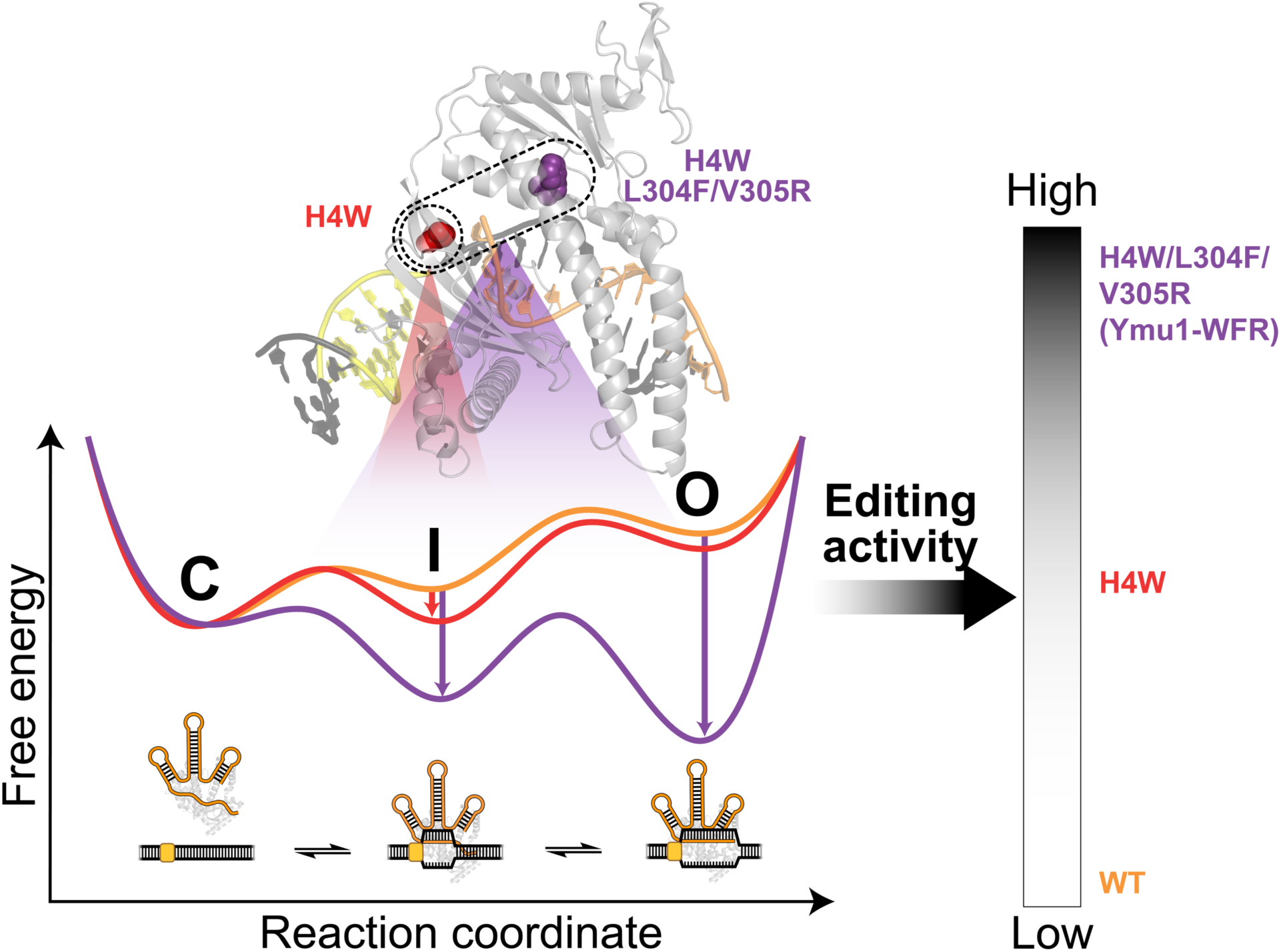
Mechanistic model for how Ymu1 TnpB mutations reshape the unwinding energy landscape to enhance genome editing. The schematic model illustrates how the three mutations in Ymu1-WFR modulate DNA-unwinding along a three-step pathway that includes closed (C), intermediate (I) and open (O) states. H4W stabilizes the I state, whereas L304F and V305R, positioned in the lid-adjacent helix, further stabilize both the I and O states. Together, these effects yield progressively higher genome-editing efficiencies from WT→H4W→WFR. Our model suggests that tuning local energetic checkpoints for *R*-loop formation provides a generalizable strategy for rational engineering of compact RNA-guided nucleases with enhanced activity

## DISCUSSION

RNA-guided nucleases recognize DNA through the energetics of base-pair exchange, yet the quantitative architecture of this process has remained poorly defined for compact enzymes such as TnpB. Our work reveals that TnpB traverses an unwinding pathway that includes formation of a partially unwound DNA intermediate state followed by a fully unwound open state. Under relaxed DNA conditions, both states are prone to collapse, and reversal is favored over progression from the intermediate to the open state. Consequently, forward progress into the fully unwound state is intrinsically limited, and TnpB-catalyzed DNA cleavage occurs only during rare, transient visits to this configuration. Thus, the stability and dynamics of conformational states upstream of strand scission constrain TnpB activity. These conformational dynamics are sensitive to protein mutagenesis and DNA supercoiling and may be further modulated by genomic context and target sequence. Stepwise *R*-loop formation has been described for Cas12 enzymes^15^ that share structural homology with TnpB. However, structural elaborations in Cas12a, including a large REC2 domain, provide extensive contacts during late *R*-loop formation that stabilize the fully unwound state and promote nuclease activation^33^; these contacts are absent in the minimal TnpB scaffold^2^, which may contribute to the transient nature of fully unwound state observed here.

Consistent with prior deep mutational scanning of Dra2 TnpB, which uncovered numerous activity-enhancing substitutions^12^, our results suggest that TnpB are not optimized for maximal DNA-cutting activity in nature. The reversibility of DNA substrate unwinding combined with strict TAM dependence naturally bias TnpB toward reversible target sampling rather than favoring the catalytically competent *R*-loop conformation. Such properties are consistent with longstanding hypotheses that transposon-associated nucleases need to balance mobility with host genome stability^4,12^, although how the bacterial host’s pervasive negative supercoiling modulates this balance *in vivo* remains an open question. When deployed in eukaryotic genomes for genome editing, the local DNA topology may also modulate this balance: localized negative supercoiling arising from transcription^34^ could partially stabilize the O state for WT TnpB at some loci, while localized positive supercoiling could further suppress *R*-loop formation, contributing to the target-site-dependent variation in editing efficiency observed in Fig. 2D.

By defining the quantitative contributions of each DNA unwinding step, our work also illuminates a modular route for enzyme improvement. The three activating mutations identified here act at distinct energetic states: H4W facilitates *R*-loop initiation, whereas the addition of L304F and V305R stabilizes more extensive DNA unwinding. This separation of effects demonstrates that compact nucleases can be modulated through a small number of targeted point mutations rather than extensive redesign of the protein scaffold. Structural hypotheses informed by sequence covariation, AF3 modeling, and cryo-EM structures of related TnpB orthologs (Fig. S6) suggest how these mutations may reshape the unwinding landscape, with H4W stabilizing early RNA-DNA engagement and L304F/V305R affecting protein conformational features associated with later unwound states (Fig. S6; Note S2). This framework retrospectively explains why mutations at homologous positions (N4 and I304 in Dra2, corresponding to H4 and V305 in Ymu1) enhance Dra2 TnpB activity in DMS^12^, suggesting that similar energetic bottlenecks exist across TnpB orthologs, even though the optimal substitution at each position can be ortholog-specific. Consistent with this framework, these activating positions have been independently shown to enhance editing across both TnpB orthologs in human cells and plants^12,16,35^.

Taken together, these findings position TnpB as a mechanistic reference point for understanding how RNA-guided systems diversified from compact ancestors. The reversible, stepwise unwinding pathway we describe provides a plausible framework by which CRISPR-associated nucleases could have evolved faster DNA cleavage kinetics through stabilization of specific unwound intermediates^15,19,20^. In addition, stabilization of the fully unwound state may also be critical for emerging applications of TnpB that rely on durable DNA binding without cleavage, such as transcriptional modulation^36^ and base editing^37^. More broadly, comparative unwinding-landscape measurements across TnpB orthologs and related RNA-guided enzymes including IscB and Fanzors, where activity-enhancing variants are already emerging^38–41^, may enable further rational engineering of compact RNA-guided nucleases.

### Limitations of the Study

This study provides mechanistic insights into how DNA unwinding dynamics govern TnpB activity; however, several limitations should be considered. First, our biochemical cleavage assays were performed on short oligonucleotide or plasmid substrates in purified systems, which do not fully recapitulate the complexity of target search in cellular environments, including chromatinized genomic DNA. Second, the genome-editing and off-target experiments in this study were conducted exclusively in *Arabidopsis*, and whether the enhanced activity and specificity profile of Ymu1-WFR extend to mammalian or other cellular contexts has not been assessed. Third, the mechanistic conclusions rest on kinetic and thermodynamic measurements without direct structural validation. High-resolution cryo-EM structures of Ymu1-WFR in complex with target DNA would provide atomic-level insight into how the identified mutations alter the *R*-loop propagation energy landscape. Finally, the mechanistic model developed here is based on single-molecule measurements of a single TnpB ortholog (Ymu1) using one target DNA sequence. Extending these measurements to additional target sequences, guide-target mismatches, supercoiling-dependent conditions, as well as to other Ymu1 variants and TnpB orthologs, would further test and generalize the stepwise unwinding framework.

### Resource Availability

#### Lead Contact

Further information and requests for resources and reagents should be directed to and will be fulfilled by the lead contact, Jennifer A. Doudna.

#### Materials Availability

Plasmids generated in this study will be deposited to Addgene upon publication. Addgene IDs will be available in the key resources table. This study did not generate new unique reagents.

#### Data and Code Availability

- NGS sequencing data have been deposited in the National Institutes of Health NCBI SRA under the BioProject ID: PRJNA1432103 (Amplicon sequencing), PRJNA1482598 (Whole-genome off-target sequencing), PRJNA1451272 (Bacteria TAM assay), and in the Gene Expression Omnibus under accession GSE327273 (small RNA-seq and SHAPE-MaP), and are publicly available as of date of publication. Single-molecule data have been deposited in the Stanford Digital Repository (https://doi.org/10.25740/qg757fx8501) as MATLAB .fig files and are publicly available as of the date of publication. All other source data, including biochemical analysis data points, genome-editing rates, uncropped gel images and processed single-molecule data used to generate figures, have been deposited at Mendeley (https://doi.org/10.17632/gdpxk2xszx.1) and are publicly available as of the date of publication.
- Customized codes for genome-editing amp-seq analysis (https://doi.org/10.5281/zenodo.20838702) and TAM analysis (https://doi.org/10.5281/zenodo.20838708) have been deposited online and made publicly available on Github. Customized MATLAB code for single-molecule data analysis has been deposited in the Stanford Digital Repository (https://doi.org/10.25740/qg757fx8501).
- Any additional information required to reanalyze the data reported in this paper is available from the lead contact upon request.

## Acknowledgements

We thank members of the Doudna, Bryant and Jacobsen labs for helpful discussions. We thank Kaihong Zhou and Jinjuan Ye for invaluable technical and organizational support of the Doudna laboratory, and Keana Lucas for outstanding leadership and coordination in managing the laboratory’s operations and research activities. We acknowledge Ms. Netravathi Krishnappa (Center for Translational Genomics, Innovative Genomics Institute, UC Berkeley) and Dr. Suhua Feng (High-Throughput Sequencing Core, Broad Stem Cell Research Center, UCLA) for Next-generation Sequencing (NGS). We thank Dr. Colette Picard for assistance with NGS data analysis. Schematics in Fig. 1E and Fig. 2B were created with BioRender.com. I.S.-D. is a Stanford Bio-X Undergraduate Fellow. H.S. is supported in part by the Jane Coffin Childs Memorial Fund for Medical Research and a K99 award from the National Institutes of Health (K99GM160778 to H.S.).

M.M.R. was supported as a summer research student by Emerson Collective. P.H.Y., J.A., N.A-S., B.W.T., and R.F.W. are recipients of the National Science Foundation Graduate Research Fellowship. E.E.D. is supported by a fellowship award from the National Institutes of Health (F32GM153031 to E.E.D.). K.M.W. is supported by the Bakar BioEnginuity Impact Grant. J.A.D., S.E.J. and D.F.S., are Investigators of the Howard Hughes Medical Institute (HHMI). This project was supported by a National Science Foundation Plant Genome Research Program grant (2334027 to S.E.J., and J.A.D.). Single-molecule biophysical components of this project were supported by the National Institutes of Health (R01GM106159 to Z.B.). HHMI has covered open access publication charges.

## Author contributions

**Conceptualization:** Z.Z., H.S., I.S.-D., T.W., S.E.J., Z.B., J.A.D.

**Methodology:** Single-molecule RBT assays: I.S.-D., K.D.P.A., N.A.S; DNA cleavage assays: Z.Z., H.S.; Plant editing assays: T.W.

**Investigation:** Single-molecule RBT assays: I.S.-D.; Plasmid cloning: Z.Z., H.S., T.W., K.V., M.K., K.W.M.; RNP reconstitution: Z.Z., H.S.; DNA cleavage assays: Z.Z., M.M.R.; Plant editing assays: T.W., M.K., J.A.; RNA-seq: K.V., E.E.D.; SHAPE-MaP: Z.Zhou, C.L.; TAM assays: J.A.

**Formal Analysis:** Single-molecule RBT assays: I.S.-D.; DNA cleavage assays: Z.Z; Plant editing assays: T.W., Y.Z.; Phylogenetic and structural analysis: H.S., P.S., P.H.Y.; RNA-seq: M.I.T.; SHAPE-MaP: C.L.; TAM assays: J.A.

**Software:** Single-molecule RBT assays: I.S.-D.; Phylogenetic and structural analysis: P.S., P.H.Y.; RNA-seq: M.I.T.

**Resources:** S.E.J., Z.B., J.A.D., P.S., I.E.-G., E.C.D., B.W.T., R.F.W., D.F.S.

**Visualization:** Z.Z., I.S.-D., H.S., T.W., P.S., P.H.Y.

**Validation:** Single-molecule RBT assays: I.S.-D.; DNA cleavage assays: Z.Z., M.M.R.; Plant editing assays: T.W., M.K., J.A.

**Supervision:** J.A.D., Z.B., S.E.J.

**Project Administration:** Z.Z., H.S., J.A.D.

**Funding Acquisition:** J.A.D., Z.B., S.E.J.

**Writing – Original Draft:** Z.Z., H.S.

**Writing – Review & Editing:** Z.Z., H.S., J.A.D., Z.B., I.S.-D., T.W., M.K., P.H.Y., B.W.T., R.F.W., D.F.S.

## Declaration of generative AI and AI-assisted technologies

During the preparation of this work, the authors used GPT-5 to assist with text editing. After using this tool or service, the authors have reviewed and edited the content as needed, and take full responsibility for the content of the publication.

## Supplemental information

Document S1: Figures S1-S6, Tables S1-S23, Notes S1-S2.

## STAR Methods

### KEY RESOURCES TABLE

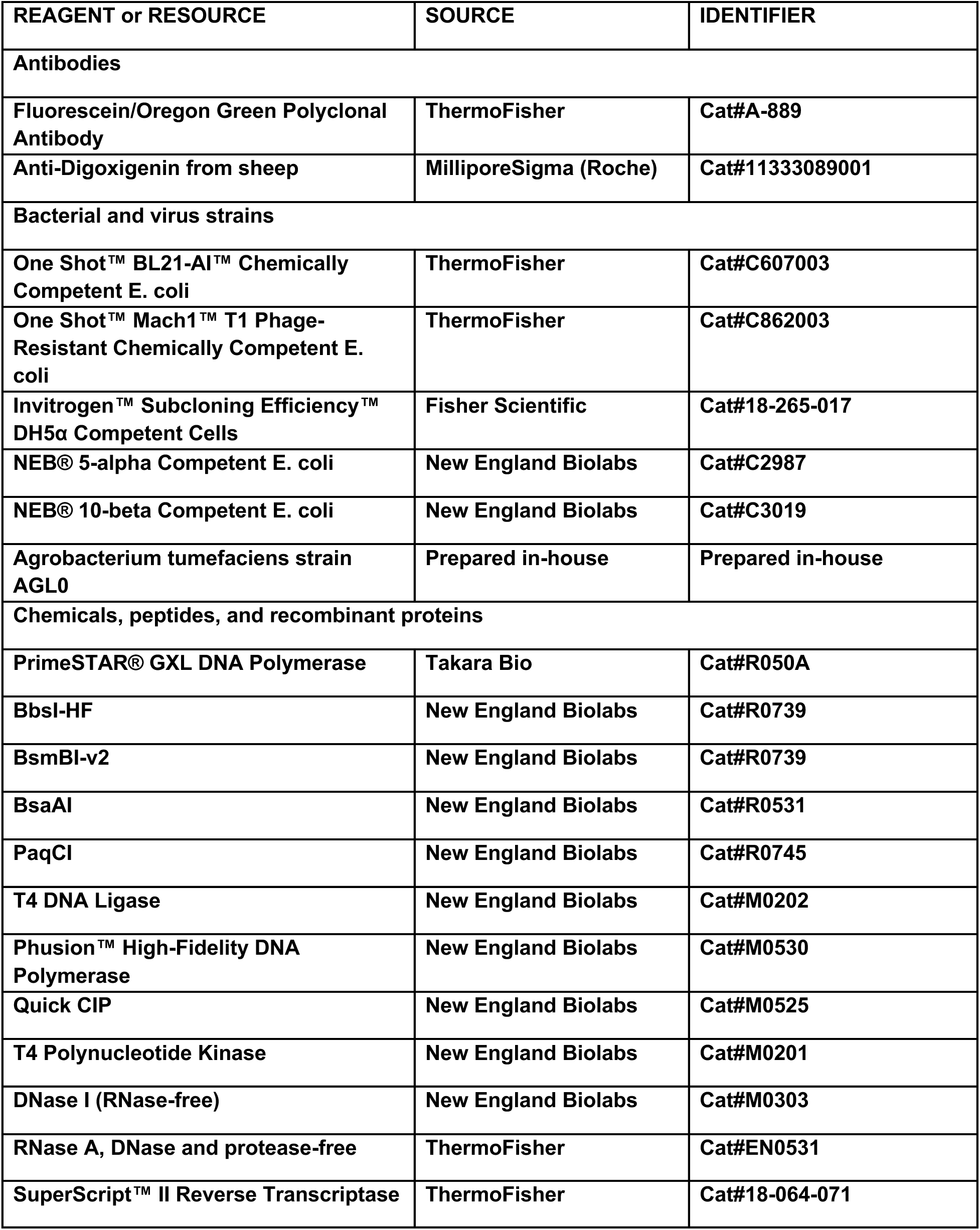
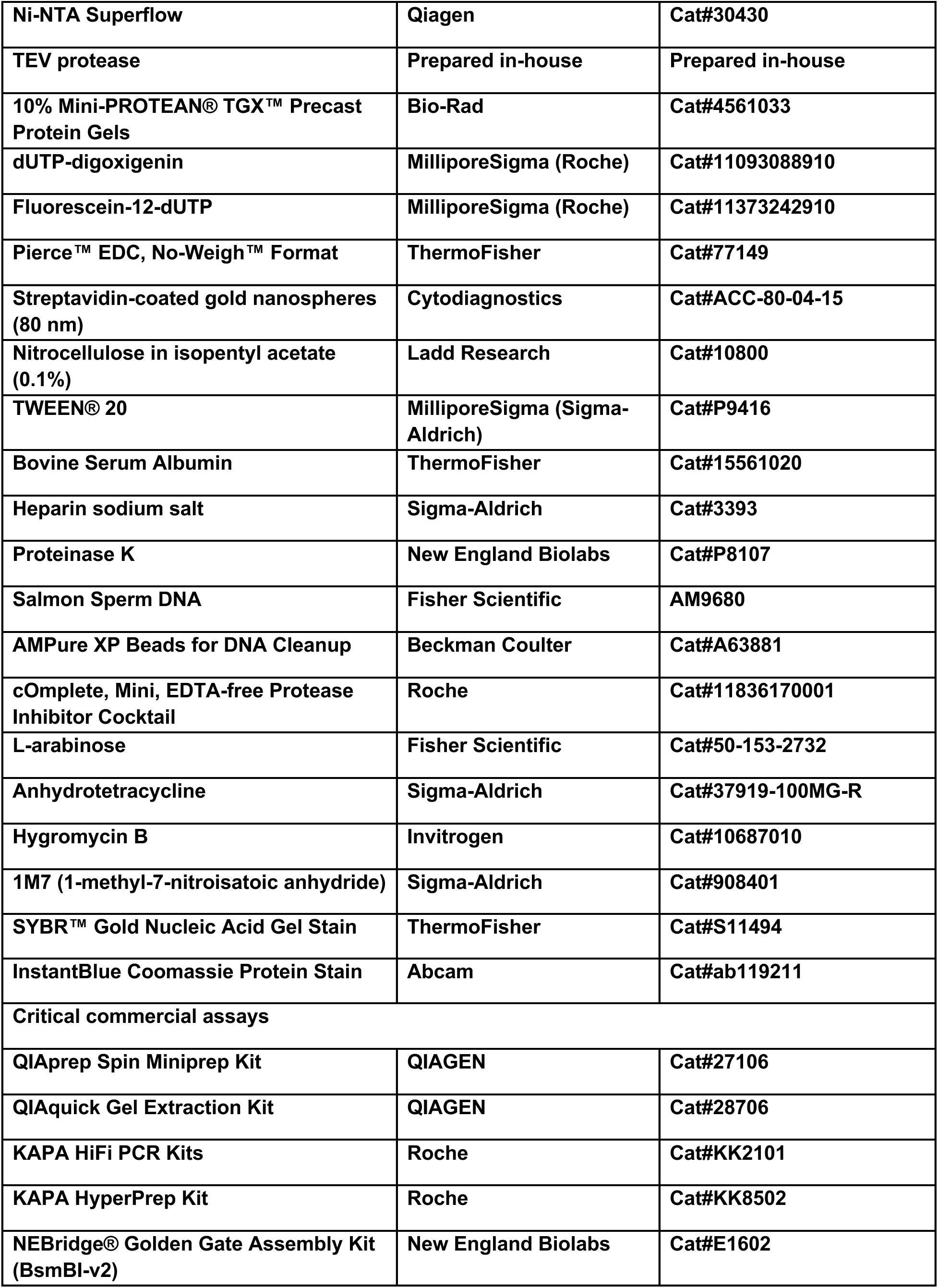
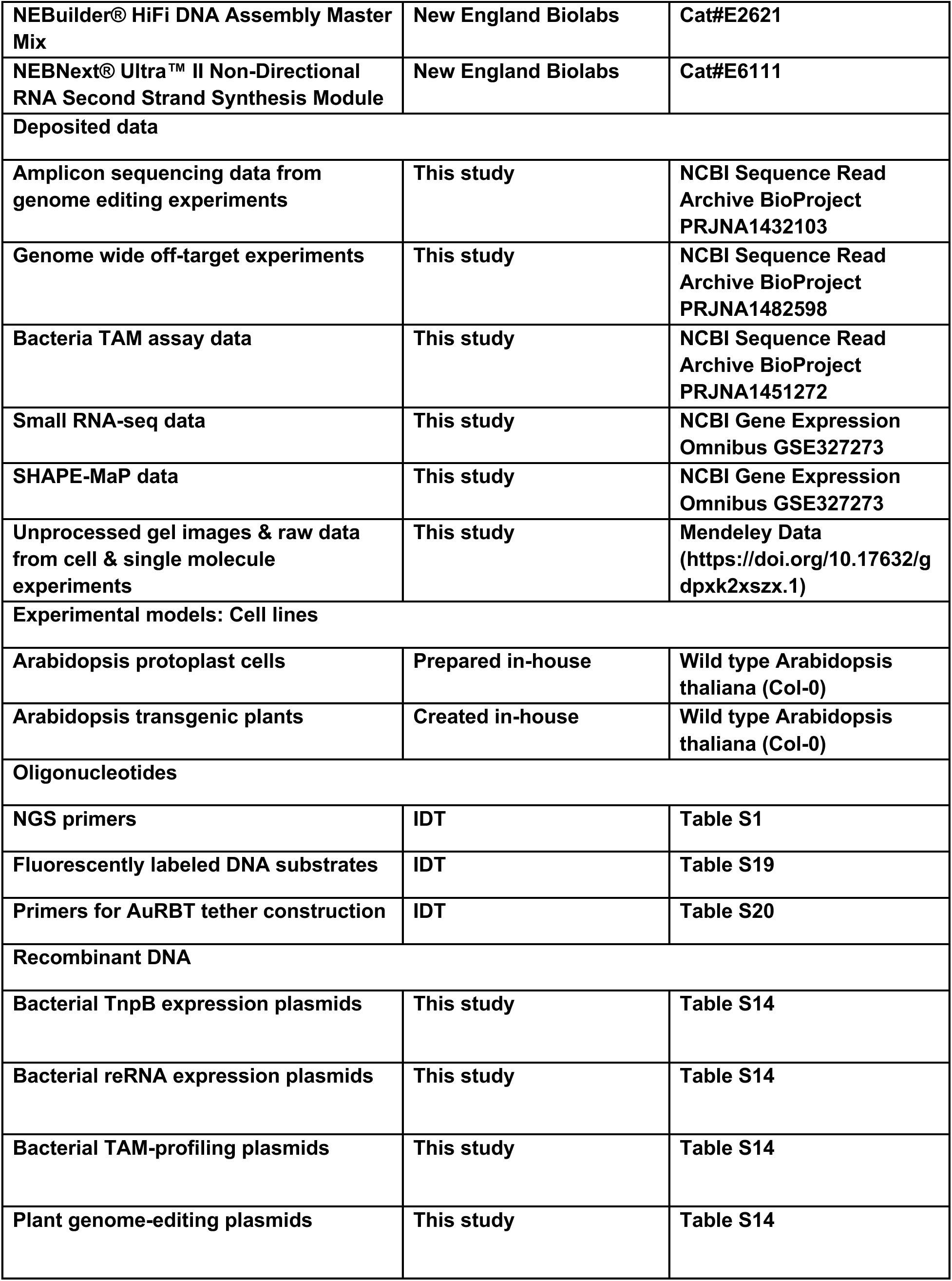
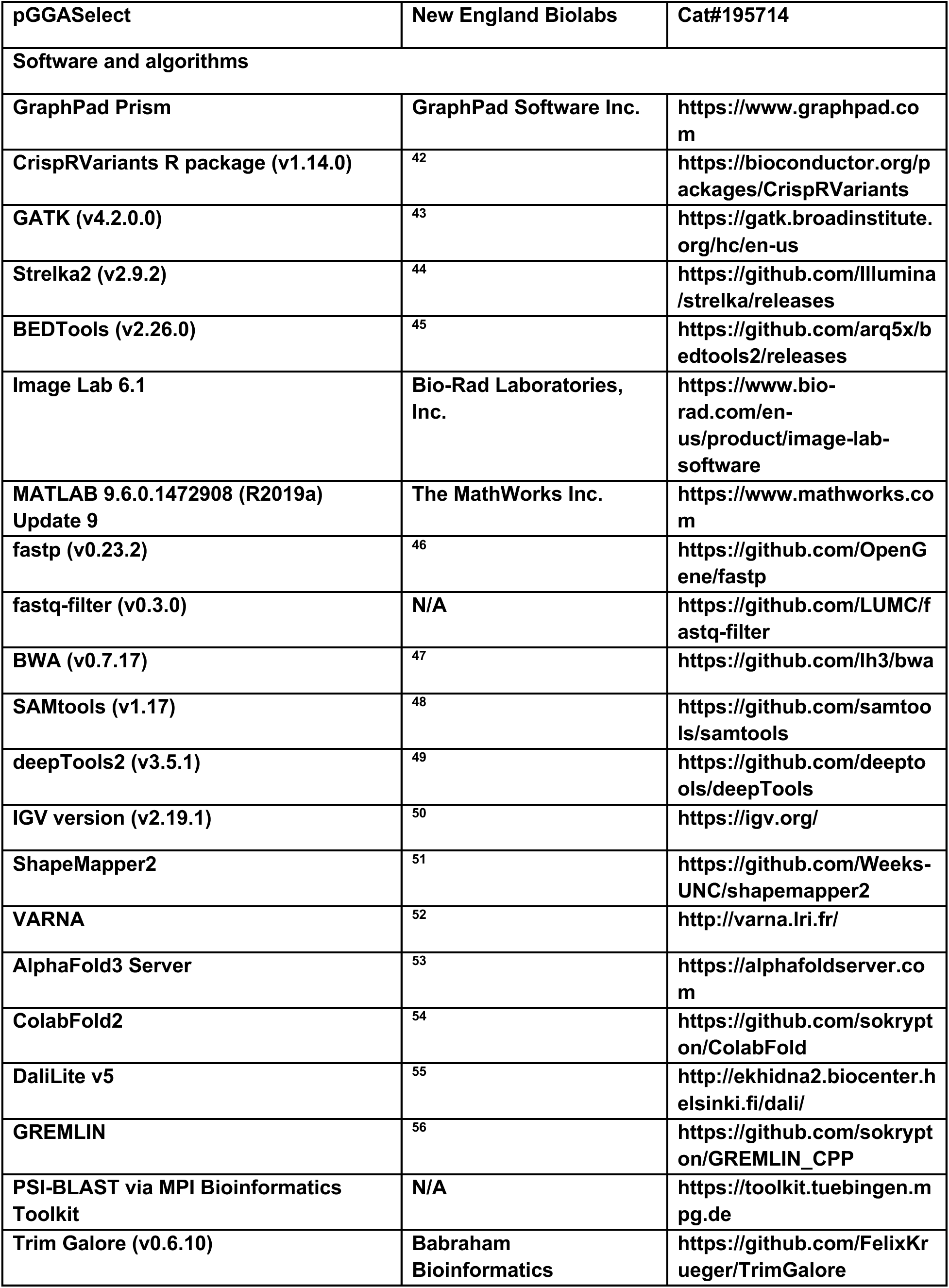
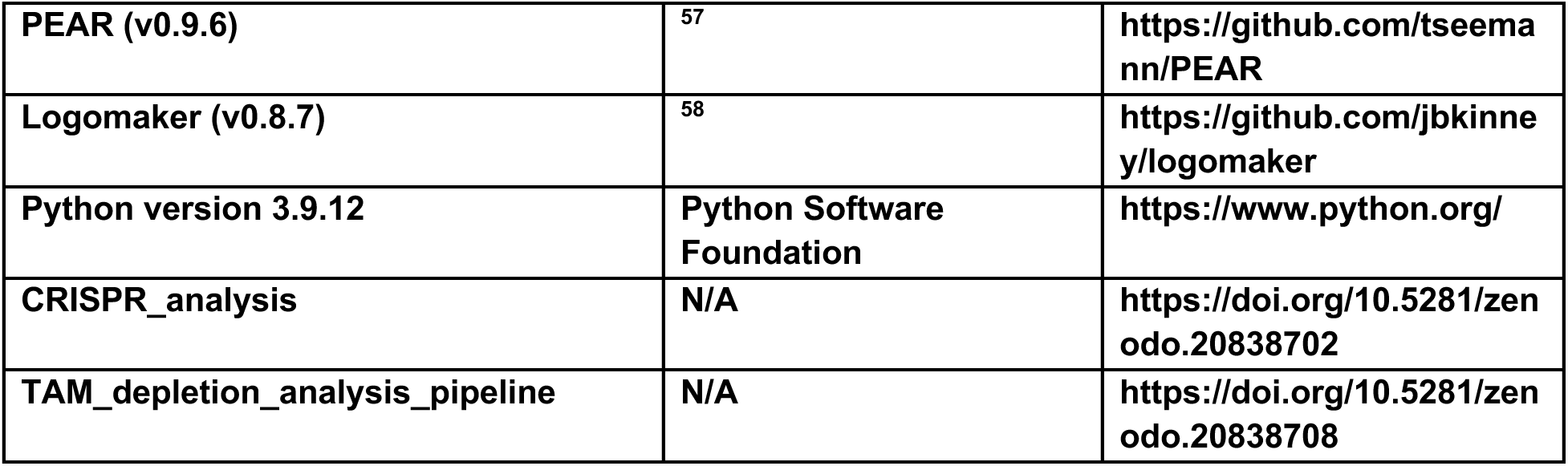

#### Experimental Model and Study Participant Detail

##### Plant materials

Arabidopsis thaliana Columbia ecotype (Col-0) was used for all plant genome editing experiments in this study. For protoplast experiments, wild-type Col-0 seeds were grown on Jiffy pucks under a 12-h/12-h light/dark photoperiod with low-light conditions at 20 °C for approximately 4 weeks prior to mesophyll protoplast isolation. For transgenic plant editing experiments, wild-type Col-0 plants were transformed using the standard floral dip protocol with the Agl0 Agrobacterium strain. T1 transgenic seedlings were selected on ½ MS plates containing 40 μg/mL hygromycin B. For whole-genome off-target analysis, three transgene-free Col-0 progeny lines harboring heritable edits generated by Ymu1-WFR were obtained from a previous study^16^.

##### Bacterial strains

Escherichia coli BL21-AI cells (ThermoFisher Scientific, C607003) were used for TnpB RNP expression and purification. DH5α (Fisher Scientific, 18-265-017), NEB 5-alpha (New England Biolabs, C2987), Mach1 T1R (ThermoFisher Scientific, C862003), and NEB 10-beta (New England Biolabs, C3019) competent cells were used for plasmid cloning and propagation. NEB 10-beta cells were additionally used for bacterial TAM depletion assays.

#### Method Details

##### Plasmid construction

All plasmids used in this study were constructed using a combination of PCR amplification, Gibson assembly, and Golden Gate assembly. Primers were synthesized by Integrated DNA Technologies (IDT). PCR reactions were performed with PrimeSTAR GXL DNA Polymerase (Takara Bio, R050A) according to the manufacturer’s instructions. Synthetic DNA fragments (gBlocks) were ordered from Twist Bioscience. For Golden Gate cloning, we used BbsI-HF (New England Biolabs, R3539), BsmBI (New England Biolabs, R0739), and PaqCI (AarI; New England Biolabs, R0745) in combination with T4 DNA Ligase (New England Biolabs, M0202), or used the NEBridge Golden Gate Assembly Kit BsmBI-v2 (New England Biolabs, E1602). Gibson assemblies were performed with the NEBuilder HiFi DNA Assembly Master Mix (New England Biolabs, E2621). All constructs were transformed into either DH5α (Fisher Scientific, 18-265-017), NEB 5-alpha (New England Biolabs, C2987H), Mach1™ T1ᴿ (ThermoFisher Scientific, C862003) or NEB 10-beta (New England Biolabs, C3019) competent *E. coli* cells and verified by Plasmidsaurus whole-plasmid sequencing to confirm complete construct integrity. A complete list of key plasmids, primer sequences, guide sequences, and vector maps is provided in Tables S1 and S14-18.

###### Bacterial TnpB protein expression plasmids

To express TnpB proteins in bacteria, we built pBAD-10×His-MBP-TEV-TnpB expression vectors. Coding sequences (CDSs) for TnpB homologs (Ymu1, Dra2, Tel2, Tfu1, and Ec41) were obtained either from a previously described Ymu1 plasmid **pMK061**^8^ or as gBlocks that included both the CDS and the downstream noncoding region encoding the reRNA^6^. For bacterial expression, all TnpB CDSs were cloned into **pHS354**, a BbsI-based dropout vector containing an araBAD promoter, an N-terminal 10×His-MBP-TEV tag, and an mKate stuffer cassette flanked by BbsI sites. Each CDS was PCR-amplified with primers introducing BbsI-compatible overhangs and inserted into **pHS354** by BbsI digestion and Golden Gate ligation, replacing the mKate stuffer with the TnpB open reading frame and yielding 10×His-MBP-TEV-TnpB constructs (Table S16). Point mutations (H4W, L304F, V305R), the catalytic inactivation mutation (dYmu1/E279A), and combinatorial variants (Ymu1-WFR, dH4W, dWFR) were generated using primer-encoded substitutions. Mutant constructs were assembled by GXL PCR followed by two-fragment Gibson assembly.

###### Bacterial reRNA expression plasmids

To express reRNAs in bacteria, we built T7-reRNA-HDV expression vectors. We first constructed pHS357, a BsmBI-based dropout vector containing a T7 promoter, a BsmBI stuffer cassette, and a 3′ HDV ribozyme. The reRNA scaffolds and their corresponding 16-nt guide sequences were PCR-amplified with primers introducing BsmBI overhangs and inserted into **pHS357** by replacing the stuffer cassette via BsmBI Golden Gate assembly (Table S17). For Ymu1, we tested two reRNA architectures: a 200-nt scaffold corresponding to the native genomic noncoding region (pHS610) and a shorter 127-nt processed scaffold identified by bacterial small-RNA sequencing (pHS607). Notably, expression of the 200-nt scaffold in bacteria yields a processed ∼127-nt reRNA species (Fig. S1D), consistent with endogenous RNA processing. Direct expression of the 127-nt scaffold resulted in higher yield of RNP in bacteria and was used for all DNA cleavage assays and equilibrium AuRBT measurements. The 200-nt scaffold construct was used to reconstitute the dYmu1 RNP employed in torque-driven AuRBT experiments. For all reRNA-expression constructs, we tested guide sequences corresponding to both Target 1 (**5′-TCTTCTGGATTGTTGT-3′**)^14,15^ and Target 2 (**5′-AAGGCAAATTCGCCGC-3′**, targeting *PDS3* g2 site)^8^.

###### Bacterial TAM-profiling assay plasmids

Single-transcript TnpB-reRNA constructs for bacterial TAM profiling were expressed from a tetracycline-inducible TetR/pTet promoter, with TnpB and reRNA encoded on the same transcript and an HDV ribozyme positioned at the 3′ end to define the reRNA terminus, as described previously^8^. The WT Ymu1 single-transcript construct (**pHS516**) was used as reported. To generate the Ymu1-WFR variant (**pHS751**), the three substitutions (H4W, L304F, and V305R) were introduced by PCR using mutagenic primers, and the single-transcript cassette was reassembled by Gibson assembly into the same expression scaffold.

###### Plasmid substrates for cleavage assays

To generate the plasmid substrate used in supercoiled versus linearized cleavage assays, a preannealed oligonucleotide duplex encoding the Target 1 sequence and TAM with 4-bp overhangs compatible with BsmBI-generated ends was cloned into the pGGAselect Destination Vector (New England Biolabs, N0309) using the NEBridge Golden Gate Assembly Kit (BsmBI-v2) (New England Biolabs, E1602). The resulting plasmid (pHS697, ∼2.2 kb) was verified by whole-plasmid sequencing. The full plasmid sequence is provided in Table S18.

###### Plant genome-editing plasmids

Plant genome-editing constructs for *Arabidopsis* thaliana consisted of a single transcriptional unit expressing TnpB, the reRNA, and a 3′ HDV ribozyme (TnpB-reRNA-HDV), driven by the UBQ10 promoter and terminated by the RbcS-E9t terminator.

To construct a subset of Ymu1 protein variant plant editors targeting *PDS3* g2 locus, we used **pMK525**, a TnpB-reRNA ccdb dropout cloning vector for PaqCI golden gate cloning to insert gene fragments (from Twist Bioscience) containing the Ymu1 TnpB, reRNA, and *PDS3* g2 sequence. Additional plant-editing vectors targeting other endogenous genomic loci were created by cloning guide sequences into Ymu1 guide ccdb dropout vectors **pMK025**, **pTW2503**, or **pTW2543**. Guide sequences were introduced by phosphorylating and annealing complementary IDT oligos and inserting the resulting duplex into **pMK025**, **pTW2503**, or **pTW2543** using PaqCI Golden Gate assembly. All the endogenous target sequences are listed in Table S1. All assembled plasmids were transformed into NEB 10-beta cells and validated by whole-plasmid sequencing. The benchmark WT Ymu1 protoplast experiments (Fig. 1F) and all whole-plant data in the main figures (Fig. 2C-D) were generated using the 127-nt reRNA scaffold. For the *PDS3* g2 locus, WT Ymu1, H4W, and Ymu1 WFR were additionally tested in parallel using the short reRNA scaffold to assess compatibility (Fig. S2D). All mutagenesis screening experiments in protoplast were performed using the short reRNA scaffold (Fig. 2A).

For high-throughput screening of protein mutations in *Arabidopsis* protoplasts targeting *PDS3* g2, we constructed a single-transcript vector (**pKV86**) containing a short reRNA scaffold derived from **pMK061**. To generate **pKV86**, we amplified two backbone fragments (12 kb and 800 bp) from **pMK061** and assembled them with a 200-bp gBlock encoding the short reRNA plus guide sequence in a three-part Gibson reaction. To generate TnpB variant constructs for protoplast screening, we used **pHS550** as the backbone, which is a PaqCI dropout plasmid containing an mRFP1 stuffer under the UBQ10 promoter, followed by an HDV ribozyme and the RbcS-E9t terminator. Point mutations were encoded directly on PaqCI-compatible primers. The TnpB-short-reRNA-guide cassette (as in **pKV86**) was split into two non-overlapping PCR fragments, with the junction positioned such that each desired mutation resided on one fragment. These fragments were amplified separately and assembled with the **pHS550** backbone using a three-part PaqCI Golden Gate reaction, yielding complete single-transcript TnpB-reRNA-HDV constructs. Combinatorial variants were generated similarly, except that the cassette was divided into three PCR fragments to accommodate multiple mutated regions, which were assembled with the **pHS550** backbone in a four-part PaqCI Golden Gate reaction.

##### Plant genome-editing experiments

Protoplast experiments were performed as described in a previous study^8^. WT *Arabidopsis* Columbia ecotype (Col-0) seeds were grown on Jiffy pucks under 12-h/12-h light/dark photoperiod with low-light condition at 20 °C for about 4 weeks. Mesophyll protoplast isolation was then performed as described previously^59^. Isolated protoplast cells were transfected with 20 µg of plasmid and incubated in 6-well plates at 26 °C for 48 h with a 37 °C heat-shock treatment for 2 h at 16 h post transfection. At 48 h post transfection, protoplasts were collected for genomic DNA extraction.

For transgenic plant editing experiments, the *Arabidopsis* Columbia ecotype (Col-0) was used. Transgenic plants were created using the standard floral dip protocol with the Agl0 *Agrobacterium* strain^60^. After the floral dip procedure, seeds were harvested and plated on ½ MS plates containing 40 μg ml^−1^ hygromycin B, kept at 4 °C in the dark for 3 days to stratify, and then grown under a 16-h/8-h light/dark cycle at 23 °C. After 12 days, transgenic seedlings (except for transgenic seedlings targeting *PDS3* g2) were collected for genomic DNA extraction. The transgenic seedlings expressing Ymu1 targeting *PDS3* g2 were transplanted from the hygromycin B selection plates to soil and grown in a greenhouse (23 °C) for three weeks. After three weeks, leaf tissue was sampled for genomic DNA extraction by collecting and pooling three random leaves on an individual plant. Protoplast and transgenic plant tissue samples underwent genomic DNA extraction using Qiagen DNeasy plant mini kit (Qiagen, 69106). Amplicon sequencing was performed using the Illumina NovaSeqX platform. Analysis of amplicon sequencing was performed using the CrispRvariants R package (v.1.14.0) as previously described^8^. Primers used for amplicon sequencing can be found in Table S1.

##### Whole-genome off-target analysis

Three transgene-free *Arabidopsis* progeny lines harboring heritable edits generated by Ymu1-WFR at the *CHLI1* g4 and *PDS3* g12 sites were obtained from a previous study^16^. Two wild-type Col-0 replicates sequenced in a prior study^8^ were used as controls. Genomic DNA (50 ng) was sheared to approximately 300 bp using a Covaris S2 sonicator, and libraries were prepared using the Kapa Hyper DNA Kit (Roche, KK8502) following the manufacturer’s instructions. Sequencing achieved an average depth of approximately 3,737× (2,406× to 5,605×) coverage, with greater than 99% of the genome covered by mapped reads.

Whole-genome off-target analysis was performed following a previously established pipeline^8^ with minor modifications. Briefly, reads were mapped to the TAIR10 reference genome with BWA-MEM (v0.7.17)^47^, and duplicates were marked using GATK (v4.2.0.0) MarkDuplicates^43^. Base quality scores were then recalibrated with GATK BaseRecalibrator/ApplyBQSR, using a bootstrapped set of high-confidence variants (raw HaplotypeCaller calls passing the hard filters below) as known sites. SNPs and InDels were jointly called from the recalibrated alignments by GATK HaplotypeCaller and Strelka2 (v2.9.2)^44^, retaining only variants identified by both callers (intersection via BEDTools v2.30.0)^45^. Standard GATK hard filters were applied (SNPs: QD < 2.0, FS > 60.0, MQ < 40.0, SOR > 4.0; InDels: QD < 2.0, FS > 200.0, SOR > 10.0). Variants present in the wild-type control samples were subtracted, and remaining candidates with depth below 30× were excluded. All retained variants were manually inspected in BAM alignments to remove calling artifacts. Potential off-target sites were predicted using Cas-OFFinder (v3.0.0)^32^, allowing up to four mismatches across the TAM and guide RNA sequences (Tables S11-12).

##### TnpB-reRNA co-purification

Briefly, Escherichia coli BL21-AI cells (ThermoFisher Scientific, C607003) were co-transformed with the 10×His-MBP-TnpB expression plasmid and the corresponding T7-reRNA-HDV plasmid. Starter cultures were grown in 2XYT medium supplemented with ampicillin (0.1 mg/mL), chloramphenicol (0.034 mg/mL), and tetracycline (0.01 mg/mL) for ∼18 h at 37°C. A 5-mL aliquot of the starter culture was then used to inoculate 4.5 L of 2XYT expression medium containing the same antibiotics. Cultures were grown at 37°C until they reached an OD600 of 0.6-0.8, at which point TnpB expression was induced with 0.2% (w/v) L-arabinose. Induction proceeded overnight at 16°C for ∼16 h.

Cells were harvested by centrifugation and resuspended in Lysis Buffer (20 mM Tris-HCl, pH 8.0; 250 mM NaCl; 25 mM imidazole; 5% (v/v) glycerol; 1 mM TCEP) supplemented with cOmplete EDTA-free protease inhibitor (Roche). Cells were lysed by sonication and clarified by ultracentrifugation. The supernatant was applied to Ni-NTA resin (QIAGEN), washed five times with Wash Buffer (20 mM Tris-HCl, pH 8.0; 500 mM NaCl; 25 mM imidazole; 5% (v/v) glycerol; 1 mM TCEP) and eluted using 300 mM imidazole in the same buffer. Eluted protein was dialyzed overnight at 4°C in Dialysis Buffer (20 mM Tris-HCl, pH 8.0; 250 mM NaCl; 25 mM imidazole; 5% glycerol; 1 mM TCEP) in the presence of TEV protease.

Following TEV cleavage, the digest was passed over a HisTrap column (Cytiva) to separate the cleaved TnpB protein from the His-MBP tag. The flow-through containing tag-free TnpB was concentrated and further purified using a HiTrap Heparin HP affinity column (Cytiva), eluting the protein with Ion-Exchange Buffer (20 mM Tris-HCl, pH 8.0; X mM NaCl; 10% (v/v) glycerol; 1 mM TCEP), where a NaCl gradient from 250 mM to 1 M. The fraction was then concentrated and finally purified with size-exclusion chromatography on a Superdex 200 Increase 10/300 GL column (Cytiva) equilibrated in Gel Filtration Buffer (20 mM Tris-HCl, pH 8.0; 250 mM NaCl; 10% (v/v) glycerol; 1 mM TCEP). Purified TnpB was concentrated, aliquoted, snap-frozen in liquid nitrogen, and stored at -80°C. RNP concentrations were quantified by measuring A260 on a NanoDrop spectrophotometer using extinction coefficients calculated for each reRNA.

The homogeneity of proteins within purified RNPs was assessed by SDS-PAGE. Protein samples were mixed with 5× SDS Loading Dye (10% SDS; 30% glycerol; 75 mM EDTA; 250 mM Tris-HCl, pH 6.8; supplemented with bromophenol blue) at a 4:1 ratio and boiled at 95°C for 2 min. Samples were loaded onto Mini-PROTEAN TGX Precast Gels (Bio-Rad) and electrophoresed in a 1× SDS running buffer at 150 V for 45 min. Gels were stained using InstantBlue Coomassie Protein Stain (Abcam, ab119211) for ≥ 30 min, rinsed with MilliQ water, and imaged on a ChemiDoc MP system under the Coomassie Blue Gel 590/110 White Trans setting.

To assess the homogeneity of reRNA within purified RNPs, 3.5 μL of 2 μM RNP was prepared for three conditions: untreated, RNase-treated, and DNase-treated. RNase digestion was performed with DNase- and protease-free RNase A (ThermoFisher Scientific, EN0531), and DNase digestion was performed using RNase-free DNase I (New England Biolabs, M0303S) with 0.5 μL 10× DNase I Reaction Buffer (New England Biolabs, B0303S). Gel Filtration Buffer was added to each sample to a final volume of 5 μL. Reactions were incubated at 37°C for 10 min and quenched by adding an equal volume of colorless denaturing loading dye (94% formamide, 30 mM EDTA, 400 μg/mL heparin). Samples were resolved on a 15% denaturing urea-PAGE gel in 0.5× TBE for 1 h at 25 W. The gel was stained with SYBR Gold (ThermoFisher Scientific, S11494) for 30 min and imaged using a ChemiDoc MP imaging system (Bio-Rad) under the SYBR Gold 590/110 Blue Trans setting.

##### Oligonucleotide substrates for cleavage assays

All DNA oligonucleotides were synthesized by IDT and HPLC-purified. For the double-stranded DNA (dsDNA) substrates used in cleavage assays, the target strand (TS) was 5′-labeled with FAM and the non-target strand (NTS) was 5′-labeled with Cy5, unless otherwise specified.

To generate dsDNA substrates, equimolar amounts (typically 40 µM each) of complementary single-stranded oligonucleotides were mixed in DNA Annealing Buffer (10 mM Tris-HCl, pH 8.0; 100 mM NaCl; 1 mM EDTA), heated to 95 °C for 5 min, and then slowly cooled to 35 °C over 45 min on a thermocycler. Annealed products were quantified by measuring A260 on a NanoDrop spectrophotometer (ThermoFisher Scientific), and concentrations were calculated using manufacturer-provided extinction coefficients.

The annealing of dsDNA was confirmed by electrophoresis on an 8% native polyacrylamide gel run at 150 V at 4°C for 4 h. Working stocks were diluted to 200 nM and stored at -20 °C. The same procedure was used to prepare canonical and mismatched DNA substrates. Sequences of all DNA oligonucleotides used in this study are provided in Table S19.

##### Oligonucleotide cleavage assay

For DNA cleavage assays, TnpB-reRNA RNP complexes were prepared by incubating 100 nM RNP in Cleavage Buffer (1× condition: 10 mM Tris-HCl, pH 7.5; 100 mM NaCl; 2 mM MgCl₂; 1 mM EDTA; 1 mM DTT) at 26 °C for 15 min. Cleavage reactions were initiated by adding the pre-assembled RNP (76 μL) to annealed dsDNA substrate (4 μL) to yield a final concentration of 10 nM dsDNA and 100 nM TnpB RNP in 1× Cleavage Buffer. For competitive cleavage assays, salmon sperm DNA (Fisher Scientific, AM9680) was added to the reaction at a 30-fold mass excess relative to the dsDNA substrate prior to initiation (final concentration: 12 ng/μL salmon sperm DNA to 10 nM (0.4 ng/μL) dsDNA substrate). For substrate-excess assays, the concentrations were adjusted to 100 nM dsDNA and 20 nM TnpB RNP.

Reactions were quenched at defined time points (0.5, 1, 5, 10, 30, 60, and 120 min for standard enzyme-excess assays; with additional time points at 180, 240, and 1140 min for substrate-excess assays) by mixing an equal volume of 2× Quench Buffer (94% formamide; 30 mM EDTA; 400 μg/mL heparin). Samples were resolved on 15% denaturing urea-PAGE gels in 0.5× TBE at 40 W for 1 h. Gels were imaged using a Amersham Typhoon scanner (Cytiva), detecting FAM at 488 nm with a Cy2 emission filter (525BP20) or detecting Cy5 at 635 nm with a Cy5 emission filter (670BP30).

Band intensities were quantified using Bio-Rad ImageLab 6.1. Cleavage extent was calculated as the fraction of cleaved product relative to total lane signal. For all datasets, cleavage kinetics were fitted in GraphPad Prism using a mono-exponential model:

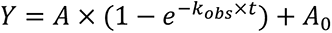

where Y is the fraction cleaved, A is the reaction amplitude, *k*_obs_ is the apparent rate constant, t is time (min), and A_0_ is a baseline offset. The *k*_obs_ values are reported as mean ± SD from independent mono-exponential fits to each reaction time course (n=3).

##### Plasmid cleavage assay

For plasmid cleavage assays, negatively supercoiled pHS697 (∼2.2 kb) was purified from DH5α cells (Fisher Scientific, 18-265-017) using the QIAprep Spin Miniprep Kit (QIAGEN, 27106). Supercoiling was confirmed by agarose gel electrophoresis. To prepare the linearized substrate, supercoiled pHS697 was digested with BsaAI using CutSmart buffer at 37°C (New England Biolabs, R0531) and purified using the QIAquick Gel Extraction Kit (QIAGEN, 28706). TnpB-reRNA RNP complexes were prepared by incubating 100 nM RNP in Cleavage Buffer (10 mM Tris-HCl, pH 7.5; 100 mM NaCl; 2 mM MgCl_2_; 1 mM EDTA; 1 mM DTT) at 26 °C for 15 min. Cleavage reactions were initiated by adding pre-assembled RNP (66 μL) to the plasmid substrate (16.5 μL) to yield final concentrations of 100 nM TnpB RNP and 3 nM plasmid in 1× Cleavage Buffer. Reactions were quenched at defined time points (linearized: 0.5, 1, 5, 10, 30, 60, 120, 180 min; supercoiled: 0.25, 0.5, 1, 2, 5, 10, 30, 120 min) by adding the samples to an equivalent volume of 2× Quenching Buffer (0.2% SDS, 50 mM EDTA). ⅕ the volume of 6× Orange DNA Loading Dye (Thermo Scientific) were added to the samples for gel visualization and products were resolved on 0.6% agarose gels in 1× TAE supplemented with 0.001% SYBR-SAFE DNA Gel Stain (Invitrogen) and imaged using a ChemiDoc MP Imaging System (Bio-Rad). Band intensities were quantified using ImageLab 6.1 (Bio-Rad) under the Blue Tray, SYBR Safe Gel (590/100) setting under Auto Optimal exposure and cleavage kinetics were fitted as described above.

##### Rotor bead tracking

###### Reagents

AuRBT studies were conducted with catalytically inactive Ymu1 TnpB RNP (dYmu1, dH4W, and dYmu1-WFR) with E279A mutation. DNA tethers for torque-driven and torsionally relaxed assays were assembled by ligating restriction enzyme digested PCR products as previously described^20,21,61^. Detailed information on the tether construction, including building blocks, can be found in Fig. S4A and Tables S20-22. Magnetic beads (ThermoFisher Scientific, Dynabeads MyOne Carboxylic Acid) were crosslinked with an antibody to Fluorescein/Oregon Green (ThermoFisher Scientific, A889) via EDC (ThermoFisher Scientific, 77149), as previously described^62^. These beads were prepared and stored at 4°C for up to two months.

###### Chambers

Flow chambers were constructed as previously described^19^, by sandwiching custom laser-cut Nescofilm channels between a hole-punched vinyl coverslip and glass coverslip spin coated with 0.1% nitrocellulose in isopentyl acetate (Ladd Research, 10800). On the day before experiments, streptavidin-coated gold nanospheres (20 μL original suspended volume per channel) with a nominal diameter of 80 nm (Cytodiagnostics, AC-80-04-15) were washed twice in Au wash buffer (50 mM Tris-HCl, pH 8.0; 0.05% TWEEN-20), centrifuging at 2000g for 5 minutes between washes. These were finally resuspended to their original suspended volume in blocking buffer (40 mM Tris-HCl, pH 8.0; 0.5 M NaCl; 0.2% TWEEN-20; 0.01% sodium azide; 5 mg/mL BSA). DNA tethers (∼2 pM) and magnetic beads (1.5 μL original suspended volume per channel) were added to the gold nanospheres in the blocking buffer and incubated overnight at 4°C on a rotator. Channels were also incubated overnight at 4°C with 12 μg/mL anti-digoxigenin in the PBS buffer. On the day of experiments, channels were (1) incubated with blocking buffer supplemented with 0.25% w/v casein for 1 h, (2) incubated with DNA tethers and both magnetic and rotor beads for 1 h, and (3) washed with approximately 15 channel volumes of C12T buffer (20 mM Tris-HCl, pH 8.0; 150 mM KCl; 5 mM MgCl_2_; 1 mM TCEP) supplemented with 0.2% Tween-20 (Sigma-Aldrich, P9416) and 0.2 mg/mL BSA (Invitrogen, 15561-020). All flow steps were controllably carried out with a syringe pump.

###### Microscopy

Experiments were conducted using a custom-built AuRBT microscope as previously described^21^. Briefly, this comprised a modified Nikon Eclipse Ti-S inverted microscope, with evanescent excitation provided by an intensity-stabilized 845-nm laser diode (Lumics LU0845M200) directly coupled to a polarization-maintaining fiber. A half-wave plate was used to achieve s-polarization at the sample interface. The return beam was collected on a position-sensitive detector to provide a signal for focus stabilization. Scattered light was collected with a Nikon Apo TIRF objective (60× /1.49 numerical aperture, oil) and imaged through an optical path splitter (Cairn, Optosplit III) onto a high-speed CMOS camera (Mikrotron, EoSens CL). Magnetic tweezers were implemented using a pair of 0.25 inch × 0.25 inch × 0.5 inch rectangular neodymium magnets (K&J Magnetics B448) mounted on a stage equipped with rotary and vertical servomotors (Physik Instrumente C-150.P and Physik Instrumente M-126.PD1). Samples were mounted on a three-axis nanopositioning stage (Mad City Labs PDQ-series).

###### Data Collection

Rotor bead tracking was done at 5 kHz, with DNA tethers subject to 5 pN of tension. To ensure proper attachments to the glass coverslip and magnetic bead, all DNA molecules were recorded for at least 5 minutes prior to data collection. Attachment type of the DNA molecule to the magnetic bead (free swivel or torsionally constrained) was further determined by rotating the magnets. For torque-driven assays, DNA twist was ramped between (+7.5, -5) by rotating the magnets at 3 RPM. Torque was calculated from the angular deflection of a transducer segment of torsional stiffness 0.26 pN nm/rad^20,63^. The zero-torque angular position of the magnets was calculated as the position of maximum extension of the DNA tether at low force^64,65^ and this zero was subtracted to obtain the reported twist values. A software delay between recorded and actual magnet angle was left uncorrected because of negligible effects on the results for the slow magnet rotation speed used in this study.

Prior to use in experiments, RNP was incubated at 100 nM in RBT buffer (1× condition: 10 mM Tris-HCl, pH 7.5; 10 mM MgCl_2_; 1 mM EDTA; 100 mM NaCl; 1 mM DTT) for 15 minutes at 37°C. Before data collection, we flowed in approximately 3 channel volumes of RNP in the RBT buffer supplemented with 0.2% Tween-20 (Sigma-Aldrich, P9416) and 0.2 mg/mL BSA (Invitrogen, 15561-020). RBT buffer was prepared from a 10× frozen stock for each experiment.

###### Torsionally Relaxed Assays

Angular traces of fluctuating rotor beads were analyzed by automated changepoint detection followed by merging of dwells for assigning *R*-loop states and scoring transition events^19^. Unlike previous work^19^, DNA tethers (Fig. S4A and Tables S20-22) had multiple modifications at each end for attachment to the surface and magnetic bead, but only tethers with free-swivel attachments to the magnetic bead (verified by cycling the magnets) were used for data collection. Cumulative rotor bead angles were directly converted to units of base pairs unwound assuming B-DNA helicity of 10.5 bp/turn.

Transitions between states were scored using the Steppi change-point analysis tool^22^, modeling the data as originating from an Ornstein-Uhlenbeck process. Global stiffness and coupling parameters were fixed by analyzing a portion of the trace before the introduction of any RNP. The only free model parameters were thus the mean angle of the rotor bead and the change point time. Adjacent dwells occupying the same state of *R*-loop formation (C, I, O) were subsequently merged as in prior work^15,19,20^. Boundaries between *R*-loop states were calculated using the arithmetic mean between lifetime-weighted average unwinding values of adjacent states. Traces were zeroed so that the predominant closed state corresponded to *Δθ*_%_=0 bp.

Transition rates between states were calculated by dividing the number of transitions from state i to state j by the total time spent in state i. Transitions were counted starting after the first transition event, typically C → I. Cartoons of free energy landscapes (Fig. 4J) were drawn with reference to the kinetic data shown in Table S3. Apparent equilibrium constants were computed using

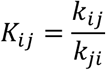

where *k_i,j_* represents the transition rate from state i to state j. Estimated free energy differences were then computed using

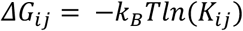

where *k_B_* represents the Boltzmann constant and T is temperature. Well positions for C, I, and O states were set to the lifetime-weighted average *Δθ*_0_ in each merged state cluster, and barrier heights relative to wells were represented as

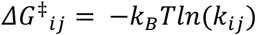

with barrier positions arbitrarily depicted halfway between free energy wells.

###### Torque-Driven Assays

For twist ramping experiments on torsionally constrained tethers, the acquired torque traces were analyzed to identify *R*-loop transitions and quantify transition rates as a function of imposed twist^15,20^. We expressed torque changes due to TnpB *R*-loop formation in units of base pairs unwound, assuming any change in equilibrium twist (*Δθ*_0_) of the tether is the result of bubble formation on B-DNA (10.5 bp per turn):

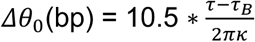

where *κ* is the torsional stiffness of the full DNA tether obtained from the slope of the torque-twist curve and *τ_B_*=*κθ* is the expected torque for unperturbed B-DNA.

*R*-loop states were identified using change point analysis as above. Twist-dependent transition rates, apparent equilibrium constants, and free energies were calculated^20^ using an approach adapted from an earlier procedure for analyzing force-dependent protein unfolding^66^. Briefly, the number of transitions between states i and j in each twist bin was counted and

normalized to obtain each rate k_ij_. Twist bins with fewer than 3 sampled transitions were excluded from analysis, and bins for positive twist ≥ 3 were omitted from kinetic and equilibrium fits and plots. Standard errors of transition rates were calculated by assuming Poisson statistics for transition events. Apparent equilibrium constants were then calculated as

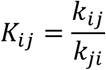

As described previously^15,20^, transitions between *R*-loop intermediates were modeled as transitions between states with changes in equilibrium twist *Δθ_ij_* and differences in free energy *ΔG_ij_* on a DNA polymer with torsional stiffness *κ*. Model parameters are derived from a linear fit of *ln*(*K_ij_*) as a function of imposed twist *θ*:

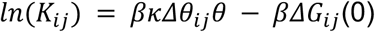

where 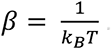. Estimated free energy differences were obtained as *ΔG*_&’_ = −*k*_(_*Tln*(*K_ij_*). Cartoon energy landscapes were drawn with reference to the above calculations, with barrier heights and well depths derived from linear fits to the transition rates and equilibrium constants, respectively, and the remainder of the energy landscapes drawn using arbitrary interpolant curves. Barrier locations were estimated by fitting a selected approximately linear portion for each *ln*(*k_ij_*) vs twist plot, assuming:

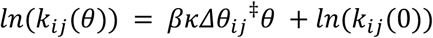

The *Δθ_ij_*^‡^ fit values were then scaled for display, so that the fractional positions of the displayed transition states correspond to *Δθ_ij_*^‡^ / (*Δθ_ij_*^‡^ − *Δθ_ji_*^‡^). Scaling was applied because fit values of *Δθ_ij_* did not reliably correspond to the physical differences in the *Δθ*_0_ values extracted from lifetime-weighted averaging, which were used to plot well positions.

##### Small RNA sequencing

Purified reRNA was isolated directly from assembled TnpB RNP complexes using the RNA Clean & Concentrator-5 Kit (Zymo Research) according to the manufacturer’s instructions. Approximately 20 ng of purified RNA was used for end-repair treatment with QuickCIP (New England Biolabs, M0525S) and T4 PNK (New England Biolabs, M0201S). RNA samples were incubated in a 20 µL reaction containing T4 PNK buffer (New England Biolabs, B0201S) and 1 µL QuickCIP at 37 °C for 30 min, followed by heat-inactivation at 85 °C for 10 min. Without additional cleanup, 1 µL T4 PNK was added directly to the same reaction and incubated at 37 °C for 30 min; ATP was then added to a final concentration of 5 mM and the mixture incubated for an additional 30 min at 37 °C. End-repaired RNA was purified again using the RNA Clean & Concentrator-5 Kit.

RNA-seq libraries were prepared using the Collibri™ Stranded RNA Library Prep Kit for Illumina Systems (Invitrogen, A38994024) following the manufacturer’s protocol. Purified RNA (10 µL) was combined with 10 µL of 2× Adapter Mix (BLUE), and adapter hybridization was performed in a thermocycler using a cooling ramp of −0.5 °C s⁻¹ with incubation at 65 °C for 10 min and 20 °C for 5 min. A ligation master mix (26.25 µL 2× Ligation Buffer (YELLOW) + 5.25 µL 10× Ligation Enzyme Mix) was added to the hybridization reaction for a total volume of 50 µL, followed by incubation at 20 °C for 15 min with the thermocycler lid off. Reverse transcription was performed by adding 50 µL of RT master mix (42 µL 2.5× RT Buffer (RED) + 10.5 µL 10× SuperScript™ IV Enzyme Mix) to the 50 µL ligation reaction and incubating at 50 °C for 10 min, followed by 85 °C for 5 min (lid at 90 °C). cDNA was purified using Dynabeads Cleanup Beads with two sequential bead cleanups and eluted in 20 µL Elution Buffer. Libraries were amplified using KAPA HiFi DNA Polymerase (Roche, KK2101) with an annealing temperature of 65 °C to minimize primer-dimer formation. Amplified libraries were size-selected by gel extraction on E-Gel™ EX 4% Agarose Gels (Invitrogen, G401004), quantified using the KAPA Library Quantification Kit (Roche, KK4873), and sequenced at the Innovative Genomics Institute Next-Generation Sequencing Core using an Illumina NextSeq 1000/2000 P2 v3 kit.

Paired-end reads were trimmed and merged with fastp (v0.23.2)^46^. Merged reads with lengths between 100 and 150 nucleotides were extracted using fastq-filter (v0.3.0) (https://github.com/LUMC/fastq-filter), then mapped to TnpB loci with BWA (v0.7.17)^47^. The resulting alignments were sorted with SAMTools (v1.17)^48^ and converted to BigWig format with deepTools2 (v3.5.1)^49^ for visualization in Integrative Genomics Viewer (IGV)^50^.

##### SHAPE-MaP RNA Structure Probing

SHAPE-MaP was performed as described previously^67^ with minor modifications. For each reaction, 1 µg purified RNP was mixed with 12 µL nuclease-free water and 6 µL 3.3× folding buffer (333 mM HEPES pH 8.0, 333 mM NaCl). Eight microliters of this mixture were combined with 1 µL 10× MgCl₂ (10 mM) and incubated at 37 °C. RNA was modified by adding 1 µL freshly prepared 100 mM 1M7 in DMSO or 1 µL DMSO, then brought to 100 µL and purified using the Qiagen RNeasy Micro Kit (QIAGEN, 74007), eluting in 14 µL water. Ten microliters of modified or control RNA were mixed with 0.8 µL random primer 9 (200 ng/µL) and 0.2 µL transcript-specific primer (2 µM), heated to 65 °C for 5 min, cooled on ice, and combined with 8 µL 2.5× MaP buffer (5× Pre-MaP buffer mixed 1:1 with 30 mM MnCl₂). After incubation at 25 °C for 2 min, 1 µL SuperScript II (ThermoFisher Scientific, 18-064-071) was added and reverse transcription was performed at 25 °C for 10 min, 42 °C for 3 h, and 70 °C for 15 min. cDNA was diluted to 68 µL and purified with MicroSpin G-25 columns (Cytiva, 27-5325-01). Double-stranded cDNA was generated with the NEBNext Second Strand Synthesis Module (New England Biolabs, E6111S) in an 80 µL reaction (68 µL cDNA, 8 µL buffer, 4 µL enzyme mix), incubated at 16 °C for 2.5 h, purified with a Zymo DNA Clean & Concentrator-5 kit, and eluted in 10 µL water. DNA was quantified by Qubit dsDNA HS (ThermoFisher Scientific, Q32851). Libraries were prepared with the Illumina Nextera XT kit. Tagmentation used 10 µL Tagment DNA buffer, 5 µL dsDNA (0.2 ng/µL), and 5 µL Amplicon Tagment Mix, incubated at 55 °C for 5 min and quenched with 5 µL NT buffer. Indexed PCR contained 25 µL tagmented DNA, 15 µL NPM, and 5 µL each index primer; cycling was 72 °C 3 min; 95 °C 30 s; 14 cycles of 95 °C 10 s, 55 °C 30 s, 72 °C 30 s; and 72 °C 5 min. Products were purified using AMPure XP beads (1.8×), washed twice with 80% ethanol, and eluted in 15 µL water. Libraries were checked on an Agilent TapeStation and molarity calculated with the Illumina Molarity Calculator and sequenced at the Innovative Genomics Institute Next-Generation Sequencing Core using an Illumina NextSeq 1000/2000 P1 v3 kit. Illumina reads were processed with ShapeMapper2 using: `shapemapper --name <RUN_NAME> --target <target.fasta> --modified --folder <MODIFIED_FASTQS_DIR> --untreated --folder <UNTREATED_FASTQS_DIR>` SHAPE reactivities were visualized in VARNA on the Ymu1 reRNA 2D structure.

##### Bacteria TAM assays

TAM depletion assays were performed in *E. coli* NEB 10-beta cells harboring a plasmid library in which all 8-nt TAM sequences are positioned immediately upstream of a single target site, following a modified version of the protocol described in a previous study^68^. For each Ymu1 variant (WT, pHS718; Ymu1-WFR, pHS751), 100 ng of the corresponding TnpB-expression plasmid was electroporated into 50 µL NEB 10-beta cells already containing the TAM library. Electroporation was performed at 1.7 kV using a time constant of 4.0-4.7 ms. Cells were recovered in NEB 10-beta/Stable Outgrowth Medium (New England Biolabs, B9035S) at 37 °C for 1 h. Following recovery, 50 µL and 100 µL aliquots were plated onto Nunc™ Square BioAssay Dishes (Thermo Scientific, 240845) containing LB agar supplemented with chloramphenicol, carbenicillin, and 2 nM anhydrotetracycline (Sigma-Aldrich, 37919-100MG-R) to induce TnpB expression. Plates were spread with sterile glass beads for 5-10 min and incubated at room temperature for 2 days to allow TAM depletion. Cells were harvested by adding 10 mL LB medium containing chloramphenicol and carbenicillin, resuspending the colonies, and recovering at 37 °C for 1 h.

For each library, 1 mL and 3 mL aliquots of the recovered culture were subjected to plasmid extraction using the QIAprep Spin Miniprep Kit (QIAGEN) and eluted in 50 µL nuclease-free water. The TAM region was amplified using Phusion™ High-Fidelity DNA Polymerase (New England Biolabs, M0530) and prepared for deep sequencing. Libraries were sequenced on a NovaSeq X Plus 1.5B flow cell (Illumina) with 150 bp paired-end reads at the UCLA Broad Stem Cell Research Center High-Throughput Sequencing Core.

Raw FASTQ files were quality-trimmed using Trim Galore v0.6.10 (Babraham Bioinformatics) with a minimum read length of 75 bp and a quality threshold of Q20. Paired-end reads were merged using PEAR v0.9.6^57^. Downstream analysis was performed in Python 3.8 using Biopython v1.85 for FASTQ parsing, pandas v1.4.0 and NumPy v1.22.1 for TAM extraction and quantification, and custom scripts to calculate log₂ fold-changes relative to both the naïve library and a non-targeting negative control. Motif logos were generated from position-weight matrices derived from the 20 most depleted TAM sequences using Logomaker v0.8.7 and matplotlib v3.5.1.

##### AF3 structural modeling

A SHAPE-informed AF3 model of the Ymu1 TnpB-reRNA-DNA complex was generated using the public AlphaFold Server (https://alphafoldserver.com)^53^. AF3 was run in multi-chain mode using the exact protein, RNA, and DNA sequences listed in Table S23. Specifically, the protein chain comprised full-length Ymu1 TnpB (382 aa). The RNA chain contained the 127-nt reRNA scaffold with a 16-nt guide base-paired to Target 1, including the TAM. To preserve experimentally supported RNA secondary features, we introduced nucleotide substitutions during AF3 runs that enforce known elements such as the pseudoknot (guided by SHAPE reactivity). The DNA duplex was represented by a 22-nt target strand spanning the TAM and protospacer and an 8-nt non- target strand containing only the TAM and immediately downstream bases, which prevents unintended TS-NTS annealing while preserving the local *R*-loop interface. Our AF3 modeling of Ymu1 TnpB consistently converges on a conformation in which the lid domain occludes the RuvC active site and positions the catalytic residues far from the scissile phosphates, matching the ternary state captured experimentally in Dra2 TnpB (ternary conformation 1)^2^. Because the detailed architecture of the distal reRNA scaffold remains uncertain and is not central to our mechanistic conclusions, we do not interpret or discuss its fine structural features. Accordingly, in all figures we display only the protein and DNA components of the AF3 model (omitting the reRNA scaffold) for clarity.

##### Protein mutagenesis

Residues were chosen based on structural proximity to nucleic acid in an AF3 model of the Ymu1 ternary complex and correspondence to mutationally sensitive positions identified by DMS of Dra2 TnpB^12^ where alignment was reliable. We prioritized three structural regions: the WED domain, bridge helix, and lid subdomain. At H4 (WED domain, adjacent to the first DNA-RNA base pair), hydrophobic aromatic residues (W, F, Y) were tested to probe stabilization of the initial base pair through stacking interactions, motivated by aromatic enrichment at the homologous Dra2 position^12^ and analogous engineering of Cas12f^69^. At K229 and R230 (bridge helix), charge-altering (A, E, Q) and hydrophobic (I, L, V) substitutions were tested to evaluate coupling between this helix and distal *R*-loop formation. At positions within the lid and lid-adjacent helix (V283, G285, M287, H290, A293, S303, L304, V305), substitutions were chosen to probe conformational dynamics. The lid undergoes rearrangement upon *R*-loop formation in Cas12a and related Cas12s^70–72^, and this region includes positions homologous to DMS-sensitive residues in Dra2 TnpB (Dra2 E302/Ymu1 S303; Dra2 I304/Ymu1 V305). At S303, nine substitutions spanning charge, polarity, and hydrophobicity were tested; at L304, three (F, K, R); at V305, two (F, R). Combinatorial variants were generated by pairing the nondeleterious single mutations (L304F-V305R, H4W-V305R, H4W-L304F, and H4W-L304F-V305R). In total, 54 single and combinatorial variants were screened (Fig. 2A).

##### Bioinformatic analysis

For generation of the tree used in Fig. 1A, a selection of TnpB and Cas12 genes spanning diverse types were selected from a prior study^6^ and CasPEDIA database^73^ and folded using ColabFold2^54^ on default settings. The dendrogram was generated using DaliLite.v5^55^ with the --matrix flag.

For additional phylogenetic analyses of the three substitutions (Fig. S6 and Note S2), TnpB homologous sequences were first identified using PSI-BLAST via the MPI Bioinformatics Toolkit (https://toolkit.tuebingen.mpg.de), searching against the NR70 databases. PSI-BLAST was run for 8 iterations using the BLOSUM45 scoring matrix, with E-value cutoffs of 1×10^-3^ for both reporting and inclusion in subsequent iterations, and a maximum of 10,000 target hits. The resulting multiple sequence alignments were used directly for downstream analysis. Each MSA was filtered to remove sequences containing gaps at any of the analyzed positions. Residue frequencies at each column were computed by tallying amino acid symbols and dividing by the total number of non-gap residues (i.e., the number of sequences in the filtered alignment). To identify residues that co-evolve and are likely to mediate interactions across structural interfaces in TnpB, we trained a Potts model on the TnpB multiple sequence alignment (MSA). Specifically, we used the TnpB MSA derived from the NR70 dataset (as described above) and removed all alignment columns containing gaps in the Ymu1 TnpB reference sequence. We then analyzed this MSA with the GREMLIN Colab notebook^56^ and obtained average product-corrected (APC) coupling matrices. In these matrices, we examined positions 4, 304 and 305 corresponding to Ymu1 residues H4, L304 and V305, and selected the top five positions with APC scores > 5 standard deviations above the mean. H4 was weakly coupled to Q3, L150, indicating low significance of these potential contacts for the overall structure across TnpB homologs. Two clusters of strongly coupled residues: (1) D201, N202, L262, S266, L308 for L304, and (2) D196, I278, D280, Q309, R321 for V305, were mapped onto WT Ymu1 and Ymu1-WFR AF3 structures (Fig. S6H, S6J), and the homologous residues were analyzed in the ESM-IF-generated TnpB variant experimental structures (Fig. S6K, S6L).

## QUANTIFICATION AND STATISTICAL ANALYSIS

Statistical details for all experiments are reported in the corresponding figure legends and supplemental tables. No statistical methods were used to predetermine sample size. Data are presented as mean ± SEM for plant genome-editing experiments (biological replicates) and mean ± SD for biochemical cleavage assays (independent reactions).

For plant genome-editing experiments (Fig. 1F, 2A, 2C, 2D), n refers to the number of independent biological replicates (individual plants or protoplast transfections), as indicated in each figure legend. Statistical significance for comparisons between WT and mutant editing efficiencies (Fig. 2C, 2D) was assessed using unpaired two-tailed Student’s t-tests, with exact p-values reported in the figures.

For biochemical cleavage assays (Figs. 3, 5G-H, 5J, and S3-S5), n = 3 independent reactions were performed for each condition. Apparent rate constants (*k*_obs_) were obtained by fitting each individual reaction time course to a mono-exponential model in GraphPad Prism (see Method Details), and values are reported as mean ± SD from three independent fits. For mismatch cleavage assays, *k*_obs_ values were not determined for conditions where total cleavage remained below 10% at the final time point (Table S10).

For single-molecule AuRBT measurements (Figs. 4, 5A-F, and S4-S5), n refers to the number of independent DNA tethers analyzed. Complete data collection statistics are given in tables S4 and S7, including the number of unique tethers and channels for each experimental condition and the number of scored events for each transition type. Reported errors for transition rates (Tables S3 and S5) were calculated assuming Poisson statistics of state transitions, and resulting errors for derived equilibrium constants and free energy differences were calculated using error propagation. Linear fits to ln(*K*) and ln(*k*) versus imposed twist (Fig. 5E and Fig. S5F-G) were performed by least-squares regression; fit parameters are summarized in Table S6.

All data analysis and curve fitting were performed using GraphPad Prism, MATLAB (R2019a), and custom Python scripts. No data points were excluded from analyses, apart from exclusions noted above in the analysis of torque-driven assays.

