## Supplemental Document for "Stepwise DNA unwinding gates TnpB genome-editing activity"

### Supplemental Figures

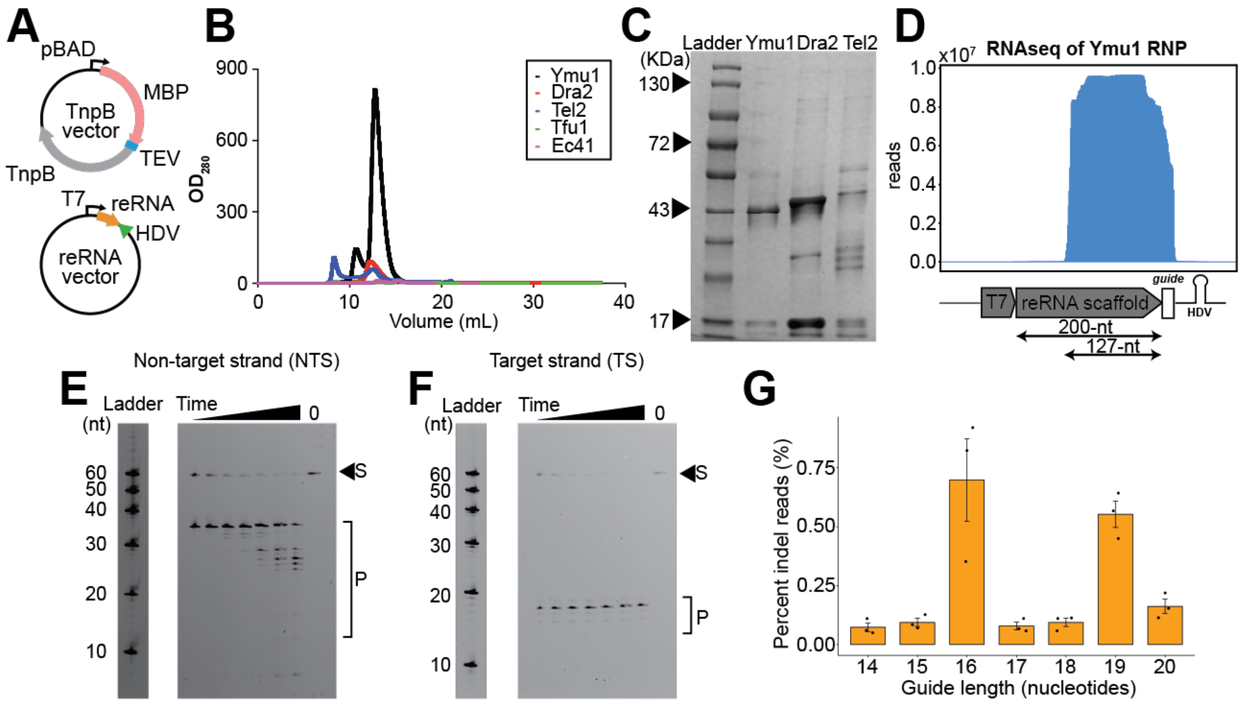

**Fig. S1. Additional characterization of Ymu1 TnpB and guide configuration (related to Fig. 1). (A)** Schematic of the bacterial expression system used for TnpB ribonucleoprotein (RNP) production. TnpB protein is expressed from an arabinose-inducible promoter, while the reRNA is expressed from a separate plasmid under a T7 promoter and processed by a downstream HDV ribozyme. **(B)** Size-exclusion chromatography traces of five TnpB orthologs (Ymu1, Dra2, Tel2, Tfu1, Ec41). Absorbance at 280 nm (y-axis) is plotted against elution volume (x-axis, mL). **(C)** SDS-PAGE analysis of purified TnpB orthologs corresponding to panel A, confirming protein homogeneity following affinity and size-exclusion purification. **(D)** RNA-seq analysis of Ymu1 reRNA expressed in E. coli from a T7 promoter using a 200-nt reRNA scaffold. **(E-F)** Representative denaturing gels of Target 1 cleavage by reconstituted Ymu1 RNP, showing **(E)** non-target strand (NTS) and **(F)** target strand (TS) cleavage. The uncleaved substrate (S) and cleavage product (P) are indicated. **(G)** Guide-length screen in *Arabidopsis* protoplasts using a single-transcript TnpB-reRNA-HDV design with the WT TnpB and 127-nt reRNA. The percentage of indel reads (y-axis) at the endogenous PDS3 g2 target site using guide length ranging from 14 to 20 nts (x-axis). Bars represent mean ± SEM (n = 3 biological replicates).

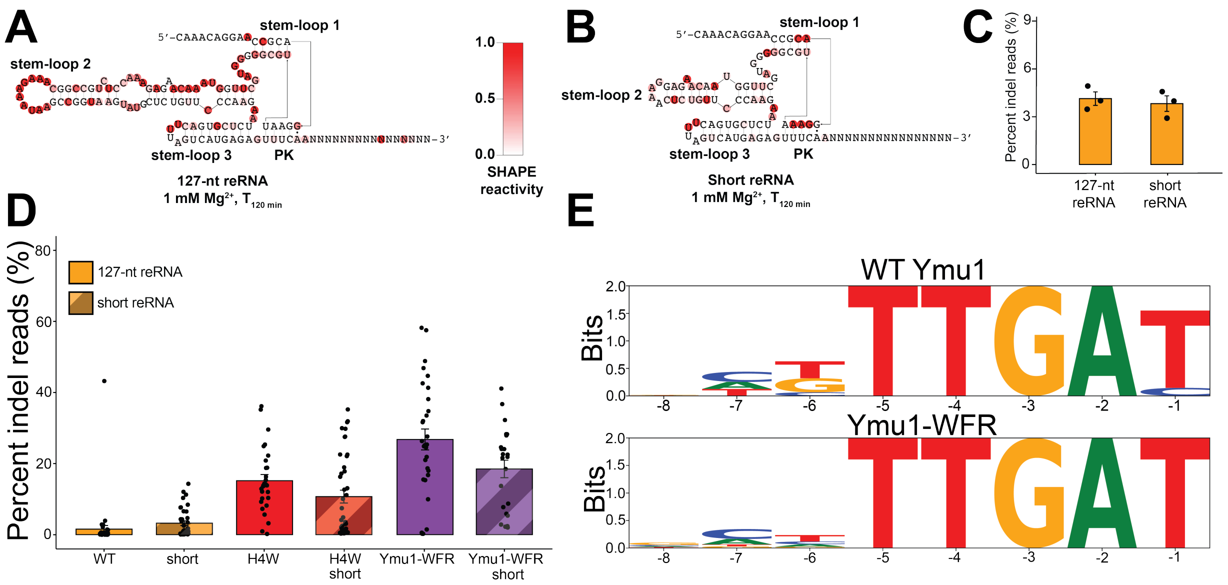

**Fig. S2. Structural and functional analysis of Ymu1 reRNA and scaffold variants (related to Fig. 2).** **(A-B)** SHAPE-MaP reactivity profile for the **(A)** 127-nt reRNA and **(B)** short reRNA scaffold. Reactivities are mapped onto the predicted secondary structure, with major stem-loop elements and pseudoknot (PK) indicated. **(C)** Protoplast editing efficiencies comparing the 127-nt and short reRNA scaffolds using WT Ymu1. Percentage of indel reads (y-axis) obtained from amplicon sequencing the *Arabidopsis* PDS3 g2 site (x-axis). Bars represent mean ± SEM (n = 3 biological replicates). **(D)** Comparison of T1 whole-plant editing efficiencies targeting *Arabidopsis* PDS3 g2 using full-length versus short reRNA scaffolds. Bar plot shows percentage of indel reads (mean ± SEM). **(E)** Bacterial TAM-discovery assay for WT Ymu1 and Ymu1-WFR. Sequence logos summarize nucleotide enrichment adjacent to the target site.

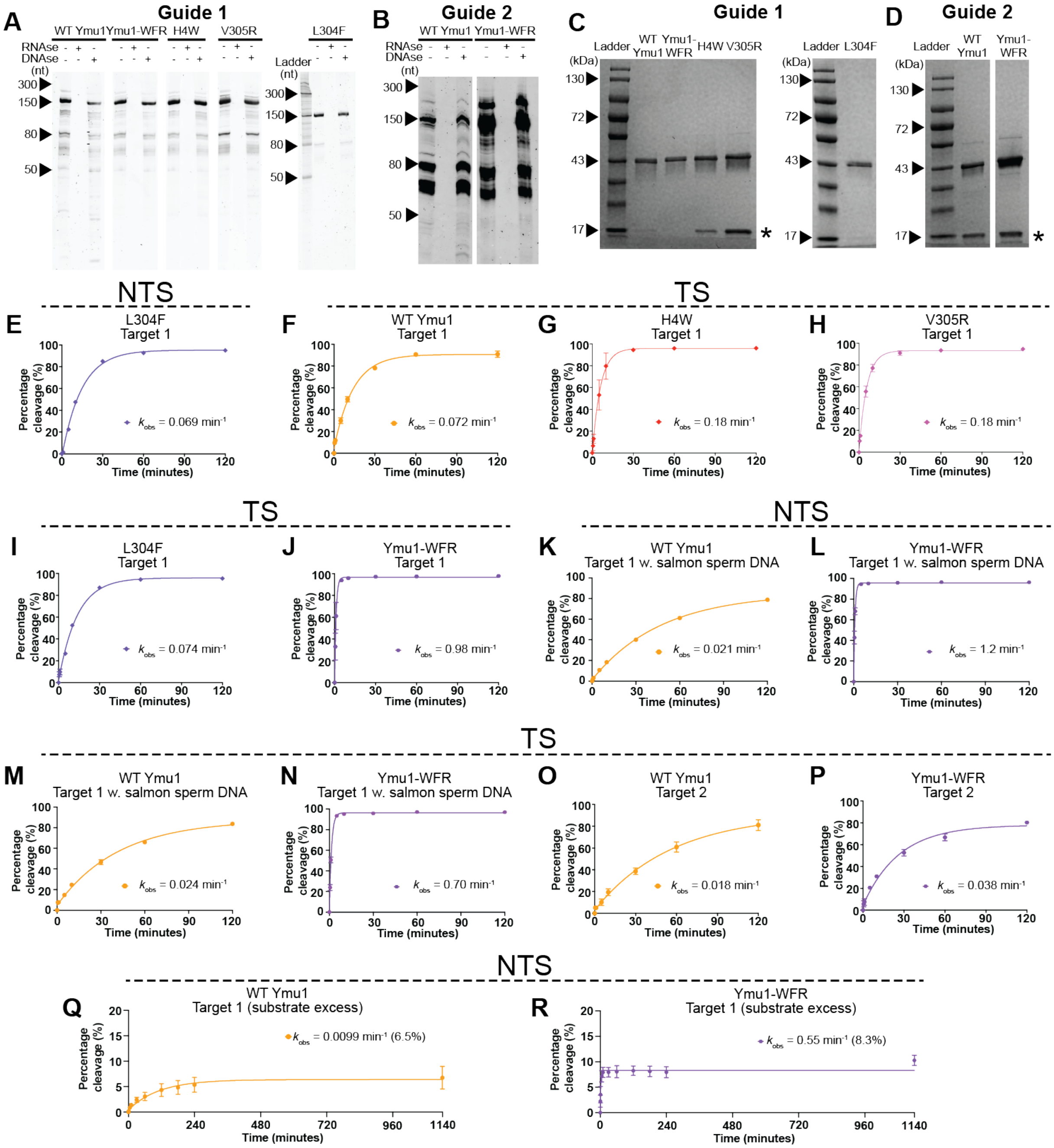

**Fig. S3. Additional cleavage assays and biochemical quality controls (related to Fig. 3) (A-B)** Denaturing PAGE analysis of the reRNA homogeneity for RNP samples used in cleavage assays targeting **(A)** Target 1 and **(B)** Target 2. Gels are shown under untreated, RNase-treated, and DNase-treated conditions. **(C-D)** SDS-PAGE analysis of the TnpB protein homogeneity for RNP samples used in cleavage assays targeting **(C)** Target 1 and **(D)** Target 2. The asterisk (*) indicates a ~17 kDa minor band identified by mass spectrometry as an N-terminal REC lobe fragment. The percentage of full-length (~44 kDa) protein estimated by densitometry is: **(C)** WT 84.0%, H4W 73.2%, V305R 61.0%, L304F 82.3%, Ymu1-WFR 82.8% (Target 1); **(D)** WT 64.4%, Ymu1-WFR 69.8% (Target 2). **(E)** NTS cleavage profiles for L304F (Target 1). **(F-L)** TS cleavage profiles for Ymu1 TnpB variants. Percentage of cleaved substrate (y-axis) is plotted over time (x-axis). Each point represents mean ± SD (n = 3 independent reactions); $k_{obs}$ values reflect the mean from three independent mono-exponential fits on each reaction time course (n=3). **(F)** WT Ymu1 (Target 1). **(G)** H4W (Target 1). **(H)** V305R (Target 1). **(I)** L304F (Target 1). **(J)** Ymu1-WFR (Target 1). **(K-N)** NTS and TS cleavage profiles in the presence of 30-fold mass excess salmon sperm competitor DNA (Methods). **(K)** WT Ymu1 NTS (Target 1). **(L)** Ymu1-WFR NTS (Target 1). **(M)** WT Ymu1 TS (Target 1). **(N)** Ymu1-WFR TS (Target 1). **(O-P)** TS cleavage profiles for **(O)** WT Ymu1 (Target 2) and **(P)** Ymu1-WFR (Target 2). **(Q-R)** Substrate-excess cleavage assays (100 nM dsDNA, 20 nM RNP) for **(Q)** WT Ymu1 and **(R)** Ymu1-WFR on Target 1. Mono-exponential fits modeling single-turnover behavior are shown, with fitted amplitudes (%) in parentheses. $k_{obs}$ values (mean ± SD) are also provided in Table S2.

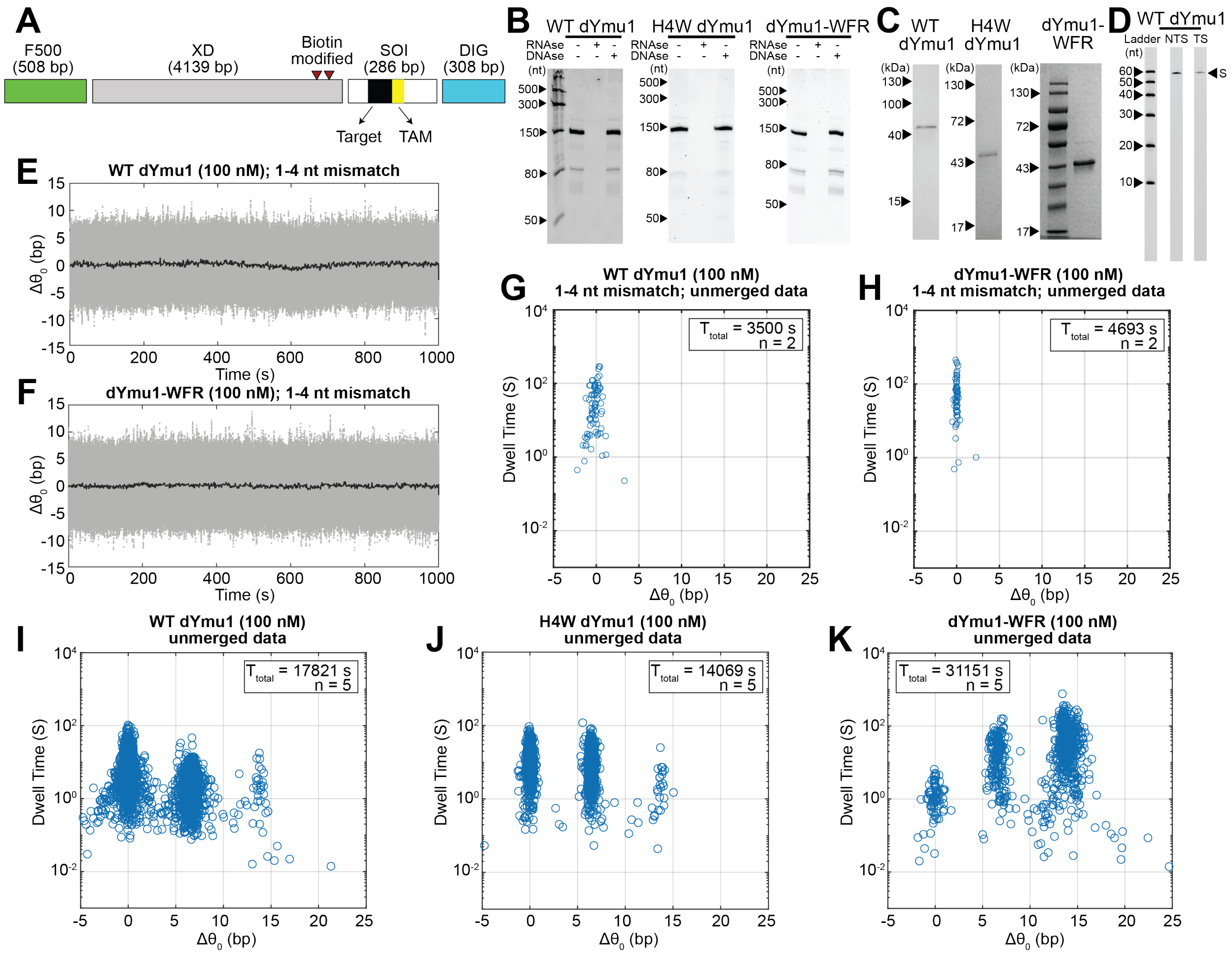

**Fig. S4. Controls and validation for equilibrium AuRBT experiments (related to Fig. 4).**

**(A)** Schematic of DNA tether used in AuRBT. Tethers were assembled by ligating four PCR-generated fragments (Table S20). The F500 and DIG segments contained fluorescein- and digoxigenin-modified dUTPs for bead and coverslip attachment, respectively. The transducer segment (XD) carried two biotin modifications for rotor-bead linkage. The sequence-of-interest (SOI) included a 5′-TTGAT-3′ TAM followed by the Target 1 sequence for dYmu1 TnpB binding (Table S21). The full tether sequence is provided in Table S22. **(B)** Denaturing PAGE analysis of the reRNA homogeneity for WT dYmu1, H4W dYmu1, and dYmu1-WFR TnpB samples used in torsionally-relaxed equilibrium AuRBT experiments. Gels are shown under untreated, RNase-treated, and DNase-treated conditions. **(C)** SDS-PAGE analysis of the TnpB protein homogeneity for WT dYmu1, H4W dYmu1, and dYmu1-WFR samples used in torsionally-relaxed equilibrium AuRBT experiments. **(D)** Representative denaturing gel of Target 1 oligonucleotide dsDNA cleavage by reconstituted WT dYmu1 (E279A) RNP in AuRBT buffer conditions showing no detectable cleavage after 120 min. **(E-F)** Representative trajectories of $\Delta\theta_0$ (bp) over time (s) using tethers containing DNA-RNA mismatches introduced into nt 1-4 of the target DNA sequence. **(E)** WT dYmu1 and **(F)** dYmu1-WFR. Low-pass-filtered traces (1 Hz) are shown in black. **(G-H)** Scatter plots of dwell times (s) versus $\Delta\theta_0$ (bp) for all unmerged Steppi-assigned dwells for experiments with 1-4 nt

DNA-RNA mismatches. **(G)** WT dYmu1 and **(H)** dYmu1-WFR. The total collection time ($T_{total}$) and the number of DNA tethers analyzed (n) are reported in the figure legend. *R*-loop formation events are not observed, consistent with a requirement for initial seed matching. **(I-K)** Scatter plots of state dwell times (s) versus $\Delta\theta_0$ (bp) for all unmerged Steppi-assigned states of **(I)** WT dYmu1, **(J)** H4W dYmu1, and **(K)** dYmu1-WFR. The total collection time ($T_{total}$) and the number of DNA tethers analyzed (n) are reported in the figure legend. Complete AuRBT trace statistics, including the numbers of detected transitions, are summarized in Table S4.

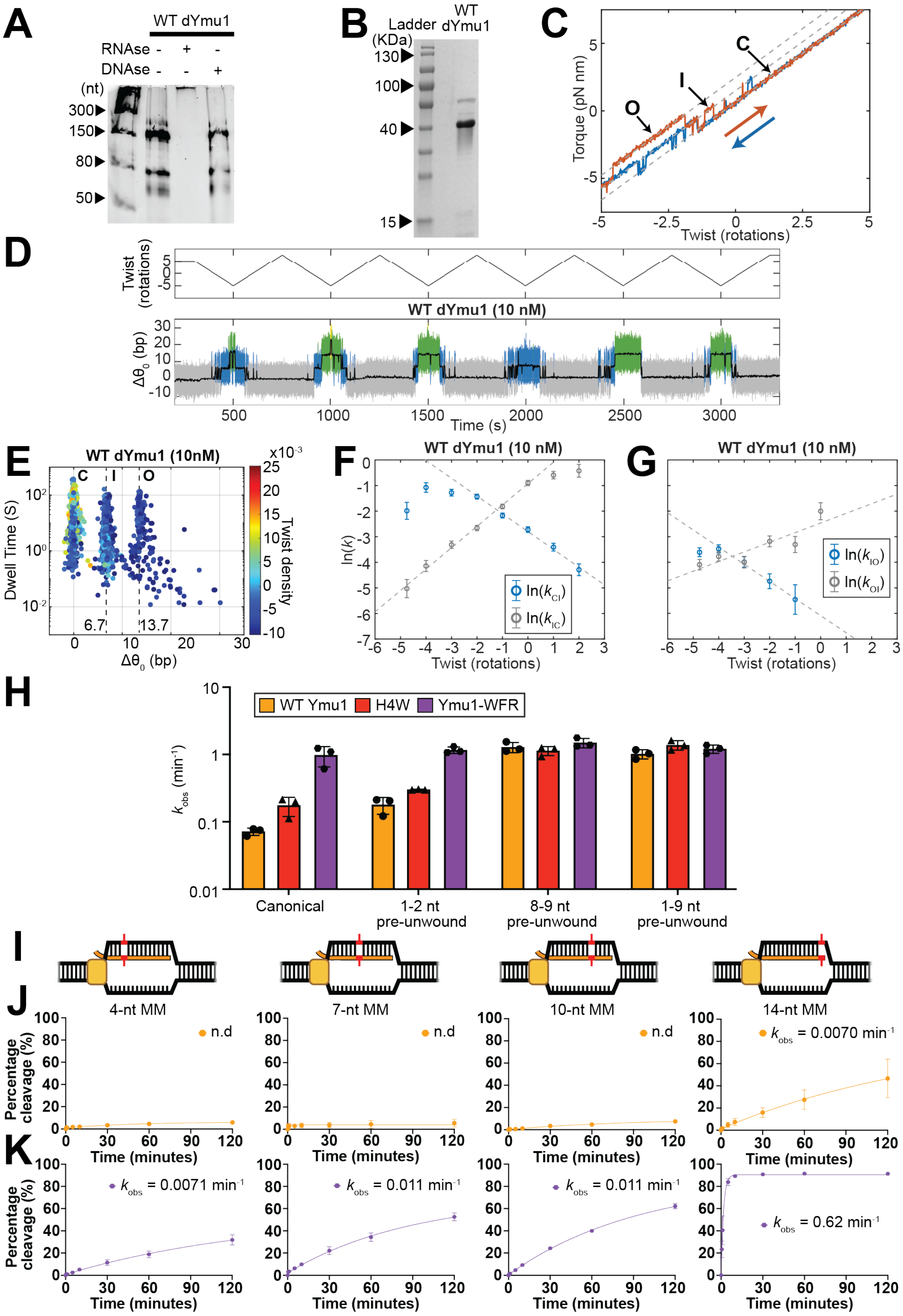

A
WT dYmu1
RNAse - + -
DNAse - - +
(nt)
300
150
80
50
B
Ladder
(KDa)
WT dYmu1
130
100
40
15
C
Torque (pN nm)
Twist (rotations)
O
I
C
D
Twist (rotations)
WT dYmu1 (10 nM)
Δθ0 (bp)
Time (s)
E
WT dYmu1 (10nM)
Dwell Time (S)
Δθ0 (bp)
Twist density
x10-3
6.7
13.7
F
WT dYmu1 (10 nM)
ln(k)
ln(kCI)
ln(kIC)
Twist (rotations)
G
WT dYmu1 (10 nM)
ln(kIO)
ln(kOI)
Twist (rotations)
H
WT Ymu1
H4W
Ymu1-WFR
kobs (min-1)
Canonical
1-2 nt pre-unwound
8-9 nt pre-unwound
1-9 nt pre-unwound
I
J
4-nt MM
7-nt MM
10-nt MM
14-nt MM
Percentage cleavage (%)
Time (minutes)
n.d
kobs = 0.0070 min-1
K
kobs = 0.0071 min-1
kobs = 0.011 min-1
kobs = 0.011 min-1
kobs = 0.62 min-1

**Fig. S5. Additional analyses for DNA unwinding, supercoiling, and mismatch cleavage experiments (related to Fig. 5) (A)** Denaturing PAGE analysis of the reRNA homogeneity for the WT dYmu1 sample used in torque-driven AuRBT experiments. RNA gels are shown under untreated, RNase-treated, and DNase-treated conditions. **(B)** SDS-PAGE of purified dYmu1 TnpB protein used in the torque-driven AuRBT experiments. The WT dYmu1 RNP sample in Fig. 5 and Fig. S5 was prepared independently from that used in equilibrium AuRBT assays. **(C)** Torque vs. twist for a twist ramping cycle in the presence of WT dYmu1. Blue (unwinding) and orange (rewinding) arrows indicate the direction of ramping. **(D)** Representative twist ramping trace (a portion of this trace is shown in Fig. 5B). (Top) Imposed twist (rotations). (Bottom) $\Delta\theta_0$ (expressed in units of base-pairs unwound assuming changes arise from local DNA unwinding of B-DNA). Transitions between distinct states are scored using automated change-point detection followed by merging to prevent overscoring[S1,2]. States are color-coded as closed (C, gray), intermediate (I, blue), open (O, green), and extended open (O′, yellow). Low-pass-filtered traces (1 Hz) are shown in black. **(E)** Scatter plot of state dwell times (s) (y-axis) versus $\Delta\theta_0$ (bp) (x-axis), colored by twist density at state onset. Twist density was calculated as imposed twist (in turns) divided by the relaxed linking number of the DNA tether ($Lk_0$=N/10.5 where N is the length of the tether in bp). **(F-G)** Plots of ln(*k*) versus imposed twist (rot), for forward and reverse transition rates $k_{ij}$ in $s^{-1}$ of **(F)** C↔I and **(G)** I↔O transitions for WT dYmu1. Data points are listed in Table S5 and linear fit parameters are summarized in Table S6. AuRBT trace statistics, including the number of tethers analyzed, total tracking time, and number of detected transitions, are summarized in Table S7. **(H)** Pre-unwound DNA cleavage assays showing apparent cleavage rates ($k_{obs}$) as a function of initial duplex opening of WT, H4W, and Ymu1-WFR. Shown are TS cleavage. NTS cleavage profiles are provided in Fig. 5J. **(I)** Schematic of dsDNA substrates containing single nucleotide RNA-DNA mismatches at positions 4, 7, 10, and 14 within the guide-target heteroduplex. **(J-K)** NTS cleavage profiles for WT Ymu1 and Ymu1-WFR on mismatched substrates. **(J)** WT Ymu1 and **(K)** Ymu1-WFR. Percentage of cleaved substrate (y-axis) is plotted over time (x-axis). Each point represents mean ± SD (n = 3 independent reactions); $k_{obs}$ values reflect the mean from three independent mono-exponential fits on each reaction time course (n=3). n.d., not determined due to low cleavage levels that precluded reliable fitting. $k_{obs}$ values (mean ± SD) for both NTS and TS cleavage are provided in Table S10.

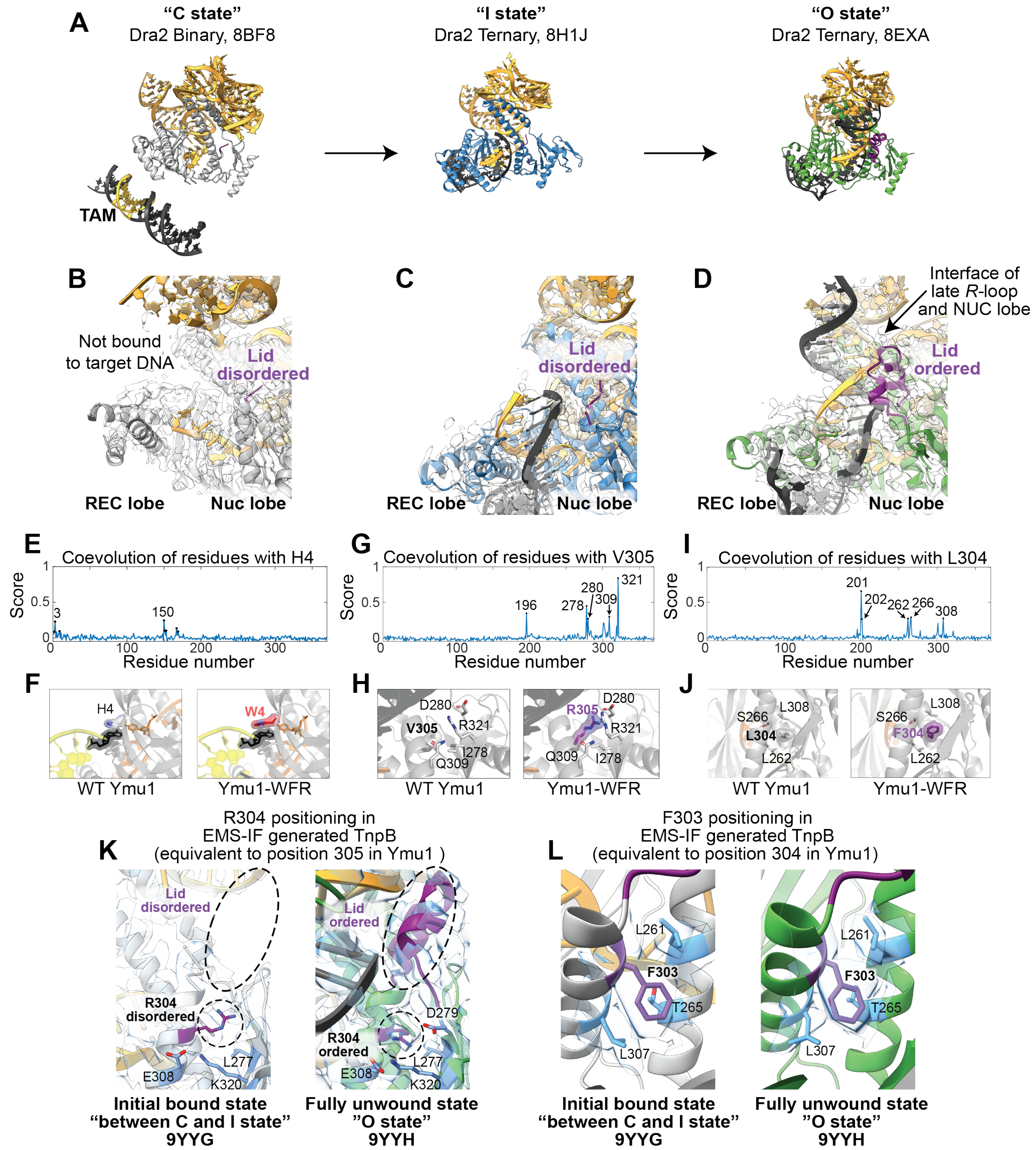

**Fig. S6. Analyses supporting mutational effects at positions 4, 305, and 304 (related to Fig. 6).**
**(A)** Comparison of cryo-EM structures of Dra2 TnpB in distinct conformational states. The closed ("C") state is defined by intact duplex DNA and corresponds to a DNA-free or binary-like protein complex conformation in solution. The Dra2 binary complex lacking target DNA[S3] (PDB: 8BF8) is therefore shown as a reference for the protein conformation associated with the C state. The ternary state reported by Nakagawa et al.[S4] (PDB: 8H1J) is proposed to correspond to the intermediate ("I") state, and the ternary state reported by Sasnauskas et al.[S3] (PDB: 8EXA) is proposed to correspond to the fully-formed open

(“O”) state. **(B-D)** Experimentally resolved heteroduplex and lid domain structures across conformational states of Dra2 TnpB. **(B)** In 8BF8 (“C state”), the lid domain is disordered and DNA is not unwound. **(C)** In 8H1J (“I state”), the lid remains mostly unstructured, possibly allowing transient interactions with a partially formed DNA heteroduplex, which mostly binds the REC lobe. 6-8 bp of heteroduplex are most strongly resolved in the density; 8 bp are shown in the figure, fewer than the 12 bp included in the 8H1J model[S4]. **(D)** In 8EXA (“O state”), the lid domain becomes ordered and facilitates binding of TnpB with the RNA-DNA heteroduplex along its whole length, consistent with the fully unwound O state. The heteroduplex is stabilized by both the REC and the NUC lobes. **(E)** Co-evolution profile for residue 4 with other protein residues across ~4,500 non-redundant TnpB homologs. **(F)** AF3 ternary-complex model of Ymu1 TnpB highlighting residue 4 in the WED domain. The H4W substitution is positioned adjacent to the first RNA-DNA base pair of the heteroduplex. **(G)** Co-evolution profile for residue 305. **(H)** AF3 ternary-complex model highlighting residue 305 within the lid-adjacent helix, proximal to a network of residues in the catalytic RuvC domain. **(I)** Co-evolution profile for residue 304. **(J)** AF3 ternary-complex model highlighting residue 304, located within a hydrophobic interface between the lid-adjacent and bridge helices. **(K-L)** Local structural environments of residues in cryo-EM structures of an ESM inverse-folding (ESM-IF) generated TnpB[S5] homologous to the Ymu1 WFR substitutions. **(K)** Residue R304 in the ESM-IF-generated TnpB (homologous to Ymu1 V305R substitution) does not form cryo-EM density-supported contacts in the initial bound state (PDB: 9YYG), whereas it is associated with a strong density in the fully unwound state (PDB: 9YYH). **(L)** Residue F303 in the ESM-IF-generated TnpB (homologous to Ymu1 substitution L304F) is structured in both early and late *R*-loop states. For each panel, the left view shows an initial bound state preceding the *R*-loop formation (PDB: 9YYG), and the right view shows the fully unwound state (PDB: 9YYH). Density and model views illustrate state-dependent ordering of the lid region and associated residues. Cryo-EM maps in **(B-D)** were contoured at 7.0, 0.5, 8.0, respectively, whereas **(K-L)** were contoured at 0.1 (dataset-dependent map units).

### Supplemental Notes

#### Note S1. Kinetic and equilibrium analysis of TnpB cleavage and binding

To relate the bulk cleavage rate ($k_{obs}$) to the stepwise transition rates measured by AuRBT, we consider the overall reaction in which RNP binding and *R*-loop initiation is followed by R-loop propagation and finally strand scission. We assume the following simple model:

$$RNP + DNA_C \underset{k'_{CI}}{\overset{k_{IC}}{\leftrightharpoons}} DNA_I \underset{k_{IO}}{\overset{k_{OI}}{\leftrightharpoons}} DNA_O \xrightarrow{k_{cleave}} DNA_{cleaved}$$

Here, $k'_{CI}$ is a bimolecular rate constant for binding and *R*-loop initiation, approximated as a single kinetic step and related to the measured rate $k_{CI}$ by $k'_{CI} = \frac{k_{CI}}{[RNP]}$. $k_{cleave}$ is the effective rate of completing DNA scission from the O state. All other constants are as defined in Table S3.

Under the standard conditions of our bulk cleavage assay (Fig. 3 and Fig. S3; Methods), a single-turnover reaction is initiated by mixing, and the assay monitors the accumulation of $DNA_{cleaved}$. The time course is well approximated by a mono-exponential model, and the fit time constant $k_{obs}$ represents a close estimate of the inverse of the mean first passage time for the complete reaction pathway, given by:

$$k_{obs} \sim \tau^{-1}{}_{mfp} = \frac{[RNP]*(\frac{k_{IO}k_{cleave}}{k_{OI}+k_{cleave}+k_{IO}})}{(\frac{k_{IC}}{k_{CI}}*\frac{k_{OI}}{k_{cleave}}+\frac{k_{IC}}{k_{CI}}+\frac{k_{IO}}{k_{CI}})*(\frac{k_{cleave}}{k_{OI}+k_{cleave}+k_{IO}})+[RNP]}$$

By assuming that all rates apart from $k_{cleave}$ are the same as measured in single-molecule measurements of dYmu1 (Fig. 4, Table S3), we estimate $k_{cleave} \approx 0.45\ s^{-1}$ by setting $\tau^{-1}{}_{mfp,WT} = 0.0012\ s^{-1}$ equal to the measured $k_{obs,WT}$ in biochemical assays with 100 nM WT RNP. Assuming that $k_{cleave}$ is unchanged in WFR, this predicts $\tau^{-1}{}_{mfp,WFR} = 0.012\ s^{-1}$. This simple model is sufficient to predict a ~10-fold acceleration for WFR, with an overall cleavage rate that is below but on the same order as the measured $k_{obs,WFR} = 0.033\ s^{-1}$. Note that for the [RNP] used in the assay, WFR is predicted to have an overall rate that is close to saturation as a function of [RNP], and is principally limited by the slow I→O transition rate $k_{IO}$**.** In the limit of low [RNP] where neither WT nor WFR is at saturation, the kinetic difference is predicted to be larger, approaching ~180-fold rate enhancement of WFR over WT.

The single-molecule measurements also provide information that directly relates to the dissociation constants of *R*-loop complexes for the dYmu1 variants used in this study. In particular, the measurements imply much tighter on-target binding for WFR than for WT, which may be valuable for synthetic biology applications of dYmu1-WFR. A state-specific dissociation constant can be defined for each of the *R*-loop states, quantifying the binding strength of the RNP to the DNA: $K_{D,I} = \frac{k_{IC}}{k'_{CI}}$ and $K_{D,O} = \frac{k_{IC}}{k'_{CI}} * \frac{k_{OI}}{k_{IO}}$ are the dissociation constants for the I and O states, respectively, where the measured $k_{CI} = k'_{CI} * [RNP]$. An overall dissociation constant is then given by:

$$K_D = \frac{1}{\frac{k'_{CI}}{k_{IC}} + \frac{k'_{CI}}{k_{IC}} * \frac{k_{IO}}{k_{OI}}} = \frac{1}{(K_{D,I})^{-1} + (K_{D,O})^{-1}}$$

Using the measured single-molecule kinetic parameters (Table S3), for WT TnpB, $K_D \sim K_{D,I} \sim 400\ nM$, while for WFR TnpB $K_D \sim K_{D,O} \sim 0.4\ nM$, indicating ~1000-fold tighter binding for WFR. This analysis assumes that binding of RNP to DNA in the absence of *R*-loop formation is weak and can be neglected, as is known for SpyCas9[S6].

**Note S2. Phylogenetic and structural analysis of Ymu1 TnpB mutations**

To understand how mutations in Ymu1-WFR reshape the DNA unwinding landscape, we combined sequence co-evolution analysis[S7] of ~4,500 non-redundant TnpB homologs, AlphaFold3 (AF3) modeling of the Ymu1-reRNA-DNA ternary complex, and comparison with available cryo-EM structures of related TnpB homologs[S3-5] (see Methods). Together, these analyses suggest two classes of effects: stabilization of early RNA-DNA engagement and modulation of protein conformational transitions that favor later, more extensively unwound states.

<u>Proposed structural correspondence of C, I, and O states</u>

We propose a structural framework relating cryo-EM conformations of a related ortholog (Dra2) to the unwinding intermediates resolved by AuRBT. The Dra2 protein-RNA binary complex[S3] is most relevant to the closed (C) state (Fig. S6A-B). The Dra2 ternary structure reported by Nakagawa et al.[S4] exhibits cryo-EM density consistent with limited ordering of the TAM-distal RNA-DNA heteroduplex, and the strongly resolved proximal heteroduplex principally engages the REC lobe (Fig. S6A, S6C). We propose that the partially unwound intermediate (I) state observed in our experiments, in which only a short RNA-DNA heteroduplex is formed, may adopt a protein-RNA-DNA architecture resembling this structure. By contrast, the ternary structure reported by Sasnauskas et al.[S3], which resolves ~15 bp of ordered heteroduplex, may correspond to the fully unwound open (O) state (Fig. S6A, S6D). In this putative O-state architecture, the extended RNA-DNA heteroduplex engages both the REC and NUC lobes, including the bridge helix and lid domain (Fig. S6D). Across these structures, the lid domain appears disordered in the C state and becomes progressively ordered. We propose the lid may have transient interactions with the partially formed RNA-DNA heteroduplex in the I state, and then becomes well-ordered and engages the heteroduplex in the fully unwound O state, suggesting coupling between lid ordering and late-stage *R*-loop formation as observed in related Cas12 enzymes[S8-10]. AF3 modeling of the Ymu1 ternary complex converges on a similar fully unwound O-state architecture (Methods), providing a structural reference for interpreting mutational effects discussed shortly.

<u>Proposed role of H4W in initiation of unwinding</u>

While sequence co-evolution analysis revealed little evidence of coupling between H4 and other amino acids in TnpB (Fig. S6E), AF3 modeling places H4 adjacent to the first RNA-DNA base pair of the heteroduplex (Fig. S6F). Substitution of His with Trp may introduce a larger aromatic surface to enhance π-stacking at the heteroduplex terminus, stabilizing initial heteroduplex formation. Together with our AuRBT data showing preferential stabilization of the I state by H4W, and prior reports of activity-enhancing substitutions at homologous positions in TnpB and Cas12f enzymes[S11,12], these observations support a model in which H4W enhances unwinding by stabilizing early RNA-DNA engagement.

<u>Proposed role of L304F and V305R in post-initiation unwinding</u>

Co-conservation analysis identifies several evolutionarily coupled residues surrounding V305 (I278, D280, Q309, and R321) (Fig. S6G), consistent with the AF3 model in which V305R is immediately adjacent to this network (Fig. S6H). More specifically, the AF3-predicted arginine 305 residue in the lid-adjacent helix is oriented to form an interaction with D280 on the counterpart RuvC β-strand, with I278, Q309, and R321 forming additional contacts, suggesting a molecular mechanism for lid domain stabilization in the ternary conformation. Co-evolution analysis of L304 likewise identifies a hydrophobic cluster (L262, S266, L308), consistent with AF3 modeling, which places L304 in a tightly packed hydrophobic pocket formed by these residues from the lid-adjacent helix and the bridge helix (Fig. S6I-J). Substitution with a bulkier aromatic side chain (L304F) could provide improved packing within this pocket. Consistent with this interpretation, deep mutational scanning of Dra2 TnpB indicates that the native Phe is preferred over Leu at the position homologous to Ymu1 L304[S12]. Together, these structural and covariation analyses support a model in which L304F and V305R strengthen interactions within the lid-proximal region of the RuvC domain, thereby stabilizing the I and O unwound states and complementing the effect of H4W, consistent with our AuRBT experiment (Fig. 4). The greater stabilization conferred by WFR in the O state compared with the I state (energy minima in Fig. 4J) could be explained by increased lid ordering and larger contributions to stability for lid-adjacent residues in the O state. In the I state the shorter heteroduplex adopts a distinct REC-bound confirmation that may support only transient interactions with the lid domain, whereas the lid is ordered and engages the fully formed heteroduplex in the O state.

Recent cryo-EM structures of an artificial TnpB designed using evolutionary information and ESM inverse folding (ESM-IF), in which F303 and R304 occupy positions homologous to Ymu1 L304 and V305 in a similar residues context, provide additional structural explanation to the proposed stabilization mechanisms[S5]. Across reported conformations, both the lid domain and the Ymu1's V305-homologous residue R304 are unresolved in the initial TAM-bound structure (which may correspond to a state intermediate between C and I) but become ordered in the fully unwound state (O) (Fig. S6K). Based on this experimental observation, we suggest that Ymu1's V305R similarly supports formation of the RuvC and lid domain conformations in the late unwound states (Fig. S6K). In contrast, F303 is structured in both early and late states (Fig. S6L), and may contribute to overall RuvC fold stabilization across the unwinding pathway.

### Supplemental Tables

#### Table S1. Target site sequences and amplicon-sequencing primers used in plant genome-editing experiments (related to Fig. 1 and Methods)

| Gene ID | Target site | Target site sequence (5′-3′) | Forward primer sequence (5′-3′) | Reverse primer sequence (5′-3′) |
|---|---|---|---|---|
| AT4G14210 | AtPDS3_g2 | aaggcaaattcgccgc | ACACTCTTTCCCTACACGACGCTCTTCCGATCTgaagcagttgtgagttaagttggaga | GTGACTGGAGTTCAGACGTGTGCTCTTCCGATCTttgtcttaagcgcttgagaagtgg |
| AT4G18480 | AtCHLI1_g4 | CTGTTACCTGAGATTA | ACACTCTTTCCCTACACGACGCTCTTCCGATCTGTGGTGTTATGATTATGGGAGATAGAG | GTGACTGGAGTTCAGACGTGTGCTCTTCCGATCTTCGCAATAACAGGAACTTGCTC |
| AT4G18480 | AtCHLI1_g6 | GAAGTTAATCTCTTGG | ACACTCTTTCCCTACACGACGCTCTTCCGATCTAAGCCTTTGAGCCTGGTTTG | GTGACTGGAGTTCAGACGTGTGCTCTTCCGATCTCGGGTGAGAAATCGAAATCCC |
| AT4G18480 | AtCHLI1_g9 | CGGTTTGGTATGCATG | ACACTCTTTCCCTACACGACGCTCTTCCGATCTGCGAGGTTTATCTTGATCGGTTC | GTGACTGGAGTTCAGACGTGTGCTCTTCCGATCTTCACGGAAATCCTTTGGGTTACTA |
| AT5G45930 | AtCHLI2_g1 | CTCTGTAGCGACATTC | ACACTCTTTCCCTACACGACGCTCTTCCGATCTtgcagAGAAACTTTCTGGAAGAATCCA | GTGACTGGAGTTCAGACGTGTGCTCTTCCGATCTtcattccaaaggttcaagcttttaatc |
| AT5G45930 | AtCHLI2_g3 | TTACCAGTTCCTCTAT | ACACTCTTTCCCTACACGACGCTCTTCCGATCTGGACAAGATGAGATGAAGCTATGCCTT | GTGACTGGAGTTCAGACGTGTGCTCTTCCGATCTCTCGGGTCTGAGTTATACGGATCA |
| AT5G45930 | AtCHLI2_g8 | GAAGTTAATCTTTTGG | ACACTCTTTCCCTACACGACGCTCTTCCGATCTAGGCGTTTGAGCCTGGACTACTA | GTGACTGGAGTTCAGACGTGTGCTCTTCCGATCTAGCAGGATGAGAAATCGATATCCCTTC |
| AT5G45930 | AtCHLI2_g10 | TCAGCTGCATCTGGTT | ACACTCTTTCCCTACACGACGCTCTTCCGATCTTAGCTAAAGCTAATAGAGGGATTCTTTATGTTGAT | GTGACTGGAGTTCAGACGTGTGCTCTTCCGATCTGTGGTCTAAGCTCTCCTTCTTCAGGA |
| AT2G46410 | AtCPC_g1 | cggtttgagtctgatt | ACACTCTTTCCCTACACGACGCTCTTCCGATCTCCAAGGCTTCTTGTTCCGAAG | GTGACTGGAGTTCAGACGTGTGCTCTTCCGATCTCCAACGAGTTTATACATCCGAGAA |

| | | | | |
|---|---|---|---|---|
| AT2G46410 | AtCPC_g5 | tacagcatgtttgtat | ACACTCTTTCCCTACACGACGCTCTTCCGATCTtgattctttagcaaaacatattcttaatttatgtca | GTGACTGGAGTTCAGACGTGTGCTCTTCCGATCTccgataaaaaccgcataaagtttgt |
| AT4G14210 | AtPDS3_g5 | tacccatcctaaagta | ACACTCTTTCCCTACACGACGCTCTTCCGATCTgacgtcaggaagaacatggtcatttg | GTGACTGGAGTTCAGACGTGTGCTCTTCCGATCTagaacatttcagcgctaatgctacaa |
| AT4G14210 | AtPDS3_g10 | taacttgtactacctc | ACACTCTTTCCCTACACGACGCTCTTCCGATCTtgttgcagcttatgacacacaataag | GTGACTGGAGTTCAGACGTGTGCTCTTCCGATCTgaatctttatctcactggcaatcata |
| AT4G14210 | AtPDS3_g12 | gcgttggagcatataa | ACACTCTTTCCCTACACGACGCTCTTCCGATCTgaaccgacccgagaagagatttg | GTGACTGGAGTTCAGACGTGTGCTCTTCCGATCTtggaatacacacatttgtacaacca |
| AT5G53200 | AtTRY_g3 | AGCAGGAAGAGTTCCT | ACACTCTTTCCCTACACGACGCTCTTCCGATCTtttattatgaaaatataaatgctaatgcttgggat | GTGACTGGAGTTCAGACGTGTGCTCTTCCGATCTTGGCGTCGTTTATCAGCAAAG |

**Table S2. Rate constants ($k_{obs}$) for DNA cleavage assay (n=3) (related to Fig. 3, Fig. 5 and Fig. S3, Fig. S5)**

| Protein | Target | Strand | Mean ($min^{-1}$) | SD ($min^{-1}$) |
|---|---|---|---|---|
| **WT Ymu1** | 1 | NTS | 0.0676 | 0.0033 |
| | | NTS (with salmon sperm) | 0.0212 | 0.0017 |
| | | NTS (substrate excess) | 0.00994 | 0.0040 |
| | | TS | 0.0717 | 0.0090 |
| | | TS (with salmon sperm) | 0.0238 | 0.0018 |
| | | TS (substrate excess) | 0.00541 | 0.0025 |
| | 2 | NTS | 0.0152 | 0.0019 |
| | | TS | 0.0179 | 0.0019 |
| **H4W** | 1 | NTS | 0.168 | 0.039 |

| | | | | |
|---|---|---|---|---|
| | | TS | 0.176 | 0.055 |
| **V305R** | 1 | NTS | 0.198 | 0.025 |
| | | TS | 0.179 | 0.023 |
| **Ymu1-WFR** | 1 | NTS | 1.90 | 0.74 |
| | | NTS (with salmon sperm) | 1.23 | 0.18 |
| | | NTS (substrate excess) | 0.555 | 0.20 |
| | | TS | 0.980 | 0.33 |
| | | TS (with salmon sperm) | 0.705 | 0.058 |
| | | TS (substrate excess) | 0.323 | 0.11 |
| | 2 | NTS | 0.0793 | 0.0038 |
| | | TS | 0.0380 | 0.0032 |
| **L304F** | 1 | NTS | 0.0692 | 0.0043 |
| | | TS | 0.0737 | 0.0023 |

**Table S3. Transition rate constants from equilibrium AuRBT assays with fully matched target sequences (related to Fig. 4 and Fig. S4)**

| **RNP** | **Conc** | $k_{C\to I}$ **($s^{-1}$)** | $k_{C\to I}$ **error ($s^{-1}$)** | $k_{I\to C}$ **($s^{-1}$)** | $k_{I\to C}$ **error ($s^{-1}$)** | $k_{I\to O}$ **($s^{-1}$)** | $k_{I\to O}$ **error ($s^{-1}$)** | $k_{O\to I}$ **($s^{-1}$)** | $k_{O\to I}$ **error ($s^{-1}$)** |
|---|---|---|---|---|---|---|---|---|---|
| WT dYmu1 | 100 nM | 0.084199 | 0.0024522 | 0.31662 | 0.0092328 | 0.010231 | 0.0016597 | 0.36536 | 0.05927 |
| H4W dYmu1 | 100 nM | 0.063184 | 0.0029022 | 0.073818 | 0.0033763 | 0.0043241 | 0.00081717 | 0.29539 | 0.056847 |
| dYmu1-WFR | 100 nM | 0.62308 | 0.071006 | 0.01729 | 0.0019577 | 0.012856 | 0.0016881 | 0.002112 | 0.00028222 |

**Table S4. AuRBT trace statistics for equilibrium AuRBT assays (related to Fig. 4, Fig. S4, and Methods)**

| Condition | Tethers | Unique Chambers | Time Tracked (s) | C → I | I → C | I → O | O → I | C → O | O → C |
|---|---|---|---|---|---|---|---|---|---|
| WT dYmu1 (100nM) | 5 | 3 | 17821 | 1179 | 1176 | 38 | 38 | 2 | 3 |
| H4W dYmu1 (100nM) | 5 | 5 | 14069 | 474 | 478 | 28 | 27 | 1 | 2 |
| dYmu1-WFR (100nM) | 5 | 4 | 31151 | 77 | 78 | 58 | 56 | 2 | 1 |
| WT dYmu1 (100nM) 1-4 nt mismatch | 2 | 1 | 3500 | 0 | 0 | 0 | 0 | 0 | 0 |
| dYmu1-WFR (100 nM) 1-4 nt mismatch | 2 | 1 | 4693 | 0 | 0 | 0 | 0 | 0 | 0 |

Column “i → j” gives number of transitions observed from state i to state j.

**Table S5. Transition rate constants of WT dYmu1 for non-equilibrium, torsion-driven AuRBT assays with fully matched target sequences (related to Fig. 5 and Fig. S5)**

| Twist | $k_{C\to I}$ [$s^{-1}$] | $k_{C\to I}$ error [$s^{-1}$] | $k_{I\to C}$ [$s^{-1}$] | $k_{I\to C}$ error [$s^{-1}$] | $k_{I\to O}$ [$s^{-1}$] | $k_{I\to O}$ error [$s^{-1}$] | $k_{O\to I}$ [$s^{-1}$] | $k_{O\to I}$ error [$s^{-1}$] |
|---|---|---|---|---|---|---|---|---|
| -4.75 | 0.1371 | 0.0457 | 0.0066 | 0.0023 | 0.0270 | 0.0047 | 0.0167 | 0.0034 |
| -4 | 0.3410 | 0.0633 | 0.0159 | 0.0035 | 0.0310 | 0.0048 | 0.0230 | 0.0042 |
| -3 | 0.2784 | 0.0366 | 0.0363 | 0.0052 | 0.0185 | 0.0037 | 0.0184 | 0.0042 |
| -2 | 0.2387 | 0.0231 | 0.0699 | 0.0071 | 0.0087 | 0.0025 | 0.0415 | 0.0088 |
| -1 | 0.1139 | 0.0112 | 0.1624 | 0.0152 | 0.0043 | 0.0025 | 0.0367 | 0.0116 |
| 0 | 0.0657 | 0.0073 | 0.4068 | 0.0413 | NA | NA | 0.1347 | 0.0449 |
| 1 | 0.0331 | 0.0049 | 0.5525 | 0.0815 | NA | NA | NA | NA |
| 2 | 0.0137 | 0.0031 | 0.6544 | 0.1636 | NA | NA | NA | NA |
| 3 | 0.0021 | 0.0012 | 0.9507 | 0.4753 | NA | NA | NA | NA |
| 4 | NA | NA | NA | NA | NA | NA | NA | NA |

**Table S6. Parameters from linear fits of ln($K_{ij}$) and ln($k_{ij}$) vs imposed twist (related to Fig. 5 and Fig. S5)**

| Transition | $\Delta\theta_{ij}$ (bp) | $\Delta G_{ij}(0)$ ($k_BT$) | $\Delta G_{ij}$(-5) ($k_BT$) | $\Delta\theta^{\ddagger}_{ij}$ (bp) | $ln(k_{ij}(0))$ | $\Delta\theta^{\ddagger}_{ji}$ (bp) | $ln(k_{ji}(0))$ |
|---|---|---|---|---|---|---|---|

| | | | | | | | |
|---|---|---|---|---|---|---|---|
| C↔I | 5.3 | 1.6 | -4.0 | 3.3 | -2.8 | -4.0 | -0.9 |
| I↔O | 3.6 | 2.8 | -1.0 | 3.2 | -6.1 | -1.8 | -2.5 |

$\Delta\theta_{ij}$, reported in number of base pairs unwound, gives the predicted difference in equilibrium twist between states i and j based on the slope of the $ln(K_{ij})$ vs twist plot. $\Delta G_{ij}(0)$ and $\Delta G_{ij}$(-5) give the predicted free energy difference between the states at 0 and -5 twist, respectively. Linear fits calculated from equally-weighted $ln(K_{ij})$ data points.

$\Delta\theta^{\ddagger}{}_{ij}$, reported in the number of base pairs unwound, gives the predicted location of the transition state for the i→j transition in relation to the predicted equilibrium twist for state i. $ln(k_{ij}(0))$ gives the natural log of the predicted transition rate (in $s^{-1}$) from state i to state j at 0 twist. Linear fits calculated from equally-weighted data points selected from a linear portion of $ln(k_{ij})$ data.

**Table S7. AuRBT trace statistics for non-equilibrium torque-driven assays (related to Fig. 5, Fig. S5, and Methods)**

| Condition | Tethers | Unique Chambers | Cycles |
|---|---|---|---|
| WT dYmu1 (10nM) | 13 | 13 | 68 |

| Twist interval | C → I | I → C | I → O | O → I |
|---|---|---|---|---|
| [-5, -4.5] | 9 | 8 | 33 | 24 |
| [-4.5, -3.5] | 29 | 21 | 41 | 30 |
| [-3.5, -2.5] | 58 | 49 | 25 | 19 |
| [-2.5, -1.5] | 107 | 96 | 12 | 22 |
| [-1.5, -0.5] | 104 | 114 | 3 | 10 |
| [-0.5, 0.5] | 82 | 97 | 1 | 9 |
| [0.5, 1.5] | 45 | 46 | 0 | 0 |
| [1.5, 2.5] | 19 | 16 | 0 | 0 |
| [2.5, 3.5] | 3 | 4 | 0 | 0 |
| [3.5, 4.5] | 2 | 2 | 0 | 0 |
| [4.5, 5.5] | 0 | 0 | 0 | 0 |
| [5.5, 6.5] | 0 | 0 | 0 | 0 |
| [6.5, 7.5] | 0 | 0 | 0 | 0 |

Column “i → j” gives number of transitions observed from state i to state j.

**Table S8. Rate constants ($k_{obs}$) for plasmid cleavage assay (n=3) (related to Fig. 5)**

| Substrate Type | Protein | Mean (min$^{-1}$) | SD (min$^{-1}$) |
|---|---|---|---|
| **Linearized** | WT Ymu1 | 0.0288 | 0.0067 |
| | Ymu1-WFR | 0.683 | 0.14 |
| **Supercoiled** | WT Ymu1 | 3.41 | 1.3 |
| | Ymu1-WFR | 5.40 | 0.93 |

**Table S9. Rate constants ($k_{obs}$) for pre-unwound DNA cleavage assay (n=3) (related to Fig. 5 and Fig. S5)**

| Strand | DNA target | WT Ymu1 mean (min$^{-1}$) | WT Ymu1 SD (min$^{-1}$) | H4W mean (min$^{-1}$) | H4W SD (min$^{-1}$) | Ymu1-WFR mean (min$^{-1}$) | Ymu1-WFR SD (min$^{-1}$) |
|---|---|---|---|---|---|---|---|
| NTS | 1-2 nt pre-unwound substrate | 0.162 | 0.040 | 0.217 | 0.060 | 1.42 | 0.16 |
| | 8-9 nt pre-unwound substrate | 1.77 | 0.13 | 1.57 | 0.12 | 2.17 | 0.18 |
| | 1-9 nt pre-unwound substrate | 1.51 | 0.20 | 2.17 | 0.65 | 2.44 | 0.33 |
| TS | 1-2 nt pre-unwound substrate | 0.180 | 0.050 | 0.301 | 0.0043 | 1.16 | 0.13 |
| | 8-9 nt pre-unwound substrate | 1.29 | 0.22 | 1.14 | 0.18 | 1.49 | 0.24 |
| | 1-9 nt pre-unwound substrate | 1.02 | 0.16 | 1.38 | 0.22 | 1.21 | 0.17 |

The $k_{obs}$ value for the canonical substrate is shown in Table S2.

**Table S10. Cleavage levels and rate constants ($k_{obs}$) for DNA-RNA mismatch cleavage assay (n=3) (related to Fig. 3, Fig. S3, and Fig. S5)**

| Strand | DNA target | WT Ymu1 mean endpoint cleavage level (%) | WT Ymu1 mean endpoint SD (%) | WT Ymu1 $k_{obs}$ mean (min$^{-1}$) | WT Ymu1 $k_{obs}$ SD (min$^{-1}$) | WFR mean endpoint cleavage level (%) | WFR mean endpoint SD (%) | WFR $k_{obs}$ mean (min$^{-1}$) | WFR $k_{obs}$ SD (min$^{-1}$) |
|---|---|---|---|---|---|---|---|---|---|
| NTS | On-target | 92.9 | 0.81 | 0.0676 | 0.0033 | 97.0 | 0.47 | 2.00 | 0.74 |
| | 4 nt mismatch | 5.90 | 0.36 | (0.0671)† | (0.085)† | 31.8 | 4.7 | 0.00710 | 0.00094 |
| | 7 nt mismatch | 5.57 | 3.6 | (1.26)† | (1.2)† | 52.6 | 3.5 | 0.0111 | 0.0027 |
| | 10 nt mismatch | 7.57 | 1.5 | (0.0119)† | (0.0071)† | 62.1 | 2.3 | 0.0106 | 0.00075 |
| | 14 nt mismatch | 46.6 | 17 | 0.00696 | 0.0032 | 91.4 | 1.1 | 0.621 | 0.221 |
| TS | On-target | 90.9 | 3.0 | 0.0717 | 0.0090 | 97.8 | 0.26 | 0.980 | 0.33 |
| | 4 nt mismatch | 5.60 | 4.5 | (2.129)† | (0.68)† | 40.3 | 8.2 | 0.0192 | 0.0099 |
| | 7 nt mismatch | 8.27 | 2.0 | (0.503)† | (0.85)† | 56.0 | 3.2 | 0.0158 | 0.0019 |
| | 10 nt mismatch | 6.43 | 0.40 | (0.285)† | (0.47)† | 63.6 | 5.2 | 0.0182 | 0.0031 |
| | 14 nt mismatch | 49.9 | 16 | 0.00739 | 0.0067 | 96.7 | 0.85 | 0.335 | 0.059 |

†Fitted values are shown for completeness but are not interpreted quantitatively due to low cleavage level, which renders $k_{obs}$ unreliable for kinetic comparison.

**Table S11. Cas-OFFinder predicted off-target sites for *CHLI1* g4 (related to Fig. S5)**

| Genomic DNA (5′-3′) | Chromosome | Position | Strand | Mismatches | Classification | Variants Detected |
|---|---|---|---|---|---|---|
| TTGATtTGTTACCTGAGATag | chr1 | 385713 | + | 3 | Off-target | No SNP/InDel |
| aTGATCTGTTACCTGAtcTTt | chr1 | 456752 | - | 4 | Off-target | No SNP/InDel |
| aTGATgaGTTACCTGAGATTg | chr1 | 1091615 | + | 4 | Off-target | No SNP/InDel |
| TTGATCTcTTACCTccGATTt | chr1 | 5232246 | + | 4 | Off-target | No SNP/InDel |

| | | | | | | |
|---|---|---|---|---|---|---|
| TTGtTCTGcTtCCaGAGATTA | chr1 | 5486474 | - | 4 | Off-target | No SNP/InDel |
| TTGATtTGTTACCTaAtATTg | chr1 | 6483939 | - | 4 | Off-target | No SNP/InDel |
| TTtATCTaTTtCCTGAGAaTA | chr1 | 8044117 | - | 4 | Off-target | No SNP/InDel |
| TTcATCTGTTcCtTGtGATTA | chr1 | 8655476 | + | 4 | Off-target | No SNP/InDel |
| TTGtTCTGTTACCTGAtcTTt | chr1 | 8701665 | - | 4 | Off-target | No SNP/InDel |
| TTGAaCTGTTtCCTGtGAaTA | chr1 | 9221199 | - | 4 | Off-target | No SNP/InDel |
| TTGcTgTGTTACCTGAGAaTt | chr1 | 10672791 | - | 4 | Off-target | No SNP/InDel |
| TTcATCTcgTACCTGAtATTA | chr1 | 11053135 | + | 4 | Off-target | No SNP/InDel |
| TTGAaaaGTTAgCTGAGATTA | chr1 | 12716075 | + | 4 | Off-target | No SNP/InDel |
| TactTgTGTTACCTGAGATTA | chr1 | 13482217 | - | 4 | Off-target | No SNP/InDel |
| aTcATCTGTTtCCTGAGAaTA | chr1 | 13600705 | + | 4 | Off-target | No SNP/InDel |
| TgGcTCTGTTACaTGAaATTA | chr1 | 13691906 | + | 4 | Off-target | No SNP/InDel |
| gTGATCTtTTcCCTGAaATTA | chr1 | 14832632 | + | 4 | Off-target | No SNP/InDel |
| gTGATCTtTTcCCTGAaATTA | chr1 | 14913382 | - | 4 | Off-target | No SNP/InDel |
| gTGATgTGTTcCCTGAaATTA | chr1 | 16029023 | + | 4 | Off-target | No SNP/InDel |
| TTGATaTaTTtCCTGAGATgA | chr1 | 17648564 | + | 4 | Off-target | No SNP/InDel |
| TTGAgCTaTTAgaTGAGATTA | chr1 | 18499222 | - | 4 | Off-target | No SNP/InDel |
| TTGATCTGTTtCaGAGATTt | chr1 | 19948765 | + | 4 | Off-target | No SNP/InDel |
| TTGATCTGTTACCTaActTaA | chr1 | 21519072 | + | 4 | Off-target | No SNP/InDel |
| TTGATgTGcTtCCTGAcATTA | chr1 | 22325011 | - | 4 | Off-target | No SNP/InDel |
| TcGATCTGTgACCTcAGATTt | chr1 | 24250207 | + | 4 | Off-target | No SNP/InDel |
| TTGATCTcTTAgCTcAGATTc | chr1 | 26390110 | - | 4 | Off-target | No SNP/InDel |
| TTGtTCTGTTtCCTcAaATTA | chr1 | 29715101 | - | 4 | Off-target | No SNP/InDel |
| TTGATCTGaTgCCTGAGAcTc | chr1 | 29907423 | + | 4 | Off-target | No SNP/InDel |
| TTtATCTaTTtCCTGAGAcTA | chr2 | 4774056 | - | 4 | Off-target | No SNP/InDel |

| | | | | | | |
|---|---|---|---|---|---|---|
| TTGAgCTGTTtCCTGtGATaA | chr2 | 6423541 | + | 4 | Off-target | No SNP/InDel |
| TTGAagTGTTACgTGAGAcTA | chr2 | 6984586 | - | 4 | Off-target | No SNP/InDel |
| TTGATaTGaTACaaGAGATTA | chr2 | 8181489 | - | 4 | Off-target | No SNP/InDel |
| TTGAaCTGTTcCCTtAGATaA | chr2 | 8826433 | - | 4 | Off-target | No SNP/InDel |
| TTGATCTGTTAttaaAGATTA | chr2 | 9993416 | + | 4 | Off-target | No SNP/InDel |
| TTGATgTGTgACCTGAtATTA | chr2 | 11593142 | - | 3 | Off-target | No SNP/InDel |
| TTGATCTaTTACCTaAGATcA | chr2 | 12290104 | - | 3 | Off-target | No SNP/InDel |
| TTGATCTcTTAtaTGAGgTTA | chr2 | 12900822 | + | 4 | Off-target | No SNP/InDel |
| TgGATCTGaaAtCTGAGATTA | chr2 | 13747304 | - | 4 | Off-target | No SNP/InDel |
| TTtcTCTGTTAtCTGAGAaTA | chr2 | 14024696 | - | 4 | Off-target | No SNP/InDel |
| TTGATaTcTTACtTGAGATTt | chr2 | 15133368 | + | 4 | Off-target | No SNP/InDel |
| aTGAaCTGTTACCTGAGATTc | chr2 | 15659366 | + | 3 | Off-target | No SNP/InDel |
| TTGATCaGTTACaTttGATTA | chr2 | 18248786 | - | 4 | Off-target | No SNP/InDel |
| TTGtTaTGTTAgCTGAGATTg | chr2 | 18317151 | + | 4 | Off-target | No SNP/InDel |
| TTGAaCTGTTtCaTGAGcTTA | chr2 | 18905832 | - | 4 | Off-target | No SNP/InDel |
| TTaATCTaTaACCTtAGATTA | chr3 | 218050 | - | 4 | Off-target | No SNP/InDel |
| TTGATCgGTTAggTGAGATTg | chr3 | 2589316 | - | 4 | Off-target | No SNP/InDel |
| TTGATaTGTTtCaTGAGATgA | chr3 | 4058168 | - | 4 | Off-target | No SNP/InDel |
| TTGATtTGTTgtCTGAGATTt | chr3 | 6726855 | + | 4 | Off-target | No SNP/InDel |
| gTcATCTGTTtCCTGAGAaTA | chr3 | 11854134 | + | 4 | Off-target | No SNP/InDel |
| TTGATCTGTTgatTGAGATTt | chr3 | 17319931 | + | 4 | Off-target | No SNP/InDel |
| TTttTCTGTTAgCTGAGATgA | chr3 | 18350447 | - | 4 | Off-target | No SNP/InDel |
| TTGAgCTGTTgtCTGAGATTc | chr3 | 20352482 | + | 4 | Off-target | No SNP/InDel |
| TTGATCTcTTtCCTtAGATTt | chr3 | 20820280 | - | 4 | Off-target | No SNP/InDel |
| TTtATgTGTTACtTGAaATTA | chr4 | 4089641 | - | 4 | Off-target | No SNP/InDel |

| | | | | | | |
|---|---|---|---|---|---|---|
| TTGATtTGgTACCTGAGAcTg | chr4 | 5859717 | + | 4 | Off-target | No SNP/InDel |
| TTGATtTGgTACCTGAGAcTg | chr4 | 5877024 | + | 4 | Off-target | No SNP/InDel |
| TTGATtTGgTACCTGAGAcTg | chr4 | 5903286 | + | 4 | Off-target | No SNP/InDel |
| TTtATCTGTTACCTGAaAaaA | chr4 | 8342444 | - | 4 | Off-target | No SNP/InDel |
| TTaATCTGTTAaCTaAaATTA | chr4 | 9193485 | + | 4 | Off-target | No SNP/InDel |
| TTGATCTGTTACCTGAGATTA | chr4 | 10202753 | - | 0 | On-target | Edited (InDel) |
| cgGATCaGTTACCTGAGcTTA | chr4 | 11126679 | + | 4 | Off-target | No SNP/InDel |
| TTGgTtTGTTACCTGAGAagA | chr4 | 12917278 | - | 4 | Off-target | No SNP/InDel |
| TTGATgTGTTAgCTGAaATgA | chr4 | 16603188 | - | 4 | Off-target | No SNP/InDel |
| TTGATCTtTTACCaGAcATTc | chr5 | 1158214 | - | 4 | Off-target | No SNP/InDel |
| TTGATCTtTTACCaGAGgcTA | chr5 | 3746803 | - | 4 | Off-target | No SNP/InDel |
| cTGgTCTGTcACCTGAGATaA | chr5 | 5295761 | + | 4 | Off-target | No SNP/InDel |
| TTGATCTGaTAtCTcAGATTc | chr5 | 6998326 | + | 4 | Off-target | No SNP/InDel |
| aTGATCTGTcACCTGtaATTA | chr5 | 9245291 | - | 4 | Off-target | No SNP/InDel |
| TgGtTCTGTTtCCTGAGATTt | chr5 | 9636891 | + | 4 | Off-target | No SNP/InDel |
| TTtAcCTGTTcCCTGAGtTTA | chr5 | 9794815 | + | 4 | Off-target | No SNP/InDel |
| aTcATCTGTTAgCTtAGATTA | chr5 | 10629268 | - | 4 | Off-target | No SNP/InDel |
| TaGAaCTGTTACCTcAGATTt | chr5 | 11632265 | + | 4 | Off-target | No SNP/InDel |
| TTcATCTaTTtCCTGAGAaTA | chr5 | 12436965 | - | 4 | Off-target | No SNP/InDel |
| TTGATaTGTTtCCTcAGAaTA | chr5 | 13992951 | - | 4 | Off-target | No SNP/InDel |
| TTGATCTGTaAttTGAaATTA | chr5 | 16015303 | + | 4 | Off-target | No SNP/InDel |
| TTGATtTGcTtCCTGAGATcA | chr5 | 18628687 | + | 4 | Off-target | No SNP/InDel |
| aTGtTCTGgTACCTGAGATTg | chr5 | 19728238 | + | 4 | Off-target | No SNP/InDel |
| TaGATgTGTTACCTGAaATcA | chr5 | 19844847 | - | 4 | Off-target | No SNP/InDel |
| TTGATCTtggACCaGAGATTA | chr5 | 20514416 | - | 4 | Off-target | No SNP/InDel |

| | | | | | | |
|---|---|---|---|---|---|---|
| TTGcTCTGaTACCTGAGAggA | chr5 | 20969420 | - | 4 | Off-target | No SNP/InDel |
| TTGATCTGTTACtTGAagaTA | chr5 | 23435591 | - | 4 | Off-target | No SNP/InDel |
| TTGATCaGTTACtTGtGATgA | chr5 | 24852608 | - | 4 | Off-target | No SNP/InDel |
| TTGATaTGTTcCaaGAGATTA | chr5 | 26888514 | + | 4 | Off-target | No SNP/InDel |

Potential off-target and on-target sites predicted by Cas-OFFinder (v3.0.0) for the *CHLI1* g4 site, allowing up to 4 mismatches across the TAM and guide sequence (5′-TTGATCTGTTACCTGAGATTA-3′). **Genomic DNA**: matched genomic sequence with mismatches shown in lowercase. **Chromosome**: Arabidopsis thaliana TAIR10 chromosome. **Position**: genomic coordinate. **Strand**: (+) forward, (-) reverse. **Mismatches**: number of mismatches between gRNA and genomic DNA. **Classification**: on-target or off-target. **Variants Detected**: SNPs or InDels identified at this site across all three sequenced progeny lines after filtering against wild-type controls.

#### Table S12. Cas-OFFinder predicted off-target sites for *PDS3* g12 (related to Fig. S5)

| Genomic DNA (5′-3′) | Chromosome | Position | Strand | Mismatches | Classification | Variants Detected |
|---|---|---|---|---|---|---|
| TTGATGCGTTtGAGaATcTgA | chr1 | 603266 | + | 4 | Off-target | No SNP/InDel |
| TTGATaCtTTGGAGCtTATcA | chr1 | 1112526 | - | 4 | Off-target | No SNP/InDel |
| TTGATGatTTGGtcCATATAA | chr1 | 2663557 | + | 4 | Off-target | No SNP/InDel |
| TTGATGtGTTtGAGatTATAA | chr1 | 4238242 | - | 4 | Off-target | No SNP/InDel |
| TTGATGCGTTGGtGCATAatc | chr1 | 5305662 | + | 4 | Off-target | No SNP/InDel |
| TTGgTGtGgTGaAGCATATAA | chr1 | 5880417 | - | 4 | Off-target | No SNP/InDel |
| TTGtTtCGTaGGAGCATATAt | chr1 | 9814485 | - | 4 | Off-target | No SNP/InDel |
| TTGATGCaTTatAtCATATAA | chr1 | 10322285 | - | 4 | Off-target | No SNP/InDel |
| TTGATGatTTGGAtgATATAA | chr1 | 11292702 | - | 4 | Off-target | No SNP/InDel |
| TTGgTGCtTTGGtGCATATtA | chr1 | 13817767 | - | 4 | Off-target | No SNP/InDel |
| TgGATatGTTGGAGgATATAA | chr1 | 17843545 | + | 4 | Off-target | No SNP/InDel |
| TTGAgGCGTTGGAGaATATAA | chr1 | 18348024 | + | 2 | Off-target | No SNP/InDel |
| TTGATGCtTTtGcGCATAaAA | chr1 | 19411643 | + | 4 | Off-target | No SNP/InDel |
| TTGATGgGTTGGtGgATAgAA | chr1 | 19953639 | + | 4 | Off-target | No SNP/InDel |
| TTGATGatgTGGAcCATATAA | chr1 | 23704848 | - | 4 | Off-target | No SNP/InDel |

| | | | | | | |
|---|---|---|---|---|---|---|
| TTGATGCGTTGGAtagTcTAA | chr1 | 24288161 | + | 4 | Off-target | No SNP/InDel |
| TTGtTGCtTTGGAGCATcTgA | chr1 | 25571517 | - | 4 | Off-target | No SNP/InDel |
| TTGtTatGTTGGAtCATATAA | chr1 | 28559455 | - | 4 | Off-target | No SNP/InDel |
| TaaATGCGTTGcAGCATcTAA | chr1 | 29389067 | + | 4 | Off-target | No SNP/InDel |
| TTGgTGCtTTGGtGCATATtA | chr2 | 3221482 | + | 4 | Off-target | No SNP/InDel |
| TTGATGtGTTGtAcCATAaAA | chr2 | 8176569 | + | 4 | Off-target | No SNP/InDel |
| TTGATGCGTTGaAGCtTAaAA | chr3 | 9345906 | + | 3 | Off-target | No SNP/InDel |
| aTGATGCcTTtGAGCAgATAA | chr3 | 12719791 | - | 4 | Off-target | No SNP/InDel |
| TTGATGCGTTGGAGCATcTct | chr3 | 15181264 | + | 3 | Off-target | No SNP/InDel |
| TTGATGCGTTtGAGaATgaAA | chr3 | 15369383 | - | 4 | Off-target | No SNP/InDel |
| TTGAaGCGaTGGAGaAgATAA | chr3 | 18021663 | - | 4 | Off-target | No SNP/InDel |
| TTGATGCGTTtGtGCATATgt | chr3 | 21020817 | - | 4 | Off-target | No SNP/InDel |
| TTGAgGCGTTGGtGCAcATgA | chr3 | 21997323 | - | 4 | Off-target | No SNP/InDel |
| TTaAgGCGTTGGAGCAgATcA | chr4 | 833322 | - | 4 | Off-target | No SNP/InDel |
| TcGATcCGTTtGAGCATATAt | chr4 | 2315047 | + | 4 | Off-target | No SNP/InDel |
| TTGATGCtTTGGgGaATAcAA | chr4 | 4218388 | + | 4 | Off-target | No SNP/InDel |
| TTGATGCGTTGGAGCATATAA | chr4 | 8195559 | - | 0 | On-target | Edited (InDel) |
| TTGATGCGTTGaAGaAcAaAA | chr4 | 11019148 | - | 4 | Off-target | No SNP/InDel |
| TTaATaCGTTGcAGCATATAg | chr4 | 11228135 | - | 4 | Off-target | No SNP/InDel |
| cTGATGCGaTGGAGCATAcAg | chr4 | 13346826 | + | 4 | Off-target | No SNP/InDel |
| TTcATGCGTTGcActATATAA | chr4 | 14631703 | - | 4 | Off-target | No SNP/InDel |
| TTGAgGCtTTtGAGCATATAt | chr5 | 124947 | + | 4 | Off-target | No SNP/InDel |
| TTGATGCGTatGAatATATAA | chr5 | 4677310 | - | 4 | Off-target | No SNP/InDel |
| TcGAatCGTTGGAtCATATAA | chr5 | 5784463 | + | 4 | Off-target | No SNP/InDel |
| TTtATtCGTatGAGCATATAA | chr5 | 6069715 | - | 4 | Off-target | No SNP/InDel |

| | | | | | | |
|---|---|---|---|---|---|---|
| TTGATcCGTTGGAGgAaAaAA | chr5 | 6389952 | - | 4 | Off-target | No SNP/InDel |
| gTGATGCGTTGGAGtATcTtA | chr5 | 8922973 | - | 4 | Off-target | No SNP/InDel |
| TTGATGCGgTGGAatATATAt | chr5 | 15908103 | - | 4 | Off-target | No SNP/InDel |
| TTGATGCcTTGaAGCATcTAc | chr5 | 16590425 | - | 4 | Off-target | No SNP/InDel |
| aTacTGCGTTtGAGCATATAA | chr5 | 18614163 | + | 4 | Off-target | No SNP/InDel |
| TTGATGatTTGGAtgATATAA | chr5 | 18851829 | + | 4 | Off-target | No SNP/InDel |
| TTGATtgaTTGGAaCATATAA | chr5 | 22644838 | + | 4 | Off-target | No SNP/InDel |
| TTGgTGCGTgGaAGCATAaAA | chr5 | 23387269 | + | 4 | Off-target | No SNP/InDel |
| TTGAaGCcTTGGAGaAaATAA | chr5 | 26390084 | - | 4 | Off-target | No SNP/InDel |

As described for Table S11, for the PDS3 g12 site (5′-TTGATGCGTTGGAGCATATAA-3′).

**Table S13. Filtered variants identified in whole-genome sequencing of Ymu1-WFR edited progeny (related to Fig. S5)**

| Chromosome | Start | End | Line 126 | Line 153 | Line 176 | On-target |
|---|---|---|---|---|---|---|
| chr1 | 929479 | 929579 | het SNP | ND | ND | |
| chr1 | 3825964 | 3826064 | ND | homo SNP | homo SNP | |
| chr1 | 5170759 | 5170859 | homo SNP | homo SNP | homo SNP | |
| chr1 | 8794889 | 8794989 | het SNP | het SNP | het SNP | |
| chr1 | 9550381 | 9550481 | homo SNP | homo SNP | homo SNP | |
| chr1 | 10641247 | 10641347 | ND | ND | het SNP | |
| chr1 | 11146968 | 11147068 | homo SNP | homo SNP | homo SNP | |
| chr1 | 12874513 | 12874615 | ND | ND | het InDel | |
| chr1 | 14044732 | 14045180 | het SNP | het SNP | het SNP | |
| chr1 | 15084663 | 15084769 | het SNP | het SNP | het SNP | |
| chr1 | 15096854 | 15096954 | het SNP | het SNP | het SNP | |
| chr1 | 15100724 | 15100824 | het SNP | het SNP | het SNP | |

| | | | | | |
|---|---|---|---|---|---|
| chr1 | 15208435 | 15208536 | het SNP | het SNP | het SNP |
| chr1 | 18122011 | 18122111 | het SNP | het SNP | het SNP |
| chr1 | 18437377 | 18437477 | ND | ND | het SNP |
| chr1 | 24005339 | 24005452 | het InDel | ND | ND |
| chr1 | 24972594 | 24972694 | het SNP | ND | ND |
| chr2 | 2069663 | 2069763 | het SNP | ND | ND |
| chr2 | 4196468 | 4196568 | homo SNP | homo SNP | homo SNP |
| chr2 | 5067286 | 5067386 | homo SNP | homo SNP | homo SNP |
| chr2 | 6261428 | 6261528 | het SNP | homo SNP | homo SNP |
| chr2 | 11117881 | 11117981 | homo SNP | homo SNP | homo SNP |
| chr2 | 14876309 | 14876409 | ND | het SNP | homo SNP |
| chr3 | 5279155 | 5279255 | ND | het SNP | ND |
| chr3 | 5818127 | 5818227 | ND | ND | homo SNP |
| chr3 | 11547524 | 11547624 | homo SNP | homo SNP | homo SNP |
| chr3 | 12152993 | 12153093 | homo SNP | homo SNP | homo SNP |
| chr3 | 13794450 | 13794869 | het SNP | het SNP | het SNP |
| chr3 | 14199881 | 14199881 | het SNP | het SNP | het SNP |
| chr3 | 14449540 | 14449640 | homo SNP | homo SNP | homo SNP |
| chr3 | 15396806 | 15396906 | homo SNP | homo SNP | homo SNP |
| chr3 | 20015222 | 20015322 | ND | ND | het SNP |
| chr3 | 23186204 | 23186304 | homo SNP | ND | ND |
| chr4 | 944800 | 944900 | homo SNP | homo SNP | homo SNP |
| chr4 | 3268734 | 3268834 | het SNP | het SNP | het SNP |
| chr4 | 3953947 | 3954227 | het SNP | het SNP | het SNP |
| chr4 | 4025667 | 4025767 | het SNP | het SNP | het SNP |

| | | | | | | |
|---|---|---|---|---|---|---|
| chr4 | 6179688 | 6179789 | het SNP | het SNP | het SNP | |
| chr4 | 7629083 | 7629183 | ND | het SNP | ND | |
| chr4 | 8178664 | 8178764 | het SNP | ND | ND | |
| chr4 | 8195000 | 8196007 | homo InDel | het InDel | homo InDel | PDS3 on-target |
| chr4 | 9457406 | 9457506 | ND | het SNP | homo SNP | |
| chr4 | 10202091 | 10203301 | homo InDel | het InDel | homo InDel | CHLI1 on-target |
| chr4 | 10939399 | 10939499 | homo SNP | ND | ND | |
| chr4 | 15195953 | 15196053 | ND | het SNP | ND | |
| chr4 | 16978520 | 16978620 | ND | het SNP | ND | |
| chr5 | 4786957 | 4787057 | ND | het SNP | het SNP | |
| chr5 | 10002510 | 10002610 | homo SNP | homo SNP | homo SNP | |
| chr5 | 11784970 | 11785070 | homo SNP | homo SNP | homo SNP | |
| chr5 | 13657156 | 13657256 | ND | het SNP | homo SNP | |
| chr5 | 15163048 | 15163154 | homo SNP | homo SNP | homo SNP | |
| chr5 | 19033499 | 19033599 | ND | ND | homo SNP | |
| chr5 | 19492943 | 19493043 | het SNP | ND | ND | |
| chr5 | 21434792 | 21434892 | ND | homo SNP | het SNP | |
| chr5 | 22109092 | 22109192 | ND | ND | het SNP | |
| chr5 | 24819665 | 24819765 | het SNP | ND | ND | |

Variants remaining after filtering against wild-type control samples, excluding sites with sequencing depth below 30x and manually inspected to remove calling artifacts. **Chromosome, Start, End**: genomic coordinates of the variant. **Line 126, Line 153, Line 176**: genotype call for each progeny line (homo SNP: homozygous SNP; het SNP: heterozygous SNP; het InDel: heterozygous insertion-deletion; homo InDel: homozygous deletion; ND: not detected). **On-target**: annotated if the locus is the on-target site.

#### Table S14. Plasmid vectors used in this study (related to Methods)

| Internal ID | System | Purpose |
|---|---|---|
| pZZ09 | *E.coli* | Bacterial expression of Ymu1 protein |
| pHS355 | *E.coli* | Bacterial expression of Dra2 protein |

| | | |
|---|---|---|
| pZZ049 | *E.coli* | Bacterial expression of Tel2 protein |
| pZZ034 | *E.coli* | Bacterial expression of Tfu1 protein |
| pZZ053 | *E.coli* | Bacterial expression of Ec41 protein |
| pZZ031 | *E.coli* | Bacterial expression of dYmu1 protein |
| pZZ069 | *E.coli* | Bacterial expression of V305R Ymu1 protein |
| pZZ080 | *E.coli* | Bacterial expression of H4W Ymu1 protein |
| pZZ087 | *E.coli* | Bacterial expression of H4W dYmu1 protein |
| pHS749 | *E.coli* | Bacterial expression of Ymu1-WFR protein |
| pHS750 | *E.coli* | Bacterial expression of dYmu1-WFR protein |
| pZZ089 | *E.coli* | Bacterial expression of L304F Ymu1 protein |
| pHS607 | *E.coli* | Bacterial expression of Ymu1 200-nt reRNA for target 1 |
| pHS610 | *E.coli* | Bacterial expression of Ymu1 127-nt reRNA for target 1 |
| pHS612 | *E.coli* | Bacterial expression of Ymu1 short reRNA for target 1 |
| pZZ062 | *E.coli* | Bacterial expression of Ymu1 short reRNA for target 2 |
| pZZ064 | *E.coli* | Bacterial expression of Ymu1 127-nt reRNA for target 2 |
| pZZ013 | *E.coli* | Bacterial expression of Dra2 200-nt reRNA for target 1 |
| pZZ052 | *E.coli* | Bacterial expression of Tel2 200-nt reRNA for target 1 |
| pZZ035 | *E.coli* | Bacterial expression of Tfu1 200-nt reRNA for target 1 |
| pZZ054 | *E.coli* | Bacterial expression for Ec41 200-nt reRNA for target 1 |
| pHS516 | *E.coli* | Bacterial TAM assay vector encoding WT Ymu1 and WT reRNA expressed as a single transcript, with a 16-nt guide flanked by HDV |
| pHS751 | *E.coli* | Bacterial TAM assay vector encoding Ymu1-WFR and WT reRNA expressed as a single transcript, with a 16-nt guide flanked by HDV |
| pHS697 | *E.coli* | Substrate plasmid for cleavage assay encoding TAM and Target 1 |
| pTW2036 | *Arabidopsis* | WT Ymu1 and WT reRNA with 14 nt spacer length targeting AtPDS3_g2 |
| pTW2035 | *Arabidopsis* | WT Ymu1 and WT reRNA with 15 nt spacer length targeting AtPDS3_g2 |
| pMK061 | *Arabidopsis* | WT Ymu1 and WT reRNA with 16 nt spacer length targeting AtPDS3_g2 |
| pTW2034 | *Arabidopsis* | WT Ymu1 and WT reRNA with 17 nt spacer length targeting AtPDS3_g2 |
| pTW2033 | *Arabidopsis* | WT Ymu1 and WT reRNA with 18 nt spacer length targeting AtPDS3_g2 |

| | | |
|---|---|---|
| pTW2032 | *Arabidopsis* | WT Ymu1 and WT reRNA with 19 nt spacer length targeting AtPDS3_g2 |
| pTW2031 | *Arabidopsis* | WT Ymu1 and WT reRNA with 20 nt spacer length targeting AtPDS3_g2 |
| pTW2314 | *Arabidopsis* | G285A Ymu1 variant with short reRNA targeting AtPDS3_g2 |
| pTW2315 | *Arabidopsis* | G285I Ymu1 variant with short reRNA targeting AtPDS3_g2 |
| pTW2317 | *Arabidopsis* | G285V Ymu1 variant with short reRNA targeting AtPDS3_g2 |
| pTW2321 | *Arabidopsis* | H4W-L304F Ymu1 variant with short reRNA targeting AtPDS3_g2 |
| pTW2322 | *Arabidopsis* | H4W-L304F-V305R Ymu1 variant with short reRNA targeting AtPDS3_g2 |
| pTW2397 | *Arabidopsis* | H4W-V305R Ymu1 variant with short reRNA targeting AtPDS3_g2 |
| pTW2335 | *Arabidopsis* | L304F-V305R Ymu1 variant with short reRNA targeting AtPDS3_g2 |
| pTW2145 | *Arabidopsis* | WT Ymu1 with WT reRNA targeting AtCHLI1_g4 |
| pTW2149 | *Arabidopsis* | WT Ymu1 with WT reRNA targeting AtCHLI1_g6 |
| pTW2125 | *Arabidopsis* | WT Ymu1 with WT reRNA targeting AtCHLI1_g9 |
| pTW2126 | *Arabidopsis* | WT Ymu1 with WT reRNA targeting AtCHLI2_g1 |
| pTW2128 | *Arabidopsis* | WT Ymu1 with WT reRNA targeting AtCHLI2_g3 |
| pTW2147 | *Arabidopsis* | WT Ymu1 with WT reRNA targeting AtCHLI2_g8 |
| pTW2134 | *Arabidopsis* | WT Ymu1 with WT reRNA targeting AtCHLI2_g10 |
| pTW2198 | *Arabidopsis* | WT Ymu1 with WT reRNA targeting AtCPC_g1 |
| pTW2202 | *Arabidopsis* | WT Ymu1 with WT reRNA targeting AtCPC_g5 |
| pMK064 | *Arabidopsis* | WT Ymu1 with WT reRNA targeting AtPDS3_g5 |
| pMK068 | *Arabidopsis* | WT Ymu1 with WT reRNA targeting AtPDS3_g10 |
| pMK070 | *Arabidopsis* | WT Ymu1 with WT reRNA targeting AtPDS3_g12 |
| pTW2196 | *Arabidopsis* | WT Ymu1 with WT reRNA targeting AtTRY_g3 |
| pTW2541 | *Arabidopsis* | H4W-L304F-V305R Ymu1 with WT reRNA targeting AtPDS3_g2 |
| pTW2532 | *Arabidopsis* | H4W-L304F-V305R Ymu1 with WT reRNA targeting AtCHLI1_g4 |
| pTW2533 | *Arabidopsis* | H4W-L304F-V305R Ymu1 with WT reRNA targeting AtCHLI1_g6 |
| pTW2534 | *Arabidopsis* | H4W-L304F-V305R Ymu1 with WT reRNA targeting AtCHLI1_g9 |
| pTW2536 | *Arabidopsis* | H4W-L304F-V305R Ymu1 with WT reRNA targeting AtCHLI2_g8 |
| pTW2537 | *Arabidopsis* | H4W-L304F-V305R Ymu1 with WT reRNA targeting AtCHLI2_g10 |

| | | |
|---|---|---|
| pTW2542 | *Arabidopsis* | H4W-L304F-V305R Ymu1 with WT reRNA targeting AtPDS3_g5 |
| pTW2543 | *Arabidopsis* | H4W-L304F-V305R Ymu1 with WT reRNA targeting AtPDS3_g10 |
| pTW2471 | *Arabidopsis* | H4W Ymu1with WT reRNA targeting AtPDS3_g2 |
| pTW2503 | *Arabidopsis* | H4W-L304F-V305R Ymu1 with WT reRNA ccdb gRNA cloning vector |
| pTW2453 | *Arabidopsis* | H4W Ymu1 with WT reRNA ccdb gRNA cloning vector |
| pMK525 | *Arabidopsis* | ccdb TnpB cloning vector |
| pMK025 | *Arabidopsis* | WT Ymu1 with WT reRNA ccdb gRNA cloning vector |
| pKV100 | *Arabidopsis* | H4Y Ymu1 variant with short reRNA targeting AtPDS3_g2 |
| pKV101 | *Arabidopsis* | H4F Ymu1 variant with short reRNA targeting AtPDS3_g2 |
| pKV102 | *Arabidopsis* | H4W Ymu1 variant with short reRNA targeting AtPDS3_g2 |
| pKV106 | *Arabidopsis* | K229A Ymu1 variant with short reRNA targeting AtPDS3_g2 |
| pKV105 | *Arabidopsis* | K229E Ymu1 variant with short reRNA targeting AtPDS3_g2 |
| pKV111 | *Arabidopsis* | K229I Ymu1 variant with short reRNA targeting AtPDS3_g2 |
| pKV112 | *Arabidopsis* | K229L Ymu1 variant with short reRNA targeting AtPDS3_g2 |
| pKV104 | *Arabidopsis* | K229Q Ymu1 variant with short reRNA targeting AtPDS3_g2 |
| pKV103 | *Arabidopsis* | K229R Ymu1 variant with short reRNA targeting AtPDS3_g2 |
| pKV113 | *Arabidopsis* | K229V Ymu1 variant with short reRNA targeting AtPDS3_g2 |
| pKV110 | *Arabidopsis* | R230A Ymu1 variant with short reRNA targeting AtPDS3_g2 |
| pKV109 | *Arabidopsis* | R230E Ymu1 variant with short reRNA targeting AtPDS3_g2 |
| pKV114 | *Arabidopsis* | R230I Ymu1 variant with short reRNA targeting AtPDS3_g2 |
| pKV107 | *Arabidopsis* | R230K Ymu1 variant with short reRNA targeting AtPDS3_g2 |
| pKV115 | *Arabidopsis* | R230L Ymu1 variant with short reRNA targeting AtPDS3_g2 |
| pKV108 | *Arabidopsis* | R230Q Ymu1 variant with short reRNA targeting AtPDS3_g2 |
| pKV116 | *Arabidopsis* | R230V Ymu1 variant with short reRNA targeting AtPDS3_g2 |
| pKV136 | *Arabidopsis* | V283E Ymu1 variant with short reRNA targeting AtPDS3_g2 |
| pKV138 | *Arabidopsis* | V283I Ymu1 variant with short reRNA targeting AtPDS3_g2 |
| pKV141 | *Arabidopsis* | V283K Ymu1 variant with short reRNA targeting AtPDS3_g2 |
| pKV140 | *Arabidopsis* | V283L Ymu1 variant with short reRNA targeting AtPDS3_g2 |
| pKV137 | *Arabidopsis* | V283Q Ymu1 variant with short reRNA targeting AtPDS3_g2 |
| pKV134 | *Arabidopsis* | V283R Ymu1 variant with short reRNA targeting AtPDS3_g2 |

| | | |
|---|---|---|
| pKV135 | *Arabidopsis* | V283Y Ymu1 variant with short reRNA targeting AtPDS3_g2 |
| pKV139 | *Arabidopsis* | M287R Ymu1 variant with short reRNA targeting AtPDS3_g2 |
| pKV128 | *Arabidopsis* | H290A Ymu1 variant with short reRNA targeting AtPDS3_g2 |
| pKV129 | *Arabidopsis* | H290E Ymu1 variant with short reRNA targeting AtPDS3_g2 |
| pKV126 | *Arabidopsis* | H290R Ymu1 variant with short reRNA targeting AtPDS3_g2 |
| pKV127 | *Arabidopsis* | H290Y Ymu1 variant with short reRNA targeting AtPDS3_g2 |
| pKV132 | *Arabidopsis* | A293E Ymu1 variant with short reRNA targeting AtPDS3_g2 |
| pKV133 | *Arabidopsis* | A293Q Ymu1 variant with short reRNA targeting AtPDS3_g2 |
| pKV130 | *Arabidopsis* | A293R Ymu1 variant with short reRNA targeting AtPDS3_g2 |
| pKV131 | *Arabidopsis* | A293Y Ymu1 variant with short reRNA targeting AtPDS3_g2 |
| pKV120 | *Arabidopsis* | S303F Ymu1 variant with short reRNA targeting AtPDS3_g2 |
| pKV118 | *Arabidopsis* | S303I Ymu1 variant with short reRNA targeting AtPDS3_g2 |
| pKV124 | *Arabidopsis* | S303K Ymu1 variant with short reRNA targeting AtPDS3_g2 |
| pKV117 | *Arabidopsis* | S303L Ymu1 variant with short reRNA targeting AtPDS3_g2 |
| pKV125 | *Arabidopsis* | S303M Ymu1 variant with short reRNA targeting AtPDS3_g2 |
| pKV123 | *Arabidopsis* | S303R Ymu1 variant with short reRNA targeting AtPDS3_g2 |
| pKV119 | *Arabidopsis* | S303V Ymu1 variant with short reRNA targeting AtPDS3_g2 |
| pKV122 | *Arabidopsis* | S303W Ymu1 variant with short reRNA targeting AtPDS3_g2 |
| pKV121 | *Arabidopsis* | S303Y Ymu1 variant with short reRNA targeting AtPDS3_g2 |
| pKV158 | *Arabidopsis* | L304K Ymu1 variant with short reRNA targeting AtPDS3_g2 |
| pKV159 | *Arabidopsis* | L304R Ymu1 variant with short reRNA targeting AtPDS3_g2 |
| pKV160 | *Arabidopsis* | L304F Ymu1 variant with short reRNA targeting AtPDS3_g2 |
| pKV161 | *Arabidopsis* | V305R Ymu1 variant with short reRNA targeting AtPDS3_g2 |
| pKV162 | *Arabidopsis* | V305F Ymu1 variant with short reRNA targeting AtPDS3_g2 |
| pKV87 | *Arabidopsis* | WT Ymu1 with short reRNA targeting AtPDS3_g2 |
| pHS550 | *Arabidopsis* | mRFP drop out vector with backbones for protoplast assay |

Plasmids pHS516, pMK025, pMK061, pMK64, pMK68, pMK70 are from a previous study[S13].

**Table S15. Primers used for Gibson assembly-based site-directed mutagenesis of Ymu1 TnpB for bacterial expression (related to Methods)**

| Oligonucleotide | Internal ID | Sequence (5′-3′) |
|---|---|---|

| | | |
|---|---|---|
| Reverse primer for amplifying Ymu1 backbone plasmid | oZZ080R | ttgacggcttgacggagtagcatagggtttgcag |
| Forward primer for amplifying Ymu1 backbone plasmid | oZZ081F | gcagggattctgcaaaccctatgctactccgtc |
| Forward primer to induce the E279A mutation in Ymu1 | oZZ086F | AAACCACGATATCATCTGTATCgcgGACCTTAACG TTAAGGGCATGATGC |
| Reverse primer to induce the E279A mutation in Ymu1 | oZZ087R | GCCCTTAACGTTAAGGTCcgcGATACAGATGATAT CGTGGTTTTTGACTATCTCTGTAC |
| Reverse primer to induce the H4W mutation in Ymu1 | oZZ227R | GTATTCATAGGCTTTccaCTGCAGCATTGCATTGG ATTGG |
| Forward primer to induce the H4W mutation in Ymu1 | oZZ228F | AATGCAATGCTGCAGtggAAAGCCTATGAATACCG TATCTATCCAGATAAGAAG |
| Forward primer to induce the both L304F/V305R mutations in Ymu1 | oZZ233F | TGGACGAGCttccgaTCGAAACTGCAGTACAAGGCT TC |
| Reverse primer to induce the both L304F/V305R mutations in Ymu1 | oZZ234R | CAGTTTCGAtcggaaGCTCGTCCATGATACATCAGA GATGC |
| Reverse primer to induce the L304F mutation in Ymu1 | oZZ235R | CTGCAGTTTCGATACgaaGCTCGTCCATGATACAT CAGAGATGC |
| Forward primer to induce the L304F mutation in Ymu1 | oZZ236F | GTATCATGGACGAGCttcGTATCGAAACTGCAGTA CAAGGCTTC |
| Forward primer to induce the V305R mutation in Ymu1 | oZZ217F | TCATGGACGAGCCTTagaTCGAAACTGCAGTACAA GGCTTC |
| Reverse primer to induce the V305R mutation in Ymu1 | oZZ220R | GTACTGCAGTTTCGAtctAAGGCTCGTCCATGATA CATCAGAG |

**Table S16. Amino acid sequence of the Ymu1 TnpB bacterial expression construct (related to Methods)**

| ID | Protein sequence |
|---|---|
| WT Ymu1 | MKSSHHHHHHHHHHGSSMKIEEGKLVIWINGDKGYNGLAEVGKKFEKDTGIKVTV EHPDKLEEKFPQVAATGDGPDIIFWAHDRFGGYAQSGLLAEITPDKAFQDKLYPFT WDAVRYNGKLIAYPIAVEALSLIYNKDLLPNPPKTWEEIPALDKELKAKGKSALMFN LQEPYFTWPLIAADGGYAFKYENGKYDIKDVGVDNAGAKAGLTFLVDLIKNKHMNA DTDYSIAEAAFNKGETAMTINGPWAWSNIDTSKVNYGVTVLPTFKGQPSKPFVGVL SAGINAASPNKELAKEFLENYLLTDEGLEAVNKDKPLGAVALKSYEEELAKDPRIAA TMENAQKGEIMPNIPQMSAFWYAVRTAVINAASGRQTVDEALKDAQTNSSSNNNN NNNNNNLGIEENLYFQSNAMLQHKAYEYRIYPDKKQETLIAKTIGSSRYVYNHFLEL WNKEYEETGKGLTYYACSKLLTKLKRDPETVWLCEVDKFSLQNSLRNLSDAFSRF FKGQNEHPQFKSKKSPRQSYTTQYTNNNIAVSGNCLKLPKLGLVKFADSREMKGR ILNATVRRKSSGKFFVSILCEEEICELPKTDSSVGIDLGIIDFAVMSDGSRHDNNHFT |

| | |
|---|---|
| | RQMEERLRREQRKLARRALAAEKRGISLSEARNYQKQRRKVARLYEKVANQRKE YLNKLSTEIVKNHDIICIEDLNVKGMMRNHKLAKSISDVSWTSLVSKLQYKASWYGK EVIRISRWFPSSQICSECGHKDRKKPLHVREWTCPVCHAHHDRDVNAARNILAEGL RIRALTPGS |

10 His-tags; MBP; TEV site; TnpB protein

**Table S17. DNA sequence of the 127-nt reRNA expression cassette (related to Methods)**

| ID | DNA sequence (5′-3′) |
|---|---|
| 127-nt reRNA | TAATACGACTCACTATAGGCAAACAGGAACCGCAGGAATTGCGGGGGTAGC TTGGTAAACAAGAGAAACCTCTGCCGGCAAAGAAATAAGCCGGTAAGTATGC TCTGTTCCCAAGAATCTCGTGACTTTAGTCATGAGAGTTTCAATCTTCTGGAT TGTTGTGGCCGGCATGGTCCCAGCCTCCTCGCTGGCGCCGGCTGGGCAAC ATGCTTCGGCATGGCGAATGGGAC |

T7 promoter; reRNA scaffold; guide sequence (for Target 1); HDV ribozyme

**Table S18. DNA sequence of plasmid cleavage substrate (related to Methods)**

| pHS697 |
|---|
| CGAAAAATCAATAATCAGACAACAAGATGTGCGAACTCGATATTTTACACGACTCTCTTTA CCAATTCTGCCCCGAATTACACTTAAAACGACTCAACAGCTTAACGTTGGCTTGCCACGC ATTACTTGACTGTAAAACTCTCACTCTTACCGAACTTGGCCGTAACCTGCCAACCAAAGC GAGAACAAAACATAACATCAAACGAATCGACCGATTGTTAGGTAATCGTCACCTGCAGGA AGGTTTAAACGCATTTAGGTGACACTATAGAAGTGTGTATCGCTCGAGGGATCCGAATTC GAAGTCTTGGTACGGAGCGAGACCGGAGGTCATAATGATTGAT**TCTTCTGGATTGTTGT**A AGCAGCATTTGAGCAAAAATCTGTTGCCCATGGTCTCACCATTCCTGTAGACTTCTTAATT AAGACGTCAGAATTCTCGAGGCGGCCGCATGTGAGTCTCCCTATAGTGAGTCGTATTAAT TTCGCGGGCGGAACCCCTATTTGTTTATTTTTCTAAATACATTCAAATATGTATCCGCTCA TGAGTAGCACCAGGCGTTTAAGGGCACCAATAACTGCCTTAAAAAAATTACGCCCCGCC CTGCCACTCATCGCAGTACTGTTGTAATTCATTAAGCATTCTGCCGACATGGAAGCCATC ACAAACGGCATGATGAACCTGAATCGCCAGCGGCATCAGCACCTTGTCGCCTTGCGTAT AATATTTGCCCATGGTGAAAACGGGGGCGAAGAAGTTGTCCATATTGGCCACGTTTAAAT CAAAACTGGTGAAACTCACCCAGGGATTGGCTGAGACAAAAAACATATTCTCAATAAACC CTTTAGGGAAATAGGCCAGGTTTTCACCGTAACACGCCACATCTTGCGAATATATGTGTA GAAACTGCCGGAAATCGTCGTGGTATTCACTCCAGAGCGATGAAAACGTTTCAGTTTGCT CATGGAAAACGGTGTAACAAGGGTGAACACTATCCCATATCACCAGCTCACCGTCTTTCA TTGCCATACGAAATTCCGGATGAGCATTCATCAGGCGGGCAAGAATGTGAATAAAGGCC GGATAAAACTTGTGCTTATTTTTCTTTACGGTCTTTAAAAAGGCCGTAATATCCAGCTGAA CGGTCTGGTTATAGGTACATTGAGCAACTGACTGAAATGCCTCAAAATGTTCTTTACGAT GCCATTGGGATATATCAACGGTGGTATATCCAGTGATTTTTTTCTCCATTTTAGCTTCCTT AGCTCCTGAAAATCTCGATAACTCAAAAAATACGCCCGGTAGTGATCTTATTTCATTATGG TGAAAGTTGGAACCTCTTACGTGCCGATCAAAGTCTCATTTTCGCCAAAAGTTGTCATGA CCAAAATCCCTTAACGTGAGTTTTCGTTCCACTGAGCGTCAGACCCCGTAGAAAAGATCA AAGGATCTTCTTGAGATCCTTTTTTTCTGCGCGTAATCTGCTGCTTGCAAACAAAAAAACC ACCGCTACCAGCGGTGGTTTGTTTGCCGGATCAAGAGCTACCAACTCTTTTTCCGAAGGT AACTGGCTTCAGCAGAGCGCAGATACCAAATACTGTTCTTCTAGTGTAGCCGTAGTTAGG CCACCACTTCAAGAACTCTGTAGCACCGCCTACATACCTCGCTCTGCTAATCCTGTTACC AGTGGCTGCTGCCAGTGGCGATAAGTCGTGTCTTACCGGGTTGGACTCAAGACGATAGT |

| |
|---|
| TACCGGATAAGGCGCAGCGGTCGGGCTGAACGGGGGGTTCGTGCACACAGCCCAGCTT GGAGCGAACGACCTACACCGAACTGAGATACCTACAGCGTGAGCTATGAGAAAGCGCCA CGCTTCCCGAAGGGAGAAAGGCGGACAGGTATCCGGTAAGCGGCAGGGTCGGAACAG GAGAGCGCACGAGGGAGCTTCCAGGGGGAAACGCCTGGTATCTTTATAGTCCTGTCGG GTTTCGCCACCTCTGACTTGAGCGTCGATTTTTGTGATGCTCGTCAGGGGGGCGGAGCC TATGGAAAAACGCCAGCAATGCGGCCTTTTTACGGTTCCTGGCCTTTTGCTGGCCTTTTG CTCACATGTTCTTTCCTGCGTTATCCCCTGATTCTGTGGATAACCGTATTACCGCCTTTGA GTGAGCTGATACCGCTCGCCGCAGCCGAACGACCGAGCGCAGCGAGTCAGTGAGCGA GGAAGC |

**bolded** region represents the 16-nt Target 1 sequence (the TS sequence);
yellow highlighted region represents the TAM (the complementary sequence of 5′-TTGAT-3′);

**Table S19. Sequences of all the oligonucleotide substrates used in DNA cleavage assays (related to Methods)**

| Internal ID | Description | Sequence (5′-3′) | Figures |
|---|---|---|---|
| dZZ01 | Ymu1 DNA substrate 0-16 bp corresponding to target 1; NTS; Cy5 | /5Cy5/CCGGAGGTCATAATGAT TGATTCTTCTGGATTGTTGTAA GCAGCATTTGAGCAAAAATCT | Fig. 3C-F, 5J, S1E-F, S3E-N, S3Q-R, S5H |
| dZZ01C | Ymu1 DNA substrate 2-16 bp corresponding to target 1 (substrate pre-unwound at bases 1 and 2) ; NTS; Cy5 | /5Cy5/CCGGAGGTCATAATGAT TGATAGTTCTGGATTGTTGTAA GCAGCATTTGAGCAAAAATCT | Fig. 5J, S5H |
| dZZ01F | Ymu1 DNA substrate 0-7 and 10-16 bp corresponding to target 1 (substrate pre-unwound at bases 8 and 9); NTS; Cy5 | /5Cy5/CCGGAGGTCATAATGAT TGATAGAAGACCTTTGTTGTAA GCAGCATTTGAGCAAAAATCT | Fig. 5J, S5H |
| dZZ01G | Ymu1 DNA substrate 10-16 bp corresponding to target 1 (substrate pre-unwound from bases 1 to 9); NTS; Cy5 | /5Cy5/CCGGAGGTCATAATGAT TGATTCTTCTGCCTTGTTGTAA GCAGCATTTGAGCAAAAATCT | Fig. 5J, S5H |
| dZZ02 | Ymu1 DNA substrate 0-16 bp corresponding to target 1; TS; FAM | /56-FAMN/AGATTTTTGCTCAAATG CTGCTTACAACAATCCAGAAG AATCAATCATTATGACCTCCGG | Fig. 3C-F, 5J, S1E-F, S3E-N, S3Q-R, S5H |
| dZZ013 | Ymu1 DNA substrate 0-16 bp corresponding to target 2; NTS; Cy5 | /5Cy5/CCGGAGGTCATAATGAT TGATAAGGCAAATTCGCCGCA AGCAGCATTTGAGCAAAAATC T | Fig. 3G-H, S3O-P |
| dZZ014 | Ymu1 DNA substrate 0-16 bp corresponding to target 2; TS; FAM | /56-FAMN/AGATTTTTGCTCAAATG CTGCTTGCGGCGAATTTGCCT TATCAATCATTATGACCTCCGG | Fig. 3G-H, S3O-P |

| | | | |
|---|---|---|---|
| dZZ017 | Ymu1 DNA substrate with the 4th base mismatched from the target 1 guide; NTS; Cy5 | /5Cy5/CCGGAGGTCATAATGAT TGATTCTACTGGATTGTTGTAA GCAGCATTTGAGCAAAAATCT | S5J-K |
| dZZ018 | Ymu1 DNA substrate with the 4th base mismatched from the target 1 guide; TS; Cy2 | /56-FAMN/AGATTTTTGCTCAAATG CTGCTTACAACAATCCAGTAG AATCAATCATTATGACCTCCGG | S5J-K |
| dZZ025 | Ymu1 DNA substrate with the 7th base mismatched from the target 1 guide; NTS; Cy5 | /5Cy5/CCGGAGGTCATAATGAT TGATTCTTCTCGATTGTTGTAA GCAGCATTTGAGCAAAAATCT | S5J-K |
| dZZ026 | Ymu1 DNA substrate with the 7th base mismatched from the target 1 guide; TS; Cy2 | /5Cy5/CCGGAGGTCATAATGAT TGATTCTTCTCGATTGTTGTAA GCAGCATTTGAGCAAAAATCT | S5J-K |
| dZZ033 | Ymu1 DNA substrate with the 10th base mismatched from the target 1 guide; NTS; Cy5 | /5Cy5/CCGGAGGTCATAATGAT TGATTCTTCTGGAATGTTGTAA GCAGCATTTGAGCAAAAATCT | S5J-K |
| dZZ034 | Ymu1 DNA substrate with the 10th base mismatched from the target 1 guide; TS; Cy2 | /56-FAMN/AGATTTTTGCTCAAATG CTGCTTACAACATTCCAGAAG AATCAATCATTATGACCTCCGG | S5J-K |
| dZZ029 | Ymu1 DNA substrate with the 14th base mismatched from the target 1 guide; NTS; Cy5 | /5Cy5/CCGGAGGTCATAATGAT TGATTCTTCTGGATTGTAGTAA GCAGCATTTGAGCAAAAATCT | S5J-K |
| dZZ030 | Ymu1 DNA substrate with the 14th base mismatched from the target 1 guide; TS; Cy2 | /56-FAMN/AGATTTTTGCTCAAATG CTGCTTACTACAATCCAGAAG AATCAATCATTATGACCTCCGG | S5J-K |

**Table S20. Primer sequences used to generate DNA tether fragments in AuRBT (related to Fig. S4 and Methods).**

| | Primer Sequence (5′-3′) | Length | Template | Digest |
|---|---|---|---|---|
| F500 | gatcgaagacacttagACAACCCACAAGTATAGAGGCTCCTATG<br><br>GACGCGGATATAATGACATTTCCTAAC | 520 | pFO-SE2 | BbsI |
| XD | gaagggtctcatgacTCACTATAGGGCGAATTGGAGCTCCACCGCG<br><br>gaagggtctcactaaCACTAAAGGGAACAAAAGCTGGTAC | 4165 | pFO-SE2 | BsaI |
| SOI | gatcgaagacacctgcTGACGCATCAGTCAGTACTACTGACG<br><br>gatcgaagacacgtcaGTGGTAGCTGCTGTGCGAGTG | 290 | TnpB Gblock | BbsI |
| SOI 1-4MM | gatcgaagacacctgcTGACGCATCAGTCAGTACTACTGACG<br><br>gatcgaagacacgtcaGTGGTAGCTGCTGTGCGAGTG | 290 | TnpB 1-4MMGblock | BbsI |

| | | | | |
|---|---|---|---|---|
| SPMDIG | GCGCAGCACGCAGATACACTC<br><br>gatcgaagacacgcagCATACGTCTGCGTCGCTG | 320 | C12Gblock | BbsI |

Biotin-modified nucleotides highlighted in blue.

**Table S21. Target sequences in the SOI (related to Fig. S4 and Methods).**

| DNA Target | Sequence (5′-3′) |
|---|---|
| Target 1 | TTGATTCTTCTGGATTGTTGT |
| 1-4 mismatch | TTGATAGAACTGGATTGTTGT |

TAM sequence highlighted in yellow.

**Table S22. The full DNA tether sequence with Target 1 (related to Fig. S4 and Methods)**

| Sequence (5′-3′) |
|---|
| GACGCGGATATAATGACATTTCCTAACTTTTGGGCAAAAATTCGCTATCATATGCGAGAACCGTTT<br>GCGGAGTTTCTCGGGACACTAGTTCTTGTCATTTTTGGTGTTGGTGGTAATCTTCAAGCAACTGTA<br>ACAAAAGGTAGTGGTGGTTCCTATGAATCCCTATCATTTGCATGGGGGTTCGGTTGTATGCTTGGT<br>GTTTACGTCGCAGGCGGTATTAGTGGTGGTCATATTAACCCTGCTGTTACGATTTCAATGGCAATT<br>TTTCGAAAATTCCCCTGGAAAAAGGTGCCCGTATATATTGTTGCTCAGATTATCGGTGCATATTTTG<br>GAGGAGCTATGGCTTATGGTTATTTTTGGAGCTCTATCACAGAATTTGAGGGAGGTCCGCACATAA<br>GAACAACGGCGACCGGTGCGTGTTTGTTTACTGATCCAAAGTCTTACGTCACGTGGAGAAATGCC<br>TTCTTTGACGAATTCATAGGAGCCTCTATACTTGTGGGTTGTCTAACACTAAAGGGAACAAAAGCT<br>GGTACCGGGCCCCCCCTCGAGCGGTACCCCACTTACCCACCCCGGAAATTTGAGTTATAAACGTT<br>GTTTGAGCTTTACCTAGTCTTGGTCGATCAAAAGTTCTGGTACCTTTTCACCATGTCTCCCCCCTTA<br>TTCATATAAAAAGAAGCGTATAATCGCACAGTATAACGCTCCTCTGATATATGATCTAGACCCAAGT<br>AATGAGTTACGAATCTGGGAGGTCATCCTCCTCTTCCGAGAGTACACGGCCACCAACGCTAAAAG<br>AAGAACCTAATGGTAAAATAGCTTGGGAAGAAAGTGTCAAAAAATCTAGGGAAAATAACGAAAATG<br>ACAGCACTCTCTTGAGGCGAAAGCTAGGTGAGACTCGAAAAGCAATTGAAACTGGAGGATCATCG<br>AGAAATAAACTTTCTGCTTTGACACCCTTGAAAAAAGTGGTTGACGAGAGGAAGGATTCGGTACAA<br>CCACAGGTCCCTTCCATGGGTTTTACTTATTCTTTGCCTAATTTGAAGACTTTAAACAGTTTTTCAGA<br>TGCTGAGCAAGCACGTATAATGCAAGATTATCTATCCAGGGGGGTAAATCAAGGCAACAGTAATAA<br>TTATGTAGACCCACTATATCGGCAATTAAATCCAACTATGGGTAGTAGCAGGAACAGGCCTGTTTG<br>GAGTTTAAATCAGCCGTTACCGCATGTATTGGATCGAGGCTTGGCAGCAAAGATGATACAAAAGAA<br>TATGGATGCAAGGTCCCGCGCATCATCGAGACGAGGGTCGACCGATATTTCAAGGGGGGGTTCTA<br>CTACGTCAGTGAAAGACTGGAAAAGGCTCCTTAGAGGTGCAGCACCGGGTAAAAAGCTTGGTGAC<br>ATCGAAGCTCAAACGCAACGCGATAATACTGTTGGTGCAGATGTGAAACCTACTAAGTTAGAGCCT<br>GAAAACCCACAAAAGCCCTCTAACACGCATATTGAGAATGTTTCACGTAAGAAAAAGCGTACTTCG<br>CATAATGTCAATTTTTCATTAGGCGATGAAAGCTACGCATCCTCCATAGCCGATGCAGAATCCAGA<br>AAATTAAAGAACATGCAAACCCTCGATGGTTCTACTCCGGTTTATACGAAGCTTCCTGAAGAACTTA<br>TTGAAGAGGAAAATAAAAGTACGAGTGCATTAGATGGTAATGAAATTGGTGCCTCAGAAGATGAAG<br>ACGCGGATATAATGACATTTCCTAACTTTTGGGCAAAAATTCGCTATCATATGCGAGAACCGTTTGC<br>GGAGTTTCTCGGGACACTAGTTCTTGTCATTTTTGGTGTTGGTGGTAATCTTCAAGCAACTGTAACA<br>AAAGGTAGTGGTGGTTCCTATGAATCCCTATCATTTGCATGGGGGTTCGGTTGTATGCTTGGTGTT<br>TACGTCGCAGGCGGTATTAGTGGTGGTCATATTAACCCTGCTGTTACGATTTCAATGGCAATTTTT<br>CGAAAATTCCCCTGGAAAAAGGTGCCCGTATATATTGTTGCTCAGATTATCGGTGCATATTTTGGA<br>GGAGCTATGGCTTATGGTTATTTTTGGAGCTCTATCACAGAATTTGAGGGAGGTCCGCACATAAGA<br>ACAACGGCGACCGGTGCGTGTTTGTTTACTGATCCAAAGTCTTACGTCACGTGGAGAAATGCCTTC<br>TTTGACGAATTCATAGGAGCCTCTATACTTGTGGGTTGTTTGATGGCGCTATTGGATGATAGTAAT<br>GCTCCACCTGGCAATGGTATGACCGCATTAATTATTGGATTCTTAGTCGCTGCAATTGGTATGGCC<br>CTTGGATATCAAACAAGTTTCACAATCAATCCTGCAAGAGATCTCGGTCCTCGCATATTTGCTTCCA<br>TGATTGGCTATGGTCCACATGCTTTTCATCTCACACATTGGTGGTGGACATGGGGAGCCTGGGGT<br>GGTCCAATTGCCGGCGGTATTGCTGGAGCACTCATATATGACATTTTCATTTTTACTGGATGCGAA<br>TCCCCAGTCAACTACCCAGACAACGGTTATATTGAGAATAGGGTAGGCTGCAGCCCGGGGGATCC<br>ACTAGTTCTAGAGCGGCCGCCGGTACCCCACTTACCCACCCCGGAAATTTGAGTTATAAACGTTGT |

TTGAGCTTTACCTAGTCTTGGTCGATCAAAAGTTCTGGTACCTTTTCACCATGTCTCCCCCCTTATT
CATATAAAAAGAAGCGTATAATCGCACAGTATAACGCTCCTCTGATATATGATCTAGACCCAAGTAA
TGAGTTACGAATCTGGGAGGTCATCCTCCTCTTCCGAGAGTACACGGCCACCAACGCTAAAAGAA
GAACCTAATGGTAAAATAGCTTGGGAAGAAAGTGTCAAAAAATCTAGGGAAAATAACGAAAATGAC
AGCACTCTCTTGAGGCGAAAGCTAGGTGAGACTCGAAAAGCAATTGAAACTGGAGGATCATCGAG
AAATAAACTTTCTGCTTTGACACCCTTGAAAAAAGTGGTTGACGAGAGGAAGGATTCGGTACAACC
ACAGGTCCCTTCCATGGGTTTTACTTATTCTTTGCCTAATTTGAAGACTTTAAACAGTTTTTCAGATG
CTGAGCAAGCACGTATAATGCAAGATTATCTATCCAGGGGGGTAAATCAAGGCAACAGTAATAATT
ATGTAGACCCACTATATCGGCAATTAAATCCAACTATGGGTAGTAGCAGGAACAGGCCTGTTTGGA
GTTTAAATCAGCCGTTACCGCATGTATTGGATCGAGGCTTGGCAGCAAAGATGATACAAAAGAATA
TGGATGCAAGGTCCCGCGCATCATCGAGACGAGGGTCGACCGATATTTCAAGGGGGGGTTCTAC
TACGTCAGTGAAAGACTGGAAAAGGCTCCTTAGAGGTGCAGCACCGGGTAAAAAGCTTGGTGACA
TCGAAGCTCAAACGCAACGCGATAATACTGTTGGTGCAGATGTGAAACCTACTAAGTTAGAGCCTG
AAAACCCACAAAAGCCCTCTAACACGCATATTGAGAATGTTTCACGTAAGAAAAAGCGTACTTCGC
ATAATGTCAATTTTTCATTAGGCGATGAAAGCTACGCATCCTCCATAGCCGATGCAGAATCCAGAA
AATTAAAGAACATGCAAACCCTCGATGGTTCTACTCCGGTTTATACGAAGCTTCCTGAAGAACTTAT
TGAAGAGGAAAATAAAAGTACGAGTGCATTAGATGGTAATGAAATTGGTGCCTCAGAAGATGAAGA
CGCGGATATAATGACATTTCCTAACTTTTGGGCAAAAATTCGCTATCATATGCGAGAACCGTTTGC
GGAGTTTCTCGGGACACTAGTTCTTGTCATTTTTGGTGTTGGTGGTAATCTTCAAGCAACTGTAACA
AAAGGTAGTGGTGGTTCCTATGAATCCCTATCATTTGCATGGGGGTTCGGTTGTATGCTTGGTGTT
TACGTCGCAGGCGGTATTAGTGGTGGTCATATTAACCCTGCTGTTACGATTTCAATGGCAATTTTT
CGAAAATTCCCCTGGAAAAAGGTGCCCGTATATATTGTTGCTCAGATTATCGGTGCATATTTTGGA
GGAGCTATGGCTTATGGTTATTTTTGGAGCTCTATCACAGAATTTGAGGGAGGTCCGCACATAAGA
ACAACGGCGACCGGTGCGTGTTTGTTTACTGATCCAAAGTCTTACGTCACGTGGAGAAATGCCTTC
TTTGACGAATTCATAGGAGCCTCTATACTTGTGGGTTGTTTGATGGCGCTATTGGATGATAGTAAT
GCTCCACCTGGCAATGGTATGACCGCATTAATTATTGGATTCTTAGTCGCTGCAATTGGTATGGCC
CTTGGATATCAAACAAGTTTCACAATCAATCCTGCAAGAGATCTCGGTCCTCGCATATTTGCTTCCA
TGATTGGCTATGGTCCACATGCTTTTCATCTCACACATTGGTGGTGGACATGGGGAGCCTGGGGT
GGTCCAATTGCCGGCGGTATTGCTGGAGCACTCATATATGACATTTTCATTTTTACTGGATGCGAA
TCCCCAGTCAACTACCCAGACAACGGTTATATTGAGAATAGGGTAGGGGCCGCCACCGCGGTGG
AGCTCCAATTCGCCCTATAGTGAGTCAGTGGTAGCTGCTGTGCGAGTGTATCTGCGTGCTGCGCT
AGTGAGATCACGTGCTCATGATACGTGATCTCGTCGCGTAGTGCAGCGAGATCGCGTATGTACTG
CTT**ACAACAATCCAGAAGA**ATCAAGCAGCGATATCTCGTGTGCAGACAGCTCGCGACGATGCGC
TACGTCATCTAGTAGTGCGTACTGACTCTCAGTATACGACATATCTAGACGCTGCTGACTGTATAT
CTCTATAGCTCAGGCATGCTAGTCTAGCGTCAGTAGTACTGACTGATGCGTCAGCAGCATACGTCT
GCGTCGCTGTGGTGTATACGTGCGACATACTCAGAGTCGCGCTCTCTACGGTGTCACTCGAGCTG
TAGATGCTCAGTGCGATACAGTACGCTCCATGCTGCTCATCAGCTGCTCAGCTCTGTATCTGCTCT
GACATGACGTGACTCAGCGCGAGCGTATGATCGAGATACTGATCGTACACTCACTGGTCAGTGTG
TAGCGTGACTGCGTGCAGCTCAGTCAGAGCATCACTCCTCGCCATCATGTCCGAGACTAGTGGTG
GTAGCTGCTGTGCGAGTGTATCTGCGTGCTGCGC

A represents adenine nucleotides paired with biotin-modified thymines, where the streptavidin-coated gold rotor bead can be attached;
**bolded** region represents the 16-nt Target 1 sequence (the TS sequence);
yellow highlighted region represents the TAM (the complementary sequence of 5′-TTGAT-3′);
cyan highlighted region represents the segment with multiple incorporated dUTP-digoxigenin (Roche);
green highlighted region represents the segment with multiple incorporated fluorescein-dUTPs (Roche).

**Table S23. AF3 query sequences for the WT Ymu1 ternary complex (related to Methods)**

| | Sequences |
|---|---|
| Protein | MLQHKAYEYRIYPDKKQETLIAKTIGSSRYVYNHFLELWNKEYEETGKGLTYYACSKL LTKLKRDPETVWLCEVDKFSLQNSLRNLSDAFSRFFKGQNEHPQFKSKKSPRQSYT |

| | |
|---|---|
| | TQYTNNNIAVSGNCLKLPKLGLVKFADSREMKGRILNATVRRKSSGKFFVSILCEEEI<br>CELPKTDSSVGIDLGIIDFAVMSDGSRHDNNHFTRQMEERLRREQRKLARRALAAEK<br>RGISLSEARNYQKQRRKVARLYEKVANQRKEYLNKLSTEIVKNHDIICIEDLNVKGMM<br>RNHKLAKSISDVSWTSLVSKLQYKASWYGKEVIRISRWFPSSQICSECGHKDRKKPL<br>HVREWTCPVCHAHHDRDVNAARNILAEGLRIRALTPGS |
| RNA (5′-3′) | CCGCACCCCCUGCGGGGGUAGCUUGGUAAACAAGAGAAACCUCUGCCGGCAA<br>AGAAAUAAGCCGGUAAGUAUGCUCUGUUCCCAAGAAUCUCGUGACUUUAGUCA<br>UGAGAGGGGGGAUCUUCUGGAUUGUU |
| DNA (5′-3′) | AACAATCCAGAAGAATCAATCA |
| DNA (5′-3′) | TGATTGAT |
